# Spaceflight causes strain dependent gene expression changes associated with lipid and extracellular matrix dysregulation in the mouse kidney in vivo

**DOI:** 10.1101/2024.03.13.584781

**Authors:** Rebecca H. Finch, Geraldine Vitry, Keith Siew, Stephen B. Walsh, Afshin Behesti, Gary Hardiman, Willian A. da Silveira

## Abstract

To explore new worlds we must ensure humans can survive and thrive in the space environment. Incidence of kidney stones in astronauts is a major risk factor associated with long term missions, caused by increased blood calcium levels due to bone demineralisation triggered by microgravity and space radiation. Transcriptomic changes have been observed in other tissues during spaceflight, including the kidney. We analysed kidney transcriptome patterns in two different strains of mice flown on the International Space Station, C57BL/6J and BALB/c. Here we show a link between spaceflight and transcriptome patterns associated with dysregulation of lipid and extracellular matrix metabolism and altered transforming growth factor-beta signalling. A stronger response was seen in C57BL/6J mice than BALB/c. Genetic differences in hyaluronan metabolism between strains may confer protection against extracellular matrix remodelling through downregulation of epithelial-mesenchymal transition. We intend for our findings to contribute to development of new countermeasures against kidney disease in astronauts and people here on Earth.

## 1. Introduction

Renal health risks are one of the key risk factors facing astronauts on long term missions due to increased incidence of kidney stones after missions^1^. Mitigation strategies such as protective shielding and personal medicine are currently in place to try to reduce the health risks, but are insufficient to fully protect against the damage caused by the space environment^1^.

Space radiation is one of the stressors of spaceflight and causes increased levels of reactive oxygen species, which leads to oxidative stress, inflammation, genetic and epigenetic alterations, and mitochondrial dysfunction^2,3^. The kidney is particularly vulnerable to the stresses of spaceflight due to its large number of mitochondria; playing an important part in regulation of oxidative stress in the body and being severely affected by oxidative stress itself. It is affected in a multifactorial way and disruption of its oxidative stress regulation functions act as a positive feedback mechanism for continuous kidney damage^2^.

Previous research has shown downregulated expression of nuclear oxidative phosphorylation genes and upregulated expression of mitochondrial oxidative phosphorylation genes in space in multiple tissues including the kidney; this switch is thought to be caused by reactive oxygen species causing damage to transcripts from nuclear DNA^3^. The kidney is particularly sensitive to oxidative stress due to its large number of mitochondria. Other effects that space has been observed to exert on the kidneys include changes to endothelial cells and the cytoskeleton, vascular senescence, altered fluid distribution and neurohormonal balance^2^.

Animals have been used as models to assess the ability of humans to survive in space for the last seventy years, and the first mouse was launched into space in 1950^4^. Previous studies have shown that different strains of mice have different reactions to kidney injury. C57BL/6 mice showed more profound inflammation, intrinsic injury responses and renal architecture disruption than BALB/c mice in response to reversible unilateral ureteral obstruction, a model of renal fibrosis^5^. The two strains also show different sensitivities to radiation, C75BL/6 are more radioresistant, while BALB/c are more radiosensitive^6,7^ and BALB/c have lower ratios of mitochondrial DNA to nuclear DNA which has been associated with sensitivity to mitochondrial calcium overload^7^.

C57BL/6 mice have also been shown to be more sensitive to the effects of streptozotocin-induced diabetes than BALB/c mice^8^. Induction of kidney stones experimentally in mice has been found to be significantly more difficult than in rats, leading to the hypothesis that mice have species-specific protective mechanisms against kidney stones. A model of calcium oxalate crystal deposition in the mouse kidney was induced by injection of glyoxylate but began to decrease 12 days after administration^9^. In contrast to the difficulties inducing kidney stones in mice, they are commonly used as subjects in unilateral ureteral obstruction which is a model for chronic kidney disease and renal fibrosis^10^ both of which are diseases correlated with ageing.

Spaceflight is known to promote accelerated ageing, and microgravity in particular has been used as a model for ageing due to the similarities in physiological changes observed in both, including insulin resistance, decreased protein breakdown after meals, decreased immune function and dysregulation of cytokine production^11^.

One of the changes associated with ageing in the kidney is nephrosclerosis, which consists of glomerulosclerosis, interstitial fibrosis, and arteriosclerosis^12^. The incidence of chronic kidney disease development is higher in patients with kidney stones and therefore kidney stones are a predictor and risk factor for chronic kidney disease^13,14^. Tissue injury initiates renal fibrosis, and this injury may be caused by kidney stones. The formation of kidney stones in the papilla involves activation of inflammatory cascades, leading to the accumulation of calcium, phosphate, and oxalate ions in the interstitial space, forming Randall’s plaque. The plaque can erode into the renal pelvis leading to loss of cells and exposing it to urine supersaturated with calcium phosphate, which both contribute to renal stone formation^15^. High levels of reactive oxygen species associated with the formation of kidney stones can also cause lipid peroxidation which can damage cell membranes^15^ which is another mechanism by which fibrotic tissue injury may occur.

Fibrosis is a pathological build-up of extracellular matrix caused by trauma or injury, for instance a kidney stone, and is driven by disruption of transforming growth factor-β (TGF-β) signalling. Increased levels of TGF-β and PAI1 cause an increased production and decreased degradation of the extracellular matrix which lead to fibrotic build-up of its components^16^. Previous studies have shown an association between dysregulation of lipid metabolism, a decline in kidney function and development of chronic kidney disease^17^.

In spaceflight, profibrotic markers have been observed in mouse lung tissue^18^, and lipid dysregulation has been observed in mouse liver tissue^19^. TGF-β has also been identified as an important regulator of response to spaceflight^20^.

Interestingly, we have reported in the past unexpected differences in the intensity of transcriptomic changes in tissues, including the kidney, in mice of the C57BL/6J and BALB/c lineages when exposed to spaceflight. BALB/c presented subtle alterations in gene expression whereas C57BL/6J presented major alterations^3^.

In this study, for the first time, we focus on renal transcriptome alterations in space and how they could be impacted by genetic background.

## 2. Methods

### 2.1. Subjects

Transcriptomic data related to kidney tissue obtained in the missions Rodent Research-1 (RR-1) version 3^21^, Rodent Research-3 (RR-3) version 4^22^, and Rodent Research-7 (RR-7) version 8^23^ were obtained from NASA’s GeneLab Platform (https://genelab.nasa.gov/) from dataset identifiers OSDR-102, OSDR-163 and OSDR-253, and samples were processed as detailed in^3^.

### 2.2. Differential Expression Analysis

R Studio (version 1.4.1717)^24^ and the package DESeq2 (version 1.32.0)^25^ were used to perform differential gene expression analysis by fitting a generalised linear model to each gene following a negative binomial distribution. Differentially expressed genes were identified by Flight to Ground Control comparison on C57BL/6J mice in the OSD-102 dataset with an adjusted p value of 0.1. The same method was used on the C57BL/6J and C3H/HeJ mice in the OSD-253 dataset, and separate analyses were carried out for the 25 and 75 day timepoints compared to their ground control counterparts. For OSD-163 the Basal group was used as a common control, and differential expression analysis was carried out between Spaceflight versus Basal, and Ground Control versus Basal groups. Genes differentially expressed in the Ground Control group were excluded to leave only genes differentially expressed in Spaceflight compared to Basal. An adjusted p value of 0.1 was used for the threshold.

Heatmaps were created using threshold adjusted p-value <= 0.1 and log2 fold change (>1/<−1) with addition of manual annotation of genes of interest. The R package gplots version 3.1.1^26^ was used.

### 2.3. Pathway Level Analysis

Overrepresentation analysis was done using WebGestalt^27^ for datasets OSD-102, OSD-163 and OSD-253 from the analysis performed using Deseq2. Venny^28^ was used to determine common and different pathways between strains.

Gene set enrichment analysis was performed using GSEA (version 4.1.0)^29,30^ using datasets OSD-102, OSD-163 and OSD-253 independent from the analysis performed using Deseq2.

EnrichmentMap (versions 3.3.2 and 3.3.3) and AutoAnnotate (versions 1.3.4 and 1.3.5) applications in Cytoscape (versions 3.8.2 and 3.9.0) were used to visualise enriched pathways using FDR q-value cut off value 0.1^31,32^.

### 2.4. Comparison between strains

The set of protein inactivating genetic differences between C57BL/6J and BALB/c was taken from Timmermans and collaborators, 2017^33^. WebGestalt biological processes gene ontology database^27^ was used to assess the effect of the genetic differences between the two strains of mice on their responses to spaceflight. Venny^28^ was used to determine genes with protein inactivating mutations between C57BL/6J and BALB/c, compared to differentially expressed genes (adjusted p-value <= 0.1) in the RR-1 and RR-3 datasets. R Studio (version 1.4.1717)^24^ and package gplots^26^ was used to generate supervised heatmaps of the genes identified by these two comparisons.

Heatmaps were constructed as described in 2.2.

## 3. Results

### 3.1. Expression of genes involved in lipid metabolism, extracellular matrix and TGF-β signalling are affected in the kidney by spaceflight

Differential expression analysis of kidney transcriptome data from C57BL/6J (RR-1 mission) and BALB/c mice (RR-3 mission) was carried out to determine the differences in their responses to spaceflight. Differential expression analysis of C57BL/6J was determined by comparison of Spaceflight versus Ground Control groups and found 638 differentially expressed genes (supplementary table 1). The same analysis identified zero differentially expressed genes in RR-3 and an alternative approach to isolate specific genes altered by spaceflight on BALB/c kidney was used, i.e., both Spaceflight and Ground control groups were compared to basal levels of gene expression at the beginning of the experiment and genes that were altered only by spaceflight were selected, a total of 671 genes (supplementary table 2). The gene signature of the differentially expressed genes in C57BL/6J had a stronger and clearer separation between the Spaceflight and Ground Control groups than in BALB/c (Supplementary Fig. 3).

By focusing in on a subset of genes exhibiting a twofold increase or decrease in expression following exposure to spaceflight, we pinpointed alterations in genes involved in lipid metabolic pathways, extracellular matrix degradation, and TGF-β signalling within the kidneys of both lineages (Fig. 1). Interestingly, the *Ccl28* gene – belonging to the TGF-β signalling pathway - was the most differentially expressed gene in the C57BL/6J samples with a log2 fold change of 2.05 (adjusted p-value <= 0.1) (Fig. 1a, supplementary table 1).

**Fig. 1.**
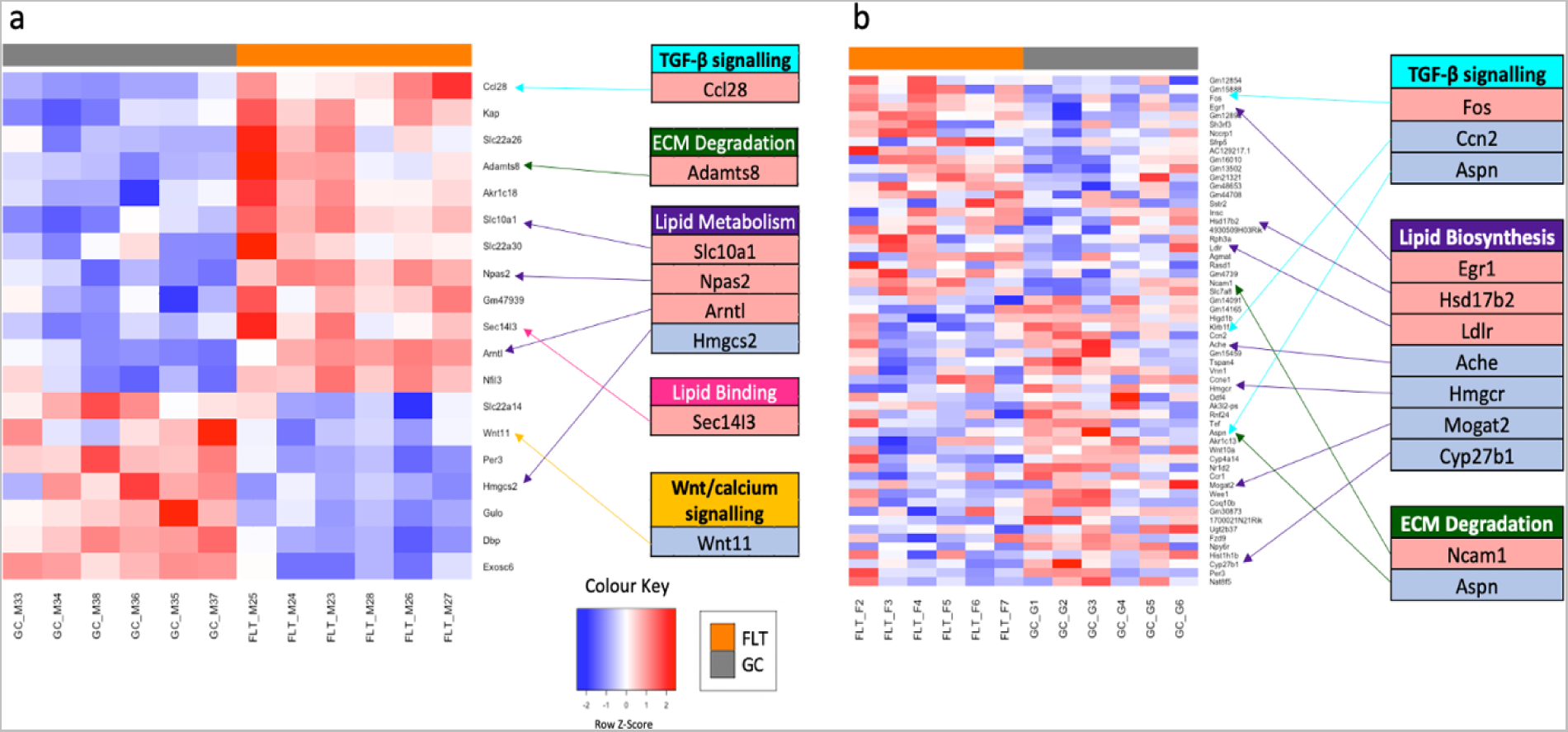
Gene expression signature of mouse kidneys exposed to spaceflight shows genes of TGF-β and Lipid Metabolism altered in C57BL/6J and BALB/c mice lineages. Differentially expressed genes in spaceflight in kidney tissue from **a)** the C57BL/6J mice on the RR-1 mission and **b)** the BALB/c mice on the RR-3 mission (adjusted p-value <= 0.1) with a minimum of twice the expected log2 fold change (>1/<−1). FLT = Spaceflight samples, GC = Ground Control Samples.

Genes involved in lipid metabolism were differentially expressed in C57BL/6J. *Slc10a1*, *Npas2* and *Arntl*, were upregulated, while *Hmgcs2* was downregulated with a log2 fold change of −1.68 (Fig. 1).

Genes involved in lipid metabolism were also differentially expressed in BALB/c, including *Egr1* which was positively differentially expressed (log2 fold change 1.59), and *Hmgcr* which was negatively differentially expressed (log2 fold change −1.13).

Both strains exhibited differential expression in genes involved in extracellular matrix degradation. *Adamts8* was upregulated in C57BL/6J, while expression of *Ncam1* and *Aspn* were altered in BALB/c (Fig. 1).

Enrichment of pathways related to TGF-β signalling was seen in genes upregulated in BALB/c (RR-3) and downregulated in C57BL/6J (RR-1) in response to spaceflight (Fig. 1). As previously mentioned, both strains showed increased cholesterol biosynthesis, and BALB/c exhibited enrichment in hallmark Myc targets, both of which affect TFG-β signalling. Both strains exhibited differential expression in genes involved in TGF-β signalling. This included *Ccl28,* the most differentially expressed gene in C57BL/6J (log2 fold change 2.05). In BALB/c *Fos* was upregulated (log2 fold change 1.60), while *Ccn2* and *Aspn* were downregulated (Fig. 1).

In C57BL/6J mice, the *Wnt11* gene shows downregulation in the spaceflight group compared to the control group, with a log2 fold change of −1.15 (adjusted p-value <= 0.1). This gene is involved in Wnt signalling, a pathway which is involved in crosstalk with TGF-beta signalling.

### 3.2. Overrepresentation analysis shows spaceflight is associated with positive enrichment of cholesterol metabolic pathways and negative enrichment of ECM pathways in the kidney

Overrepresentation analysis of both datasets showed upregulation in genes linked with cholesterol related pathways.

Genes connected with extracellular matrix and TGF-β signalling pathways were upregulated in BALB/c (RR-3) and downregulated in C57BL/6J (RR-1) (Table 1, supplementary figure 2). A GSEA analysis carried out using the Gene Ontology biological process database, supplementary table 5, showed that pathways related to increased lipid and fat metabolism were enriched in both datasets. In BALB/c, there was an enrichment in several hallmark pathways connected with dysregulated extracellular matrix metabolism, including Myc targets, adipogenesis and epithelial-mesenchymal transition (Supplementary Table 4).

**Table 1.** Spaceflight exposed kidney shows cholesterol and protein synthesis pathways positively and negatively enriched, respectively, in both lineages, while extracellular matrix pathways show negative enrichment in C57BL/6J and positive enrichment in BALB/c mice. Biological process pathways enriched by overrepresentation of differentially expressed genes in kidney transcriptomic data from C57BL/6J (RR-1) and BALB/c (RR-3) mice (FDR < 0.05).

|  | Upregulated in C57BL/6J | Downregulated in C57BL/6J |
| --- | --- | --- |
| BALB/c up | Cholesterol Biosynthesis Pathway<br>Cholesterol biosynthetic<br>Cholesterol biosynthesis<br>Superpathway of cholesterol biosynthesis<br>Cholesterol Biosynthesis with Skeletal Dysplasias<br>Cholesterol metabolism (includes both Bloch and Kandutsch-Russell pathways)<br>Steroid biosynthetic<br>Regulation of cholesterol biosynthesis by SREBP (SREBF)<br>Cholesterol biosynthesis, squalene 2,3-epoxide => cholesterol<br>Cholesterol biosynthesis II (via 24,25-dihydrolanosterol) | Genes encoding secreted soluble factors<br>Ensemble of genes encoding ECM-associated proteins including ECM-affiliated proteins, ECM regulators and secreted factors<br>Kisspeptin/Kisspeptin Receptor System in the Ovary<br>E3 ubiquitin ligases ubiquitinate target proteins<br>Ovarian steroidogenesis<br>Protein ubiquitination<br>Ensemble of genes encoding extracellular matrix and extracellular matrix-associated proteins<br>TGF-beta signaling pathway |
| BALB/c down | N/A | Peptide chain elongation<br>Eukaryotic Translation Elongation metabolism<br>Ribosome<br>Cytoplasmic Ribosomal Proteins<br>Viral mRNA Translation<br>Selenocysteine synthesis<br>Translation<br>Eukaryotic Translation Termination<br>Nonsense Mediated Decay (NMD) independent of the Exon Junction Complex (EJC) |

### 3.3. Diverging patterns of lipid synthesis, protein synthesis and circadian rhythm in response to spaceflight in C57BL/6J and BALB/c mice determined by overrepresentation analysis

Overrepresentation analysis on both C57BL/6J (RR-1) and BALB/c (RR-3) using the hallmarks database indicated increased enrichment of pathways associated with epithelial cell remodelling, but different pathways were identified in each dataset (supplementary table 3, supplementary table 4).

Comparison of statistically relevant GSEA results of biological processes for C57BL/6J (RR-1) and BALB/c (RR-3) datasets identified clusters of gene sets, including clusters with opposite patterns between datasets (Fig. 2). Notably, cholesterol biosynthesis and fatty acyl coA biosynthesis were positively correlated with spaceflight in C57BL/6J and negatively correlated in BALB/c. The opposite pattern was seen in hallmarks related to translation, protein folding and circadian rhythm. Lipid metabolism and storage, cell cycle processes and immune response were positively correlated with spaceflight in both C57BL/6J and BALB/c (Supplementary Fig. 3).

**Fig. 2.**
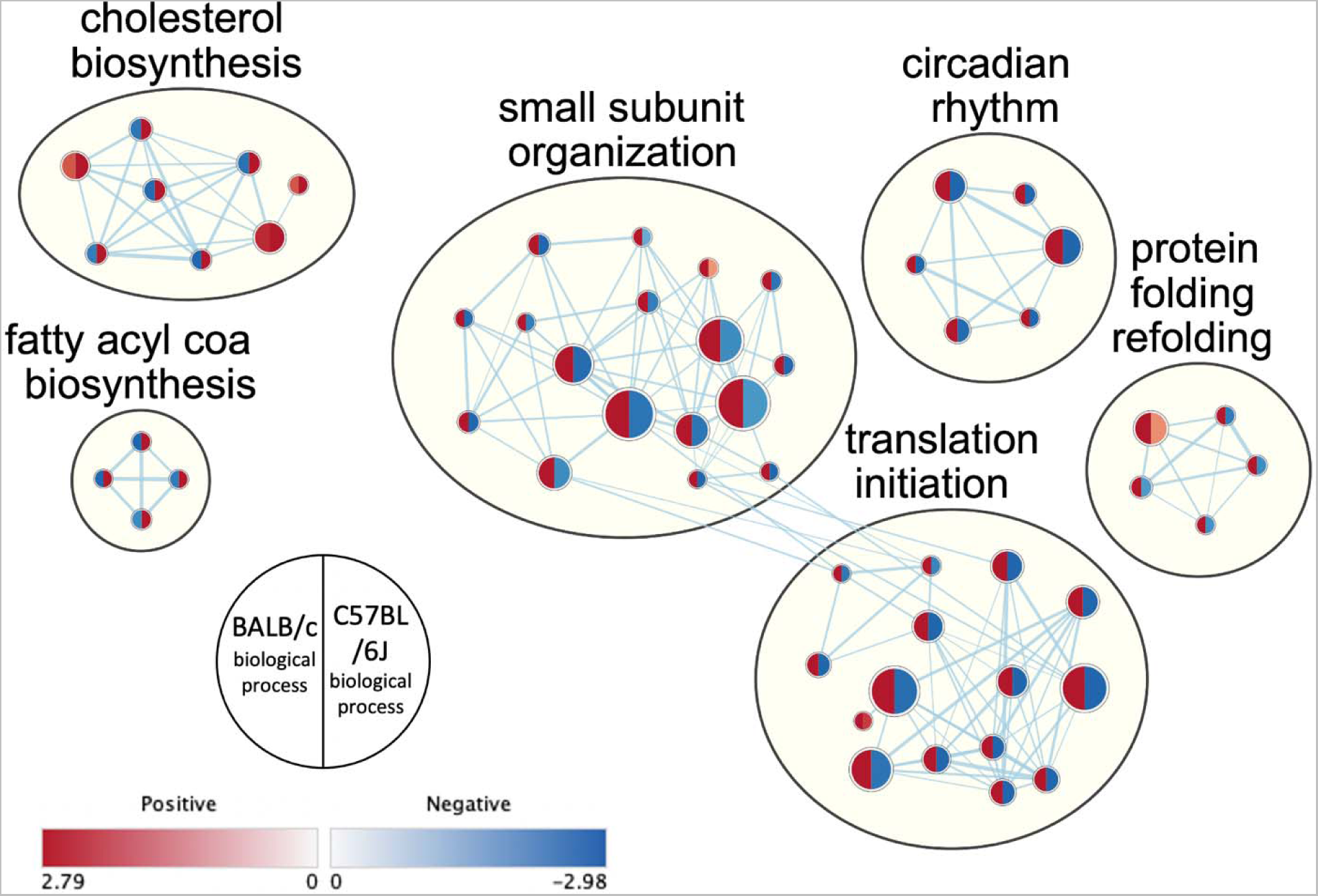
Network analysis showing clusters with opposite expression patterns in spaceflight in gene ontology biological process comparison between kidney tissue data from C57BL/6J (RR-1) and BALB/c (RR-3) datasets.

### 3.4. C57BL/6J and BALB/c mice present genetic background differences in lipid and extracellular matrix metabolism reflected in the transcriptomics difference in spaceflight

While mice in both missions were affected by the space environment, the C57BL/6J mice appeared to experience more severe effects than the BALB/c mice. The BALB/c mouse strain has previously been found to have a distinct set of genes with protein inactivating mutations^33^ compared to the C57BL/6J mouse strain. These genetic differences were assessed to provide insight into their different reactions to spaceflight, (Fig. 3, supplementary Fig. 4, and supplementary table 6). The hyaluronan metabolic pathway had the highest enrichment ratio, and genes in this pathway were differentially expressed by BALB/c mice in spaceflight. Other pathways containing protein inactivating mutations between the two strains which contained genes differentially expressed in BALB/c in spaceflight related to cytoskeleton and the innate immune system. Pathways containing protein inactivating mutations between the two strains which contained genes differentially expressed in C57BL/6J in spaceflight were connected to organic acid metabolism and negative regulation of intracellular signalling.

**Fig. 3.**
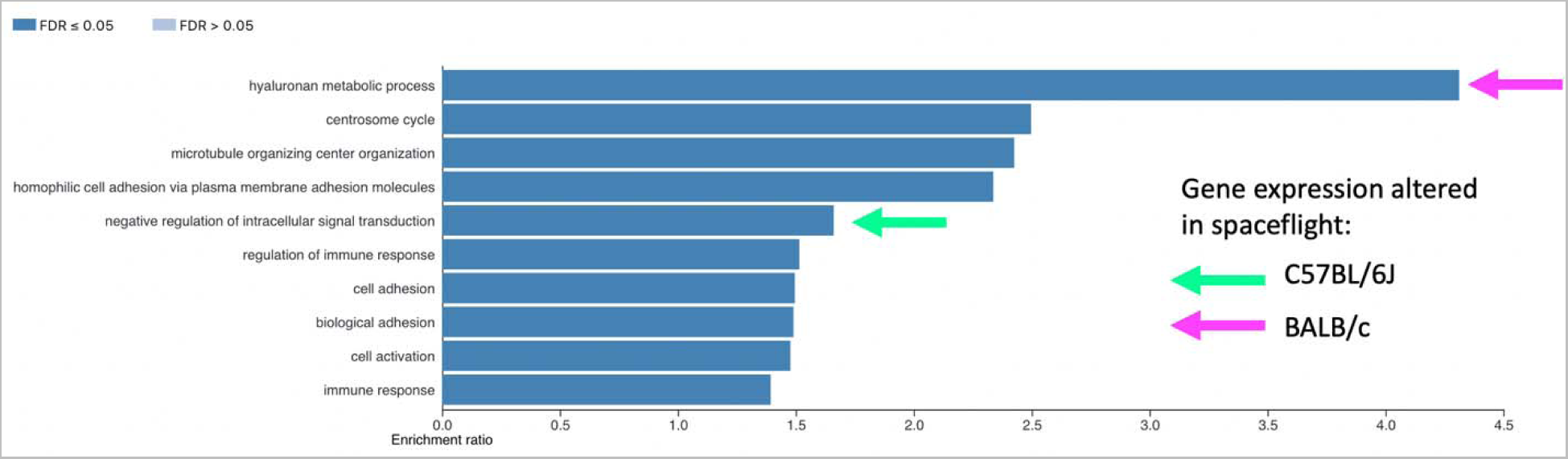
Enriched biological process pathways of protein inactivating mutations in BALB/c mice compared to C57BL/6J strain reveal connections with expression of genes involved in pathways related to hyaluronan metabolism in BALB/c and intracellular signalling in C57BL/6J. Top 10 enriched biological process pathways in the genetics of protein inactivating mutations in BALB/c strain of mice compared to C57BL/6J strain^33^ and their connections with enriched pathways in the transcriptomic data of kidneys obtained from C57BL/6J mice (RR-1) (green arrow) and BALB/c (RR-3) (pink arrow).

Comparison between genes with protein inactivating mutations in BALB/c compared to C57BL/6J and differentially expressed genes in C57BL/6J (RR-1) and BALB/c (RR-3) (Fig. 4) showed upregulation of genes related to fatty acid metabolism and downregulation of genes related to cell junction, and expression of genes related to cholesterol metabolism and PPAR signalling were altered in the comparison with differentially expressed genes in C57BL/6J. In comparison with differentially expressed genes in BALB/c, regulation of genes related to lipid transport were identified, as well as altered expression of genes involved in Wnt-protein binding, gap junction and lipid metabolism. In general, a pattern of upregulation of genes involved in lipid processes was seen in C57BL/6J and downregulation of genes involved in lipid processes was seen in BALB/c in response to spaceflight.

**Fig. 4.**
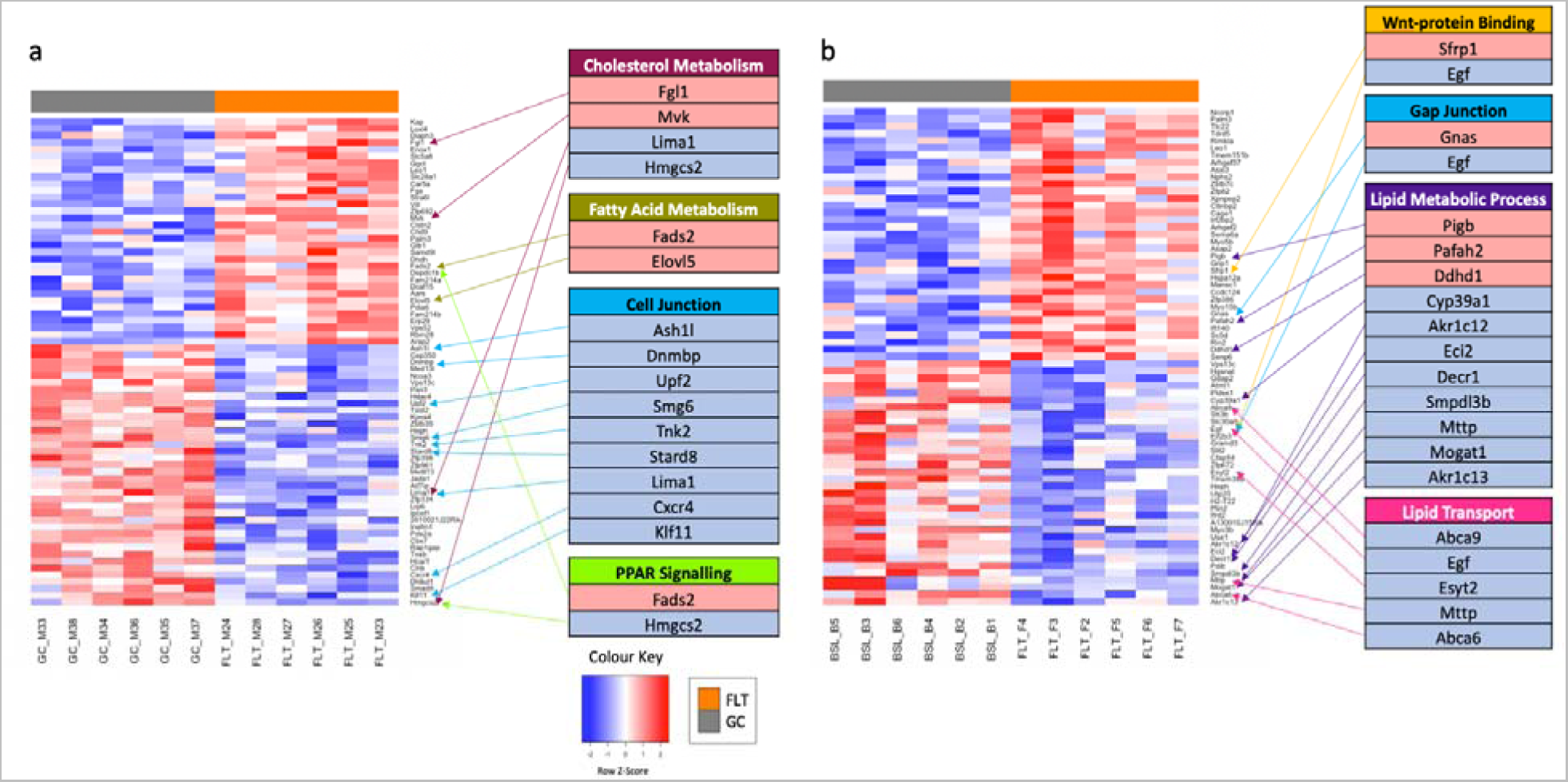
Gene expression signatures of mouse kidneys exposed to spaceflight limited by protein inactivating mutations between the BALB/c and C57BL/6J genes show genes involved in lipid metabolism and cell junction altered in C57BL/6J and BALB/c mouse lineages. Differentially expressed genes in kidney transcriptomic data from **a)** C57BL/6J mice on the RR-1 mission and **b)** BALB/c mice on the RR-3 mission (adjusted p-value <= 0.1). FLT = Spaceflight samples, GC = Ground Control Samples.

## 4. Discussion

With progressions in the space era and the advent of space tourism^34^, it will be increasingly common for larger numbers of people with different genetic backgrounds to have access to space. Taking into account the limited amount of people who have been exposed to space so far, it is unknown how the genetic backgrounds of people could affect their responses to the space environment and how they may differ from the responses to stress on Earth^35^ so it is of crucial importance for us to explore this area. In this work for the first time we show that not only do the kidneys of BALB/c and C57BL/6J mice have distinct responses to space flight but these can be linked back to their genetic differences.

Mice in both the C57BL/6J and BALB/c lineages – RR-1 and RR-3 missions, respectively - showed transcriptomic alterations associated with their exposure to space, however their responses were different. Different sets of genes were differentially expressed in each lineage, with extracellular matrix metabolism and TGF-beta signalling pathways upregulated and downregulated in BALB/c and C57BL/6J respectively.

Common alterations in pathways including lipid metabolism, extracellular matrix degradation and TGF-β signalling were found. Disruption of lipid metabolism can impair extracellular matrix degradation, via increased levels of TGF-β and PAI1, leading to a build-up of extracellular matrix and promoting fibrosis^16^. The strong expression of the *Ccl28* gene suggests an inflammatory environment in the C57BL/6J spaceflight exposed kidney as its expression is mediated by proinflammatory cytokines, and causes regulatory T cells to migrate to mucosal surfaces and increase TGF-β production^36^.

Alterations in expression of genes and enrichment of pathways connected with lipid metabolism were seen in both C57BL/6J (RR-1) and BALB/c (RR-3) mice when exposed to spaceflight. Lipid storage and fatty acid metabolism were enriched in network analysis of biological processes for both C57BL/6J and BALB/c (Supplementary Figure 3), and GSEA analysis showed enrichment of pathways related to lipid metabolism and synthesis in genes upregulated in both datasets (Table 1). This could indicate increased lipid accumulation which can cause impairment of extracellular matrix degradation^16^.

Cholesterol biosynthesis and fatty acyl co-A biosynthesis were positively correlated with spaceflight in C57BL/6J, with the opposite pattern seen in BALB/c (Fig. 2), potentially affected by the *Hmgcr* gene which plays a role in cholesterol biosynthesis^37^ which we have shown is downregulated in spaceflight exposed BALB/c (Fig. 1). Accumulation of cholesterol increases TGF-β levels and disrupts extracellular matrix degradation through inhibition of PAI1, which controls normal degradation of the extracellular matrix^38^. Decreased lipid synthesis may be an adaptive response against contribution to fibrotic damage in the BALB/c mice or may be indicative that they are less affected by oxidative stress.

Maintaining homeostasis in lipid metabolism is important to maintain normal kidney function, which is essential for blood pressure and cardiovascular system regulation. Disrupted lipid metabolism can lead to chronic kidney disease and atherosclerosis, which can in turn lead to cardiovascular disease, the leading cause of death in chronic kidney disease^17^. Interestingly, BALB/c appears to exhibit an adaptive response to lipid dysfunction in spaceflight, whereas C57BL/6J does not.

Statistically significantly differentially expressed genes in BALB/c indicate that the strain is affected by spaceflight, but exhibits an adaptive response, protecting it from lipid dysfunction. Upregulation of *Egr1* in BALB/c (Fig. 1) indicates that the strain is affected by spaceflight, as *Egr1* has been shown to play a role in lipid metabolism^39^, and also in an increase of extracellular matrix components and epithelial-mesenchymal transition, both profibrotic processes which can contribute to kidney disease^40^.

The negative differential expression of *Hmgcr* in BALB/c (Fig. 1) could represent an adaptive response. *Hmgcr* encodes HMG-CoA reductase which is an enzyme involved in the ketogenesis pathway as is *Hmgcs2* which was negatively differentially expressed in C57BL/6J. HMG-CoA reductase is also involved in the mevalonate pathway which synthesises cholesterol, catalysing the conversion of HMG-CoA to mevalonate^37^. HMG-CoA reductase inhibitors are used to reduce the risk of death from cardiovascular disease in chronic kidney disease patients, as the inhibition of mevalonate production reduces total cholesterol and low-density lipoprotein cholesterol^41^. Therefore, downregulation of *Hmgcr* in BALB/c may indicate increased lipid metabolism as shown in a previous study^19^, which could act as a protective mechanism against kidney injury during spaceflight. Together these results suggest that BALB/c mice exhibit an adaptive response to lipid dysfunction in spaceflight.

C57BL/6J however did not exhibit an adaptive response to the stresses of spaceflight, and gene expression indicates activated mechanisms related to increased inflammation. The *Hmgcs2* gene, downregulated in C57BL/6J (Fig. 1), encodes the protein 3-hydroxy-3-methylglutaryl-CoA (HMG-CoA synthase) which catalyses the conversion of acetyl-CoA and acetoacetyl-CoA to HMG-CoA and CoA, a which is a rate-limiting step in ketogenesis^42^. Impairment of ketogenesis causes hepatic injury and inflammation^43^ and downregulation of HMG-CoA synthase has been identified as a marker for kidney stone disease in a model of induced urolithiasis in rats^44^; suggesting that the downregulation of HMG-CoA synthase in our data may be a marker for kidney injury.

Although physiological examination was not carried out as part of this study to confirm signs of lipid dysfunction in mouse kidneys exposed to spaceflight, previous studies have found both transcriptomic and physiological changes associated with this condition in other organs. Abnormal lipid accumulation was detected in liver tissue from mice exposed to spaceflight by Oil Red O staining, supporting the transcriptomic data on lipid dysfunction pathways in liver^19^. In another study, increased numbers of lipid droplets were also observed in the liver of spaceflight mice by CARS microscopy and Oil Red O staining along with upregulation of triglyceride biosynthetic pathways seen in multi-omics analysis^45^.

Maintaining normal levels of turnover in the extracellular matrix is important for normal kidney function, as a fibrotic build-up of its components can cause scarring and impair function, leading to kidney disease^46^.

BALB/c exposed to spaceflight showed enrichment in pathways connected with dysregulated extracellular matrix metabolism (Supplementary Table 4) including Myc targets, adipogenesis and epithelial-mesenchymal transition. In the kidney, gene targets of the Myc group of proteins have been linked to an activation of glycolytic metabolism which increases production and deposition of extracellular matrix^47^ and promotion of TGF-β signalling via transcriptional activation of integrin αv^48^. Adipogenesis and disruption of fatty acid metabolism can impair extracellular matrix degradation via intracellular accumulation of lipids and lipotoxicity^49^.

Mesenchymal cells such as fibroblasts produce the components of the extracellular matrix, so increased levels due to epithelial-mesenchymal transition can cause a potentially fibrotic build-up of these components^50^.

Lipid accumulation - signs of which were observed in both strains of mice - can also lead to the impairment of normal extracellular matrix degradation. A build up in extracellular matrix components can lead to kidney disease as disrupted wound repair mechanisms are unable to restore kidney function^51^.

Genes in pathways related to TGF-β signalling were upregulated in BALB/c and downregulated in C57BL/6J (Fig. 1). TGF-β is a cytokine which increases the production of extracellular matrix components and can lead to kidney dysfunction via glomerulosclerosis and tubulointerstitial fibrosis, leading to renal dysfunction^52^. TGF-β has been previously identified as a master regulator of both response to spaceflight through microRNA signatures^20^ and of fibrosis^53^.

Downregulation of the *Wnt11* gene was associated with spaceflight in C57BL/6J. Its expression is essential for normal development of the glomeruli, and for uretic epithelial branching in the kidney; and knockout of the gene has been shown to be lethal to mice *in utero*^54^. *Wnt11* deficiency in older mice has been found to result in tubular abnormalities, glomerular cysts, and interstitial fibrosis^55^. *Wnt11* is involved in Wnt/calcium signalling and dysregulation of this pathway has been associated with cellular senescence and diseases related to ageing, including renal fibrosis^56^. Wnt signalling is involved in crosstalk with other profibrotic signalling pathways including the renin-angiotensin system, TGF-β, Notch and Hedgehog^56^. Inhibitors of Wnt signalling have been explored as treatments for fibrosis but have resulted in off-target effects^56^ perhaps in part due to the profibrotic effects of *Wnt11* downregulation.

In C57BL/6J there was enrichment in TNF-alpha signalling (Supplementary Table 3), which causes intense temporary inflammation and fibrosis^57^; and mTORC signalling (Supplementary Table 3) which increases fibroblast activation and interstitial fibrosis^58^. Interferon alpha response, interferon gamma response and JAK-STAT signalling indicate increase in adaptive immune response, which may be in response to increased inflammation caused by oxidative stress. Also enriched was angiogenesis (Supplementary Table 3), the formation of new blood vessels as a part of wound healing^59^, and overactivation of the wound healing process is a profibrotic process which can lead to kidney injury.

Both C57BL/6J and BALB/c showed enrichment of the hallmark E2F targets (Supplementary Table 3, Supplementary Table 4), which are involved in DNA repair, and BALB/c also shows enrichment of the DNA repair pathway (Supplementary Table 4). The DNA damage response has been found to contribute to the progression of fibrosis in systemic sclerosis^60^. The JAK-STAT signalling pathway is also enriched in BALB/c, indicating increased inflammation in both lineages.

Clusters of biological processes related to cell cycle and immune response (Supplementary Figure 3) were positively correlated with spaceflight in both datasets, which could represent responses to cellular damage by oxidative stress. Circadian rhythm was affected, negatively correlated with spaceflight in C57BL/6J and positively in BALB/c (Supplementary Figure 3). Mitochondrial gene expression and function are affected by the circadian clock^61^, which has also been seen to be dysregulated in astronauts^1^. Protein re-folding pathways were negatively correlated with spaceflight in C57BL/6J and positively in BALB/c (Supplementary Figure 3). Oxidative stress can lead to protein unfolding, and if unfolded proteins are not re-folded or destroyed, they can accumulate and cause loss of proteostasis, which is associated with ageing^62^. Ribosome activity and translation initiation were also negatively correlated with spaceflight in C57BL/6J and positively in BALB/c (Supplementary Figure 3), which has been observed previously to be disrupted in spaceflight studies and is linked to loss of proteostasis ^3^.

Photoreceptor development and light perception were correlated negatively with spaceflight in both datasets (Supplementary Figure 3); many genetic diseases affect both the kidney and the retina due to the presence of common developmental pathways^63^ so these enrichment patterns likely represent expression of genes common to both the retina and the kidney with different functions in each.

To seek explanations for the difference in responses to spaceflight from C57BL/6J and BALB/c mice, we determined enriched pathways for the protein inactivating mutations found previously between these lineages^33^ and hyaluronan metabolism was the pathway found to be most enriched (Fig. 3). Hyaluronan is a component of the extracellular matrix which plays important roles in wound healing and inflammation and is upregulated during these processes^64^.

No genes involved in the metabolism of hyaluronan were differentially expressed in C57BL/6J, but one gene involved in hyaluronan metabolism, *Egf,* was downregulated in BALB/c (Fig. 3). EGF treatment in rat mesothelial tissue was shown to result in the increase of hyaluronan synthesis and epithelial-mesenchymal transition^65^. The hyaluronan synthesis process is linked with morphological changes such as mitosis and anchorage-independent growth seen in epithelial-mesenchymal transition, by inducing de-adhesion and budding of extracellular vesicles^65^. Increased EGF protein in the urine has been identified as a hallmark of renal interstitial fibrosis, and is correlated with increased transcription of the *Egf* gene^66^. Downregulation of *Egf* in the BALB/c mice could potentially confer resistance to extracellular matrix remodelling via reduced hyaluronan synthesis.

Previous studies have shown similar, milder effects on BALB/c mice compared to C57BL/6J in response to injury and stress. A study on retinal ischemia/reperfusion injury, a model for diabetic retinopathy, caused increased inflammation and damage due to oxidative stress in C57BL/6J compared to BALB/c mice^67^. BALB/c has also been shown to have milder reactions to UVB exposure than C57BL/6J^68^, despite their lack of protective pigmentation. In this study, higher hyaluronan levels and lower collagen levels were measured in C57BL/6J mice. Hyaluronan is a component of the extracellular matrix, plays an important role in the wound repair process, and has been shown to contribute to the development of fibrosis^69^. Increased metabolism of hyaluronan showed the highest enrichment in a functional enrichment analysis of the genetic differences in BALB/c compared to C57BL/6J mice.

Genes with protein inactivating mutations between the strains were also differentially expressed in BALB/c (Supplementary Table 2, Supplementary Table 6) including genes with links to lipid transport which were downregulated, and genes linked to lipid metabolism, gap junction and Wnt-protein binding were also affected. For example, *Mogat1*, which is a lipid precursor which usually has high levels of expression in the kidney^70^ is downregulated in BALB/c during spaceflight.

In C57BL/6J, genes with protein inactivating mutations between the strains which were differentially expressed (Supplementary Table 1, Supplementary Table 6) including alterations in expression of genes involved in cholesterol metabolism and PPAR signalling, upregulation of genes related to fatty acid metabolism, and downregulation of genes related to cell junction. *Fads* and *Elovl5* are both involved in fatty acid synthesis, potentially contributing to impairment of extracellular matrix via increasing build-up of lipids and lipotoxicity^49^. These genes are not differentially expressed in BALB/c (RR-3), so the differences in these genes between the two mouse strains may confer some resistance to extracellular matrix remodelling. Variation in the *Hmgcs2* gene in BALB/c compared to C57BL/6J may also confer some fibrotic resistance to BALB/c as its downregulation as seen in C57BL/6J (RR-1) promotes proinflammatory ketogenesis^42,43^. Genes related to cell junction were downregulated, potentially contributing to de-adhesion which is a part of epithelial-mesenchymal transition.

We would expect that mice exposed to the same length of spaceflight would have similar responses to the stress, nonetheless we encountered a very different response between strains. Their different genetic background is potentially a factor for explaining this difference.

While the C57BL/6J (RR-1) and BALB/c (RR-3) mice showed pro-fibrotic hallmarks they did not develop kidney stones, as seen in astronauts. Nonetheless, a recent work performed a multi-omics analysis of mice exposed to spaceflight on different mission or to ionizing radiation and found a signature of renal fibrosis and increased risk of kidney stone formation^71^. Mice on Earth have previously proven to be poor experimental models of kidney stones^9^ but good models of renal fibrosis^10^ and this research indicates that the same may be true in space. Kidney stones and fibrosis can both result from metabolic disturbances, which have been seen both in this data and previous studies. Better understanding of the metabolic effects of spaceflight could lead to the development of new treatments and protective measures, and to inform risk assessments for astronauts.

Countermeasures against the increased risk of kidney stones were first suggested by^72^ including a physical exercise regimen and increased fluid intake for astronauts. However, the current mitigations in place are not sufficient to bring the risk of kidney issues on long term space missions to within acceptable levels^1^. In the present study we show that gene expression related to extracellular matrix dysregulation and fibrosis in the kidney are upregulated during spaceflight, which are two factors known to promote kidney stone formation. As such, considering treatments for fibrosis may be of relevance in the future as a countermeasure.

In conclusion, kidney stones are considered the primary risk to the renal health of astronauts, but because of the link between the development of kidney stones and renal fibrosis, and observations of profibrotic markers in this and other studies, fibrosis should be considered as an additional serious potential risk on long term space missions.

Further study into the influences of genetic background on the response to spaceflight should be conducted to discover potential protective genes.

## Author Contributions

RHF and G.V. analysed data. RHF wrote the manuscript and prepared the figures. KS and SBW contributed to the writing and editing of the manuscript, AB participated in the exchange of ideas and edited the Manuscript and gave feedback on the draft. GH designed the study and edited the manuscript. WAdS funding acquisition, designed the study, supervised the project, wrote and edited the manuscript.

## Acknowledgments

WAdS acknowledges this work was partially funded by the ESA grant/contract 4000131202/20/NL/PG/pt ‘‘Space Omics: Toward an integrated ESA/NASA –omics database for spaceflight and ground facilities experiments’’. SBW acknowledges this work was partially funded by the UK Space Agency through grant [ST/X000036/1] administered by the Science and Technology Facilities Council (STFC). KS acknowledges this research was funded in part by the Wellcome Trust [Grant number 110282/Z/15/Z]. For the purpose of open access, the author has applied a CC BY public copyright licence to any Author Accepted Manuscript version arising from this submission. “This work was allowed by the free access online repository data resources NASA genelab. The Rodent Research 1 The Rodent Research 3 and data collection is supervised by Jonathan Galazka, Project Scientist, NASA GeneLab and Ruth Globus, RR-1 Project Scientist NASA ARC. GH Acknowledges support from NIH U54MD010706, U01DA045300 and QUB start-up funds.

## Data availability

The datasets used in this work are publicly available at the database NASA Genelab (https://genelab.nasa.gov/) under the classification: OSD-102, OSD-163 and OSD-253.

