## Supplementary Figures for "Spaceflight causes strain dependent gene expression changes associated with lipid and extracellular matrix dysregulation in the mouse kidney in vivo"

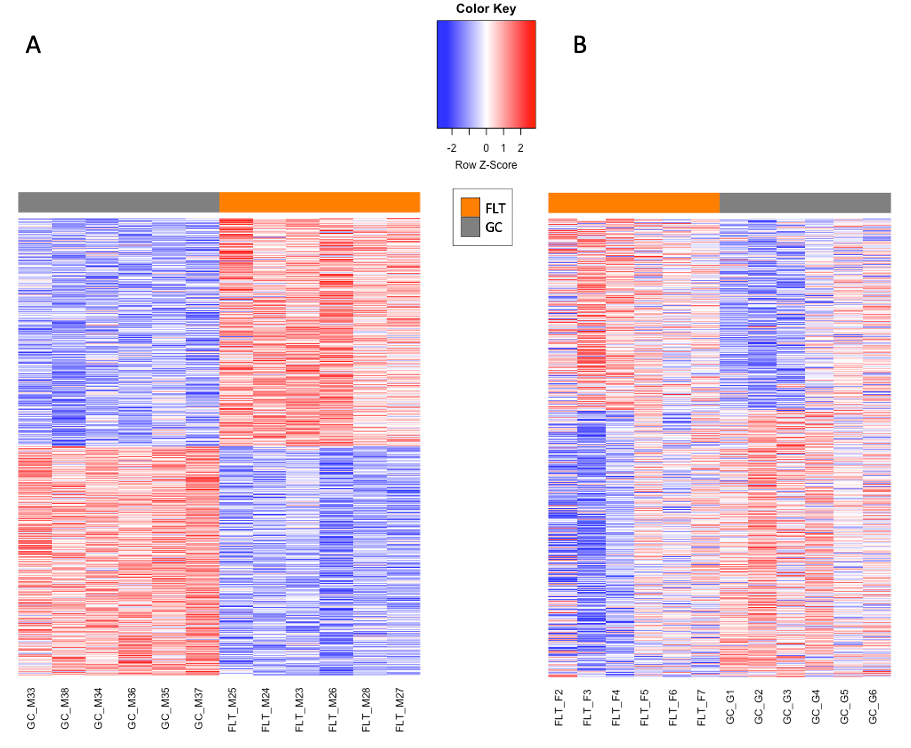

**Supplementary Figure 1** Heatmaps of genes differentially expressed in spaceflight in kidney tissue from a) C57BL/6J (RR-1) and b) BALB/c (RR-3) (adjusted p-value <= 0.1). FLT = spaceflight group , GC = ground control group.

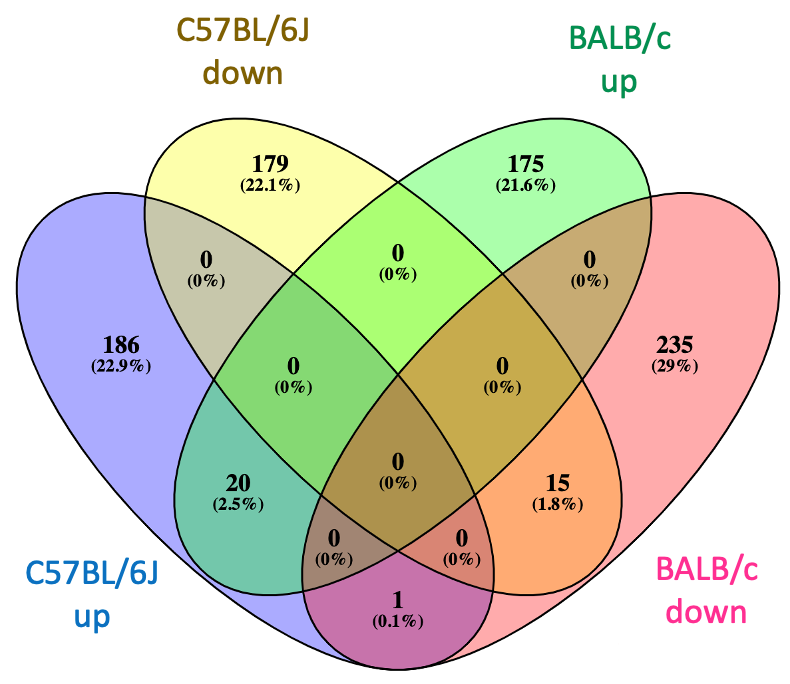

**Supplementary Figure 2**  Genes significantly upregulated and downregulated in differential expression determined by DESeq2 analysis on the effect of spaceflight on transcriptomic expression in C57BL/6J (RR-1) and BALB/c (RR-3) mice kidneys.

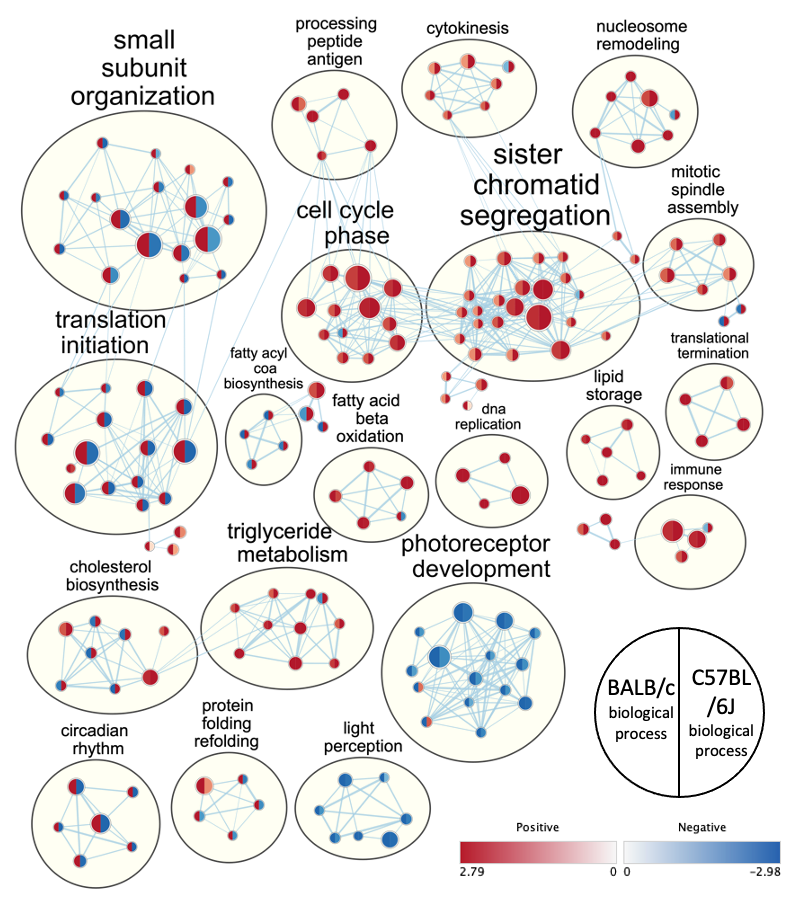

**Supplementary Figure 3** Network analysis showing top 20 clusters in gene ontology biological processes between C57BL/6J and BALB/c.

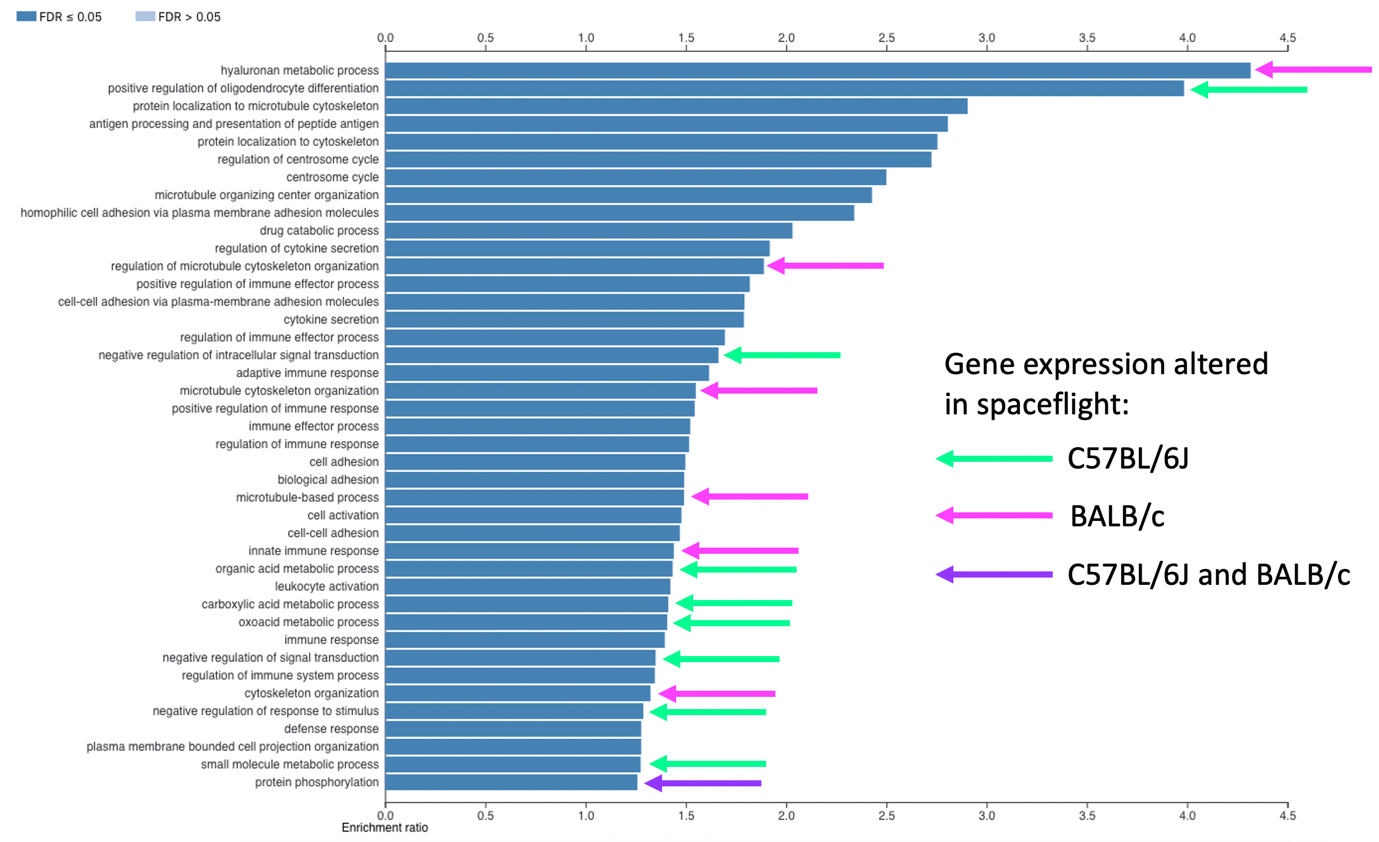

**Supplementary Figure 4 Enriched biological process pathways of non-synonymous mutations in BALB/c mice compared to C57BL/6J strain reveal connections with expression of genes involved in pathways related to hyaluronan metabolism in BALB/c and intracellular signalling in C57BL/6J.** Enriched biological process pathways in the genetics of non-synonymous mutations in BALB/c strain of mice compared to C57BL/6J strain (Timmermans, Van Montagu and Libert, 2017) and their connections with enriched pathways in the transcriptomic data of kidneys obtained from C57BL/6J mice (RR-1) and BALB/c (RR-3).

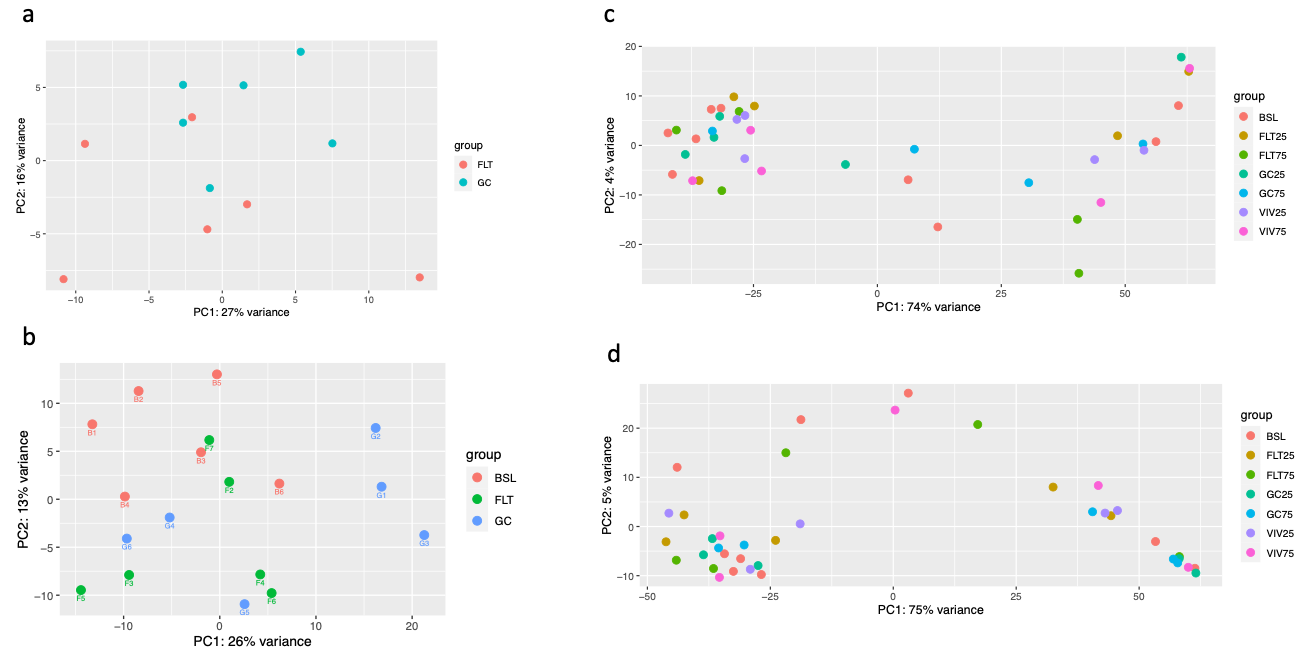

**Supplementary Figure 5** Principal component analysis (PCA) plots of differential expression analysis performed on experimental and control groups of kidney transcriptomic data from **a)** RR-1 mission involving C57BL/6J mice **b)** RR-3 mission involving BALB/c mice **c)** RR-7 mission involving C57BL/6J mice and **d)** RR-7 mission involving C3H/HeJ mice. FLT = spaceflight group, GC = ground control, BSL = basal control, FLT25 = spaceflight group 25 days in space, FLT75 = spaceflight group 75 days in space, GC = ground control group 25 days, GC75 = ground control group 75 days, VIV25 = vivarium control 25 days, VIV75 = vivarium control 75

*Lack of differences in gene expression patterns in spaceflight and ground control groups of RR-7 mission indicate need to standardise protocols for rodent research on the ISS*

Similar analyses were attempted using data from C57BL/6J and C3H/HeJ mice flown on the RR-7 mission, however transcriptomic analysis revealed few differences between the spaceflight and ground control groups. For the C57BL/6J mice in RR-7, zero differentially expressed genes were identified between the 25 and 75 day spaceflight groups each compared with their corresponding ground control groups. The same comparisons carried out C3H/HeJ mouse groups identified one differentially expressed gene at each timepoint. Likewise, functional enrichment analysis of hallmarks resulted in only one significantly enriched gene set (FDR < 0.25) at 25 days and zero at 75 days for C57BL/6J, and 3 gene sets each at 25 days and 75 days for C3H/HeJ.

Comparison of principal component analyses revealed a less clear pattern of separation between groups of both the C57BL/6J and C3H/HeJ strains of mice in the RR-7 dataset compared to C57BL/6J (RR-1) and BALB/c (RR-3), (Supplementary Figure 5).
