## Supplementary Tables for "Spaceflight causes strain dependent gene expression changes associated with lipid and extracellular matrix dysregulation in the mouse kidney in vivo"

Supplementary Table 1 C57BL/6J (RR-1 mission) differential gene expression analysis

| gene_id | gene_name | log2FoldChange | padj |
| --- | --- | --- | --- |
| ENSMUSG00000074715 | Ccl28 | 2.05038905 | 1.99E-05 |
| ENSMUSG00000032758 | Kap | 1.469959699 | 0.001036795 |
| ENSMUSG00000053303 | Slc22a26 | 1.452479228 | 0.016665891 |
| ENSMUSG00000031994 | Adamts8 | 1.334011867 | 0.001635329 |
| ENSMUSG00000021214 | Akr1c18 | 1.290958423 | 0.013901769 |
| ENSMUSG00000021135 | Slc10a1 | 1.236973328 | 4.85E-05 |
| ENSMUSG00000052562 | Slc22a30 | 1.194919329 | 0.099296306 |
| ENSMUSG00000026077 | Npas2 | 1.191666572 | 2.57E-09 |
| ENSMUSG00000112324 | Gm47939 | 1.147101435 | 0.012967515 |
| ENSMUSG00000054986 | Sec14l3 | 1.131443452 | 0.000982936 |
| ENSMUSG00000055116 | Arntl | 1.042131342 | 1.83E-07 |
| ENSMUSG00000056749 | Nfil3 | 1.028361512 | 0.000340607 |
| ENSMUSG00000026188 | Tmem169 | 0.971901354 | 6.00E-06 |
| ENSMUSG00000002250 | Ppard | 0.926172029 | 1.15E-12 |
| ENSMUSG00000023067 | Cdkn1a | 0.924416361 | 0.000463478 |
| ENSMUSG00000030111 | A2m | 0.899384744 | 0.052590918 |
| ENSMUSG00000110755 | BC049987 | 0.854525695 | 0.043271589 |
| ENSMUSG00000090264 | Eif4ebp3 | 0.848446111 | 9.41E-05 |
| ENSMUSG00000037263 | Aldh3b3 | 0.834503721 | 0.051420985 |
| ENSMUSG00000029752 | Asns | 0.832604884 | 0.000475925 |
| ENSMUSG00000042672 | Dcst1 | 0.832278691 | 0.055848938 |
| ENSMUSG00000028989 | Angptl7 | 0.830029211 | 0.050188786 |
| ENSMUSG00000046999 | 1110032F04Rik | 0.821024972 | 0.073622685 |
| ENSMUSG00000037725 | Ckap2 | 0.818864995 | 0.086208632 |
| ENSMUSG00000031283 | Chrdl1 | 0.818603142 | 0.003540241 |
| ENSMUSG00000043681 | Fam25c | 0.812977496 | 9.41E-05 |
| ENSMUSG00000069302 | Hist1h2ah | 0.812790382 | 0.078084014 |
| ENSMUSG00000032487 | Ptgs2 | 0.779654833 | 0.030335666 |
| ENSMUSG00000025185 | Loxl4 | 0.776543866 | 7.57E-05 |
| ENSMUSG00000038224 | Serpinf2 | 0.77350678 | 1.03E-06 |
| ENSMUSG00000022797 | Tfrc | 0.759068722 | 0.001365827 |
| ENSMUSG00000022021 | Diaph3 | 0.744110708 | 0.063530369 |
| ENSMUSG00000100131 | Gm28439 | 0.734736169 | 0.007185608 |
| ENSMUSG00000030137 | Tuba8 | 0.732793276 | 0.022891635 |
| ENSMUSG00000022033 | Pbk | 0.716994561 | 0.071455515 |
| ENSMUSG00000030641 | Ddias | 0.707720482 | 0.027975707 |
| ENSMUSG00000031594 | Fgl1 | 0.705116731 | 0.00235323 |
| ENSMUSG00000029093 | Sorcs2 | 0.702194523 | 0.032436674 |
| ENSMUSG00000041920 | Slc16a6 | 0.698753846 | 5.80E-05 |
| ENSMUSG00000024292 | Cyp4f14 | 0.697294267 | 0.024947552 |
| ENSMUSG00000050097 | Ces2b | 0.690980681 | 0.05884925 |
| ENSMUSG00000007783 | Cpt1c | 0.680805953 | 0.098480046 |
| ENSMUSG00000092500 | Gm20400 | 0.678386044 | 0.002015954 |
| ENSMUSG00000045328 | Cenpe | 0.675213153 | 0.051320636 |
| ENSMUSG00000023120 | Gm853 | 0.657358528 | 0.000982936 |
| ENSMUSG00000034457 | Eda2r | 0.656191502 | 0.005844588 |
| ENSMUSG00000035561 | Aldh1b1 | 0.628689205 | 0.009500191 |
| ENSMUSG00000031725 | Ces1f | 0.62721721 | 0.02289511 |
| ENSMUSG00000005667 | Mthfd2 | 0.619192987 | 0.086304013 |
| ENSMUSG00000033538 | Casp4 | 0.610839526 | 0.074068614 |
| ENSMUSG00000073405 | H2-T-ps | 0.607981493 | 0.080149399 |
| ENSMUSG00000022012 | Enox1 | 0.602244437 | 0.052661645 |
| ENSMUSG00000051397 | Tacstd2 | 0.601928042 | 0.078056757 |
| ENSMUSG00000067144 | Slc22a7 | 0.584008807 | 0.078859282 |
| ENSMUSG00000026827 | Gpd2 | 0.583975384 | 9.41E-05 |
| ENSMUSG00000037347 | Chst7 | 0.582535054 | 0.01639769 |
| ENSMUSG00000016942 | Tmprss6 | 0.572593728 | 0.048878865 |
| ENSMUSG00000020062 | Slc5a8 | 0.556719458 | 0.07808189 |
| ENSMUSG00000023259 | Slc26a6 | 0.549257871 | 0.006444439 |
| ENSMUSG00000032902 | Slc16a1 | 0.548042496 | 0.086304013 |
| ENSMUSG00000110663 | Gm39157 | 0.547440248 | 0.05884925 |
| ENSMUSG00000025735 | Rhbdl1 | 0.540775555 | 0.007281607 |
| ENSMUSG00000085042 | Abhd11os | 0.536832042 | 0.051021895 |
| ENSMUSG00000054520 | Sh3bp2 | 0.534768885 | 0.000798808 |
| ENSMUSG00000028671 | Gale | 0.528718156 | 0.000774565 |
| ENSMUSG00000045136 | Tubb2b | 0.526947999 | 0.007619929 |
| ENSMUSG00000002797 | Ggct | 0.525908913 | 4.26E-05 |
| ENSMUSG00000087574 | C030037D09Rik | 0.525288545 | 0.017000398 |
| ENSMUSG00000021670 | Hmgcr | 0.523407294 | 0.012038524 |
| ENSMUSG00000001473 | Tubb6 | 0.519899576 | 0.005733463 |
| ENSMUSG00000079261 | Gm15217 | 0.516998801 | 0.071455515 |
| ENSMUSG00000020142 | Slc1a4 | 0.512553471 | 0.040296576 |
| ENSMUSG00000041605 | Inava | 0.508780375 | 0.000584449 |
| ENSMUSG00000040998 | Npnt | 0.506192766 | 2.57E-09 |
| ENSMUSG00000027490 | E2f1 | 0.500784685 | 0.028558373 |
| ENSMUSG00000058258 | Idi1 | 0.500345924 | 0.016375499 |
| ENSMUSG00000046070 | Igfals | 0.49754579 | 0.051741568 |
| ENSMUSG00000082016 | Pgam1-ps2 | 0.4959904 | 0.043156319 |
| ENSMUSG00000026042 | Col5a2 | 0.491944942 | 0.055848938 |
| ENSMUSG00000036752 | Tubb4b | 0.489926174 | 0.003778971 |
| ENSMUSG00000042487 | Leo1 | 0.487964138 | 2.67E-05 |
| ENSMUSG00000070985 | Acnat1 | 0.48751268 | 0.000426005 |
| ENSMUSG00000037686 | Aspg | 0.487458754 | 0.031226072 |
| ENSMUSG00000032269 | Htr3a | 0.486818397 | 0.067211424 |
| ENSMUSG00000026189 | Pecr | 0.486046881 | 0.005733463 |
| ENSMUSG00000058672 | Tubb2a | 0.485918344 | 0.017000398 |
| ENSMUSG00000040694 | Apobec2 | 0.479796142 | 0.071940334 |
| ENSMUSG00000023153 | Tmem52 | 0.476166487 | 0.067860295 |
| ENSMUSG00000026202 | Tuba4a | 0.475482237 | 0.012802269 |
| ENSMUSG00000031604 | Msmo1 | 0.470866175 | 0.054083508 |
| ENSMUSG00000025726 | Slc28a1 | 0.470764251 | 0.036522168 |
| ENSMUSG00000064345 | mt-Nd2 | 0.469663447 | 0.046093823 |
| ENSMUSG00000109532 | Gm42397 | 0.468363611 | 0.047589144 |
| ENSMUSG00000095193 | Gm20939 | 0.468184944 | 0.020795761 |
| ENSMUSG00000033107 | Rnf125 | 0.466584681 | 0.014231293 |
| ENSMUSG00000021414 | Fam217a | 0.466482313 | 0.064182275 |
| ENSMUSG00000027111 | Itga6 | 0.466109898 | 2.94E-07 |
| ENSMUSG00000035910 | Dcdc2a | 0.465018776 | 0.002969322 |
| ENSMUSG00000022763 | Aifm3 | 0.462874333 | 0.000774565 |
| ENSMUSG00000026399 | Cd55 | 0.46283758 | 0.005175896 |
| ENSMUSG00000028937 | Acot7 | 0.45976607 | 0.033036876 |
| ENSMUSG00000024866 | Acy3 | 0.455022016 | 0.019979107 |
| ENSMUSG00000028834 | Trim63 | 0.452798502 | 0.093887443 |
| ENSMUSG00000025317 | Car5a | 0.452248799 | 0.039955273 |
| ENSMUSG00000036040 | Adamtsl2 | 0.448652621 | 0.027158221 |
| ENSMUSG00000038704 | Aspdh | 0.44198717 | 0.005969501 |
| ENSMUSG00000044786 | Zfp36 | 0.440638341 | 0.066903286 |
| ENSMUSG00000025203 | Scd2 | 0.440108985 | 0.06226633 |
| ENSMUSG00000028001 | Fga | 0.429273484 | 0.081207225 |
| ENSMUSG00000028327 | Stra6l | 0.429143877 | 0.091474624 |
| ENSMUSG00000019944 | Rhobtb1 | 0.429107109 | 0.07309444 |
| ENSMUSG00000043091 | Tuba1c | 0.428678477 | 1.13E-06 |
| ENSMUSG00000037169 | Mycn | 0.428358656 | 0.07351402 |
| ENSMUSG00000038775 | Vill | 0.428346096 | 0.030335666 |
| ENSMUSG00000064341 | mt-Nd1 | 0.427169144 | 0.024926943 |
| ENSMUSG00000096795 | Zfp433 | 0.425652551 | 0.036522168 |
| ENSMUSG00000101249 | Gm29216 | 0.423990077 | 0.009925648 |
| ENSMUSG00000053414 | Hunk | 0.423322133 | 0.002569009 |
| ENSMUSG00000038217 | Tlcd2 | 0.421612122 | 0.066284181 |
| ENSMUSG00000028919 | Arhgef19 | 0.421567952 | 0.003669156 |
| ENSMUSG00000059743 | Fdps | 0.419796818 | 0.000980637 |
| ENSMUSG00000054951 | 9130008F23Rik | 0.419644953 | 0.082774985 |
| ENSMUSG00000020917 | Acly | 0.413591433 | 0.07788242 |
| ENSMUSG00000032978 | Guca2b | 0.412211622 | 0.023475892 |
| ENSMUSG00000045294 | Insig1 | 0.410191526 | 0.030260779 |
| ENSMUSG00000041889 | Shisa4 | 0.407321431 | 0.00629571 |
| ENSMUSG00000037386 | Rims2 | 0.406450124 | 0.087838455 |
| ENSMUSG00000029361 | Nos1 | 0.401496088 | 0.012518038 |
| ENSMUSG00000062248 | Cks2 | 0.401141399 | 0.080344478 |
| ENSMUSG00000026249 | Serpine2 | 0.397704824 | 0.079284342 |
| ENSMUSG00000064367 | mt-Nd5 | 0.397008306 | 0.021509346 |
| ENSMUSG00000028780 | Sema3c | 0.389935152 | 0.037336983 |
| ENSMUSG00000032420 | Nt5e | 0.387428233 | 0.00025849 |
| ENSMUSG00000031349 | Nsdhl | 0.386506259 | 0.006454113 |
| ENSMUSG00000029771 | Irf5 | 0.385539952 | 0.087838455 |
| ENSMUSG00000006800 | Sulf2 | 0.383640593 | 0.017129383 |
| ENSMUSG00000037243 | Zfp692 | 0.383396499 | 0.012834796 |
| ENSMUSG00000052117 | D630039A03Rik | 0.383077029 | 0.03685592 |
| ENSMUSG00000010663 | Fads1 | 0.376745102 | 0.003639494 |
| ENSMUSG00000030880 | Polr3e | 0.374304273 | 0.036449117 |
| ENSMUSG00000022351 | Sqle | 0.374253892 | 0.001774558 |
| ENSMUSG00000028357 | Kif12 | 0.374218425 | 0.000833718 |
| ENSMUSG00000039062 | Anpep | 0.373939107 | 0.001608848 |
| ENSMUSG00000021185 | Dglucy | 0.373914064 | 0.001041817 |
| ENSMUSG00000074261 | Erich4 | 0.372593034 | 1.22E-07 |
| ENSMUSG00000041939 | Mvk | 0.369203934 | 0.009338305 |
| ENSMUSG00000068876 | Cgn | 0.368285307 | 0.020225076 |
| ENSMUSG00000001627 | Ifrd1 | 0.367505566 | 0.010817814 |
| ENSMUSG00000032452 | Clstn2 | 0.365696327 | 0.031226072 |
| ENSMUSG00000056608 | Chd9 | 0.361874999 | 0.085858965 |
| ENSMUSG00000027230 | Creb3l1 | 0.361553207 | 0.056239177 |
| ENSMUSG00000006517 | Mvd | 0.36050399 | 5.95E-05 |
| ENSMUSG00000000253 | Gmpr | 0.359425712 | 0.000596029 |
| ENSMUSG00000090236 | Car15 | 0.358570488 | 0.001037718 |
| ENSMUSG00000034371 | Tkfc | 0.356288058 | 0.041615056 |
| ENSMUSG00000020381 | Mrnip | 0.355262836 | 0.006444439 |
| ENSMUSG00000021364 | Elovl2 | 0.353395395 | 0.000157116 |
| ENSMUSG00000000594 | Gm2a | 0.353170161 | 0.005733463 |
| ENSMUSG00000064254 | Ethe1 | 0.352788237 | 0.04331772 |
| ENSMUSG00000028545 | Bend5 | 0.348990484 | 0.007185608 |
| ENSMUSG00000020326 | Ccng1 | 0.34825388 | 8.49E-05 |
| ENSMUSG00000045725 | Prr15 | 0.348221185 | 0.078056757 |
| ENSMUSG00000079465 | Col4a3 | 0.347470745 | 0.000442467 |
| ENSMUSG00000063694 | Cycs | 0.345785071 | 0.003839093 |
| ENSMUSG00000014245 | Pigl | 0.342699277 | 0.033014186 |
| ENSMUSG00000102070 | Gm28661 | 0.34206471 | 0.088320188 |
| ENSMUSG00000032786 | Alas1 | 0.341831516 | 0.093276198 |
| ENSMUSG00000022580 | Rhpn1 | 0.34032725 | 4.85E-05 |
| ENSMUSG00000024818 | Slc25a45 | 0.33997817 | 0.05884925 |
| ENSMUSG00000055725 | Paqr3 | 0.339046304 | 0.085858965 |
| ENSMUSG00000024579 | Pcyox1l | 0.338412847 | 0.011984919 |
| ENSMUSG00000020538 | Srebf1 | 0.337943092 | 0.01793752 |
| ENSMUSG00000026784 | Pdss1 | 0.336973303 | 0.021784448 |
| ENSMUSG00000032374 | Plod2 | 0.330979395 | 9.41E-05 |
| ENSMUSG00000055745 | Rtl6 | 0.330089448 | 0.01439599 |
| ENSMUSG00000071076 | Jund | 0.329213603 | 0.078833669 |
| ENSMUSG00000068220 | Lgals1 | 0.327122027 | 0.046093823 |
| ENSMUSG00000047986 | Palm3 | 0.326923089 | 0.087838455 |
| ENSMUSG00000031958 | Ldhd | 0.326053677 | 0.0250321 |
| ENSMUSG00000074170 | Plekhf1 | 0.322829038 | 0.027732583 |
| ENSMUSG00000006342 | Susd2 | 0.322248391 | 0.037169542 |
| ENSMUSG00000024064 | Galnt14 | 0.321778231 | 0.003715331 |
| ENSMUSG00000058454 | Dhcr7 | 0.321526109 | 0.007185608 |
| ENSMUSG00000029763 | Exoc4 | 0.321493893 | 0.001127832 |
| ENSMUSG00000045594 | Glb1 | 0.321333845 | 0.018191818 |
| ENSMUSG00000037455 | Slc18b1 | 0.31849381 | 0.000315527 |
| ENSMUSG00000029161 | Cgref1 | 0.315727902 | 0.080344478 |
| ENSMUSG00000026627 | Tmem206 | 0.313419393 | 0.054921911 |
| ENSMUSG00000022512 | Cldn1 | 0.31331514 | 0.012543547 |
| ENSMUSG00000038375 | Trp53inp2 | 0.311184963 | 7.41E-05 |
| ENSMUSG00000032515 | Csrnp1 | 0.307541561 | 0.054921911 |
| ENSMUSG00000020877 | Scrn2 | 0.304645533 | 0.01639769 |
| ENSMUSG00000029304 | Spp1 | 0.30369885 | 0.059015174 |
| ENSMUSG00000041959 | S100a10 | 0.302227305 | 0.014231293 |
| ENSMUSG00000024503 | Spink1 | 0.29885584 | 0.047794424 |
| ENSMUSG00000023938 | Aars2 | 0.298276717 | 0.001366557 |
| ENSMUSG00000018427 | Ypel2 | 0.297810257 | 0.080344478 |
| ENSMUSG00000030512 | Snrpa1 | 0.29675962 | 0.017594875 |
| ENSMUSG00000027327 | 1700037H04Rik | 0.296103972 | 0.075810233 |
| ENSMUSG00000047735 | Samd9l | 0.295528893 | 0.028746387 |
| ENSMUSG00000024736 | Tmem132a | 0.294505567 | 0.0337821 |
| ENSMUSG00000047281 | Sfn | 0.292135768 | 0.06113424 |
| ENSMUSG00000011382 | Dhdh | 0.288037163 | 0.012098151 |
| ENSMUSG00000023004 | Tuba1b | 0.285637832 | 0.054551942 |
| ENSMUSG00000024665 | Fads2 | 0.282910545 | 0.03890022 |
| ENSMUSG00000038527 | C1rl | 0.277620322 | 0.085004646 |
| ENSMUSG00000026179 | Pnkd | 0.277439271 | 0.00024083 |
| ENSMUSG00000032370 | Lactb | 0.276873814 | 0.010346206 |
| ENSMUSG00000063229 | Ldha | 0.273680834 | 0.027732583 |
| ENSMUSG00000015094 | Npdc1 | 0.273526976 | 0.019943321 |
| ENSMUSG00000057103 | Nat8f1 | 0.272063939 | 0.061455649 |
| ENSMUSG00000021792 | Fam213a | 0.270643824 | 0.0163947 |
| ENSMUSG00000022893 | Adamts1 | 0.269553148 | 0.070083852 |
| ENSMUSG00000040263 | Klhdc4 | 0.26655755 | 0.001335702 |
| ENSMUSG00000074890 | Lcmt2 | 0.264466553 | 0.07826053 |
| ENSMUSG00000019797 | 1700021F05Rik | 0.259594847 | 0.020795761 |
| ENSMUSG00000030739 | Myh14 | 0.256711749 | 0.0117542 |
| ENSMUSG00000048707 | Tprn | 0.255748891 | 0.001567332 |
| ENSMUSG00000093930 | Hmgcs1 | 0.255731474 | 0.098480046 |
| ENSMUSG00000032911 | Cspg4 | 0.252971242 | 0.000698378 |
| ENSMUSG00000034714 | Ttyh2 | 0.250682852 | 0.030252841 |
| ENSMUSG00000015357 | Clpx | 0.249967504 | 0.054921911 |
| ENSMUSG00000014158 | Trpv4 | 0.244908345 | 0.000116795 |
| ENSMUSG00000021697 | Depdc1b | 0.243400063 | 0.016375499 |
| ENSMUSG00000023832 | Acat2 | 0.241505003 | 0.046102028 |
| ENSMUSG00000004266 | Ptpn6 | 0.240724383 | 0.010817814 |
| ENSMUSG00000034858 | Fam214a | 0.240511302 | 0.027975707 |
| ENSMUSG00000069633 | Pex11g | 0.240501216 | 0.025898344 |
| ENSMUSG00000024993 | Fam45a | 0.239985928 | 0.028360939 |
| ENSMUSG00000032492 | Pth1r | 0.239060449 | 0.043271589 |
| ENSMUSG00000020744 | Slc25a19 | 0.238704786 | 0.014231293 |
| ENSMUSG00000027412 | Lpin3 | 0.237946331 | 0.018651528 |
| ENSMUSG00000051518 | Rps19bp1 | 0.237907293 | 0.03695359 |
| ENSMUSG00000036957 | Lrfn3 | 0.235445909 | 0.016109538 |
| ENSMUSG00000037103 | Dcaf15 | 0.233550555 | 0.098480046 |
| ENSMUSG00000069565 | Dazap1 | 0.232676629 | 0.000982936 |
| ENSMUSG00000063897 | CAAA01118383.1 | 0.232221238 | 0.061455649 |
| ENSMUSG00000031960 | Aars | 0.231409025 | 0.012543547 |
| ENSMUSG00000035845 | Alg12 | 0.230291975 | 0.046289942 |
| ENSMUSG00000057363 | Uxs1 | 0.230017303 | 0.022315585 |
| ENSMUSG00000046079 | Lrrc8d | 0.229541611 | 0.003540241 |
| ENSMUSG00000028041 | Adam15 | 0.227383375 | 0.012834796 |
| ENSMUSG00000078695 | Cisd3 | 0.226214046 | 0.072598408 |
| ENSMUSG00000032115 | Hyou1 | 0.225582333 | 0.097762914 |
| ENSMUSG00000029580 | Actb | 0.225211422 | 0.013347737 |
| ENSMUSG00000067924 | Rtl8b | 0.223338932 | 0.056239177 |
| ENSMUSG00000022540 | Rogdi | 0.221750807 | 0.00801318 |
| ENSMUSG00000023367 | Tmem176a | 0.221668758 | 0.000463478 |
| ENSMUSG00000020116 | Pno1 | 0.220740825 | 0.051320636 |
| ENSMUSG00000031781 | Ciapin1 | 0.219566251 | 0.074068614 |
| ENSMUSG00000053291 | Rab4b | 0.21890932 | 0.099099876 |
| ENSMUSG00000070283 | Ndufaf3 | 0.217502502 | 0.054551942 |
| ENSMUSG00000024958 | Gpr137 | 0.216364931 | 0.00185365 |
| ENSMUSG00000061474 | Mrps36 | 0.215993386 | 0.093480306 |
| ENSMUSG00000011752 | Pgam1 | 0.21550453 | 0.029858907 |
| ENSMUSG00000024772 | Ehd1 | 0.214047351 | 0.025513946 |
| ENSMUSG00000028538 | St3gal3 | 0.21401319 | 0.020264734 |
| ENSMUSG00000062825 | Actg1 | 0.213667766 | 0.037835649 |
| ENSMUSG00000020232 | Hmg20b | 0.213503798 | 0.00041236 |
| ENSMUSG00000029810 | Tmem176b | 0.21168028 | 0.011220415 |
| ENSMUSG00000000934 | Top1mt | 0.210039157 | 0.036944328 |
| ENSMUSG00000030538 | Cib1 | 0.20987864 | 0.02289511 |
| ENSMUSG00000018171 | Vmp1 | 0.209804739 | 0.019433008 |
| ENSMUSG00000039834 | Zfp335 | 0.209200674 | 0.003783071 |
| ENSMUSG00000036452 | Arhgap26 | 0.208864752 | 0.08550327 |
| ENSMUSG00000027610 | Gss | 0.207832464 | 0.080344478 |
| ENSMUSG00000020307 | Cdc34 | 0.207684849 | 0.093276198 |
| ENSMUSG00000066441 | Rdh11 | 0.205818375 | 0.00801318 |
| ENSMUSG00000032349 | Elovl5 | 0.204397323 | 0.019910688 |
| ENSMUSG00000084128 | Esrp2 | 0.20438744 | 0.032527597 |
| ENSMUSG00000039047 | Pigk | 0.199720926 | 0.001388697 |
| ENSMUSG00000034613 | Ppm1h | 0.197665087 | 0.031257651 |
| ENSMUSG00000032306 | Mpi | 0.19731545 | 0.012834796 |
| ENSMUSG00000027429 | Sec23b | 0.196725744 | 0.018651528 |
| ENSMUSG00000031161 | Hdac6 | 0.192697657 | 0.018444046 |
| ENSMUSG00000004843 | Chmp2b | 0.192558241 | 0.050216875 |
| ENSMUSG00000020775 | Mrpl38 | 0.192405317 | 0.099391328 |
| ENSMUSG00000017428 | Psmd11 | 0.191427343 | 0.00422981 |
| ENSMUSG00000020571 | Pdia6 | 0.188969722 | 0.096614042 |
| ENSMUSG00000026914 | Psmd14 | 0.187919964 | 0.099296306 |
| ENSMUSG00000030298 | Sec13 | 0.187370656 | 0.050216875 |
| ENSMUSG00000004565 | Pnpla6 | 0.185041292 | 0.006552273 |
| ENSMUSG00000048578 | Mlec | 0.184530741 | 0.050188786 |
| ENSMUSG00000023048 | Prr13 | 0.184435007 | 0.095782346 |
| ENSMUSG00000017210 | Med24 | 0.184066054 | 0.046658599 |
| ENSMUSG00000026036 | Nif3l1 | 0.182497312 | 0.099337335 |
| ENSMUSG00000028042 | Zbtb7b | 0.181559675 | 0.087838455 |
| ENSMUSG00000040188 | Scamp2 | 0.181506267 | 0.00457556 |
| ENSMUSG00000026307 | Scly | 0.179758442 | 0.013901769 |
| ENSMUSG00000036002 | Fam214b | 0.179124298 | 0.078056757 |
| ENSMUSG00000003813 | Rad23a | 0.17697052 | 0.071455515 |
| ENSMUSG00000020898 | Ctc1 | 0.175823619 | 0.079554987 |
| ENSMUSG00000028063 | Lmna | 0.175656721 | 0.060798796 |
| ENSMUSG00000024683 | Mrpl16 | 0.174069837 | 0.078056757 |
| ENSMUSG00000029032 | Arhgef16 | 0.17262869 | 0.00653581 |
| ENSMUSG00000021666 | Gfm2 | 0.171767708 | 0.067211424 |
| ENSMUSG00000039254 | Pomt1 | 0.171069372 | 0.079189176 |
| ENSMUSG00000029616 | Erp29 | 0.169967609 | 0.020795761 |
| ENSMUSG00000062203 | Gspt1 | 0.169290633 | 0.007281607 |
| ENSMUSG00000026201 | Stk16 | 0.168202419 | 0.085004646 |
| ENSMUSG00000067158 | Col4a4 | 0.166985426 | 0.025454787 |
| ENSMUSG00000041733 | Coq5 | 0.163919062 | 0.040569049 |
| ENSMUSG00000038023 | Atp6v0a2 | 0.162182571 | 0.010890155 |
| ENSMUSG00000024319 | Vps52 | 0.161379743 | 0.024029852 |
| ENSMUSG00000048277 | Syngr2 | 0.161239228 | 0.07808189 |
| ENSMUSG00000029701 | Rbm28 | 0.154783445 | 0.071455515 |
| ENSMUSG00000029394 | Cdk2ap1 | 0.15465162 | 0.068186546 |
| ENSMUSG00000030805 | Stx4a | 0.153477996 | 0.093887443 |
| ENSMUSG00000002393 | Nr2f6 | 0.151142992 | 0.061836061 |
| ENSMUSG00000025762 | Larp1b | 0.149923704 | 0.081278579 |
| ENSMUSG00000046603 | Tcaim | 0.14965173 | 0.085004646 |
| ENSMUSG00000037894 | H2afz | 0.149238696 | 0.017129383 |
| ENSMUSG00000035863 | Palm | 0.135989184 | 0.064874932 |
| ENSMUSG00000037999 | Arap2 | 0.135798074 | 0.050216875 |
| ENSMUSG00000031776 | Arl2bp | 0.132990816 | 0.097825476 |
| ENSMUSG00000058569 | Tmed9 | 0.131274183 | 0.078056757 |
| ENSMUSG00000037470 | Uggt1 | 0.127728029 | 0.071940334 |
| ENSMUSG00000020827 | Mink1 | 0.091564681 | 0.067937874 |
| ENSMUSG00000020716 | Nf1 | -0.106850424 | 0.067964917 |
| ENSMUSG00000038056 | Kmt2c | -0.107425107 | 0.091123501 |
| ENSMUSG00000044791 | Setd2 | -0.108027523 | 0.031213423 |
| ENSMUSG00000053110 | Yap1 | -0.11290101 | 0.065679369 |
| ENSMUSG00000005886 | Ncoa2 | -0.113612716 | 0.076581842 |
| ENSMUSG00000048170 | Mcmbp | -0.118635025 | 0.086304013 |
| ENSMUSG00000036550 | Cnot1 | -0.121668659 | 0.000833718 |
| ENSMUSG00000021144 | Mta1 | -0.122928694 | 0.096708008 |
| ENSMUSG00000050812 | Ecpas | -0.123290928 | 0.008683307 |
| ENSMUSG00000037822 | Smim14 | -0.123593011 | 0.045089513 |
| ENSMUSG00000015290 | Ubl4a | -0.123644763 | 0.093276198 |
| ENSMUSG00000033943 | Mga | -0.124148128 | 0.079189176 |
| ENSMUSG00000033237 | Arid2 | -0.125128311 | 0.066284181 |
| ENSMUSG00000027519 | Rab22a | -0.127134209 | 0.014216707 |
| ENSMUSG00000005893 | Nr2c2 | -0.127295956 | 0.031213423 |
| ENSMUSG00000052751 | Repin1 | -0.127322678 | 0.08652197 |
| ENSMUSG00000027799 | Nbea | -0.127814612 | 0.061067792 |
| ENSMUSG00000028053 | Ash1l | -0.129462147 | 0.002905291 |
| ENSMUSG00000065954 | Tacc1 | -0.129693827 | 0.093276198 |
| ENSMUSG00000014195 | Dnajc7 | -0.131979397 | 0.038856141 |
| ENSMUSG00000033671 | Cep350 | -0.132238603 | 0.078056757 |
| ENSMUSG00000037503 | Fam168b | -0.13225204 | 0.044472308 |
| ENSMUSG00000020594 | Pum2 | -0.132620328 | 0.050188786 |
| ENSMUSG00000025195 | Dnmbp | -0.132700766 | 0.098480046 |
| ENSMUSG00000034893 | Cog3 | -0.133164219 | 0.088476599 |
| ENSMUSG00000102869 | 2900097C17Rik | -0.133559213 | 0.041334035 |
| ENSMUSG00000074748 | Atxn7l3b | -0.135084594 | 0.011984919 |
| ENSMUSG00000079215 | Zfp664 | -0.135660638 | 0.055848938 |
| ENSMUSG00000031729 | Ist1 | -0.135735051 | 0.013901769 |
| ENSMUSG00000005698 | Ctcf | -0.136316828 | 0.05399549 |
| ENSMUSG00000039801 | Cplane1 | -0.136569005 | 0.080163531 |
| ENSMUSG00000040481 | Bptf | -0.136771119 | 0.087838455 |
| ENSMUSG00000009470 | Tnpo1 | -0.136979355 | 0.088348954 |
| ENSMUSG00000046897 | Zfp740 | -0.137668354 | 0.020795761 |
| ENSMUSG00000036990 | Otud4 | -0.137927447 | 0.095782346 |
| ENSMUSG00000062866 | Phactr2 | -0.138936951 | 0.099007605 |
| ENSMUSG00000021140 | Pcnx | -0.139916849 | 0.026942687 |
| ENSMUSG00000021488 | Nsd1 | -0.140158118 | 0.016607464 |
| ENSMUSG00000018076 | Med13l | -0.140244264 | 0.024037471 |
| ENSMUSG00000027678 | Ncoa3 | -0.140743803 | 0.055233018 |
| ENSMUSG00000041530 | Ago1 | -0.141746548 | 0.006979568 |
| ENSMUSG00000034297 | Med13 | -0.141757281 | 0.045510148 |
| ENSMUSG00000017776 | Crk | -0.142838617 | 9.66E-05 |
| ENSMUSG00000043991 | Pura | -0.143382981 | 0.021980406 |
| ENSMUSG00000035247 | Hectd1 | -0.144303706 | 0.018444046 |
| ENSMUSG00000038437 | Mllt6 | -0.144577731 | 0.084726052 |
| ENSMUSG00000021395 | Spin1 | -0.145330564 | 0.045217761 |
| ENSMUSG00000040865 | Ino80d | -0.146496084 | 0.032688442 |
| ENSMUSG00000066415 | Msl2 | -0.146774214 | 0.051420985 |
| ENSMUSG00000044674 | Fzd1 | -0.147244976 | 0.041282549 |
| ENSMUSG00000049739 | Zfp646 | -0.147848957 | 0.076742331 |
| ENSMUSG00000071660 | Ttc9c | -0.148650337 | 0.057470148 |
| ENSMUSG00000040209 | Zfp704 | -0.149028025 | 0.024656018 |
| ENSMUSG00000035284 | Vps13c | -0.149859759 | 0.056239177 |
| ENSMUSG00000026918 | Brd3 | -0.15077549 | 0.011364663 |
| ENSMUSG00000029647 | Pan3 | -0.151105524 | 0.033842872 |
| ENSMUSG00000026436 | Elk4 | -0.15139194 | 0.019979107 |
| ENSMUSG00000005034 | Prkacb | -0.151987266 | 0.010346206 |
| ENSMUSG00000037712 | Fermt2 | -0.152049033 | 0.091123501 |
| ENSMUSG00000057133 | Chd6 | -0.152095793 | 0.02067999 |
| ENSMUSG00000031754 | Nudt21 | -0.152344964 | 0.091869095 |
| ENSMUSG00000041037 | Irgq | -0.152620384 | 0.078833669 |
| ENSMUSG00000046480 | Scn4b | -0.152833582 | 0.06874662 |
| ENSMUSG00000026782 | Abi2 | -0.15361853 | 0.065679369 |
| ENSMUSG00000026313 | Hdac4 | -0.154054248 | 0.067211424 |
| ENSMUSG00000074994 | Qser1 | -0.15416519 | 0.051839757 |
| ENSMUSG00000060938 | Rpl26 | -0.154282648 | 0.079189176 |
| ENSMUSG00000046792 | Zfp787 | -0.154344083 | 0.05884925 |
| ENSMUSG00000043241 | Upf2 | -0.154546688 | 0.028360939 |
| ENSMUSG00000027680 | Fxr1 | -0.154698054 | 0.021680264 |
| ENSMUSG00000024431 | Nr3c1 | -0.154968185 | 0.061786262 |
| ENSMUSG00000036197 | Gxylt1 | -0.155303906 | 0.07826053 |
| ENSMUSG00000043716 | Rpl7 | -0.15558352 | 0.04331772 |
| ENSMUSG00000061477 | Rps7 | -0.155800783 | 0.080874608 |
| ENSMUSG00000037736 | Limch1 | -0.156050576 | 0.032198638 |
| ENSMUSG00000009927 | Rps25 | -0.15622053 | 0.031568976 |
| ENSMUSG00000020448 | Rnf185 | -0.15643565 | 0.073258652 |
| ENSMUSG00000018736 | Ndel1 | -0.156449666 | 0.097365008 |
| ENSMUSG00000047888 | Tnrc6b | -0.15740914 | 0.098480046 |
| ENSMUSG00000050310 | Rictor | -0.157409315 | 0.000228531 |
| ENSMUSG00000028869 | Gnl2 | -0.157501858 | 0.06874662 |
| ENSMUSG00000058793 | Cds2 | -0.157989721 | 0.017194333 |
| ENSMUSG00000039108 | Lsm14b | -0.159074854 | 0.07452549 |
| ENSMUSG00000002748 | Baz1b | -0.160226702 | 0.098480046 |
| ENSMUSG00000032086 | Bace1 | -0.160392678 | 0.007281607 |
| ENSMUSG00000037029 | Zfp146 | -0.161542295 | 0.016109538 |
| ENSMUSG00000035495 | Tstd2 | -0.161953855 | 0.040259159 |
| ENSMUSG00000038481 | Cdk19 | -0.163411879 | 0.08645857 |
| ENSMUSG00000025997 | Ikzf2 | -0.165100929 | 0.097762914 |
| ENSMUSG00000020257 | Wdr82 | -0.165418315 | 0.001036795 |
| ENSMUSG00000039068 | Zzz3 | -0.165642262 | 0.000419273 |
| ENSMUSG00000031393 | Mecp2 | -0.165659422 | 0.000461435 |
| ENSMUSG00000051864 | Tbc1d22a | -0.167305619 | 0.053264773 |
| ENSMUSG00000045098 | Kmt5b | -0.167471537 | 0.010714927 |
| ENSMUSG00000019866 | Crybg1 | -0.167931404 | 0.037336983 |
| ENSMUSG00000027782 | Kpna4 | -0.168789383 | 0.070926832 |
| ENSMUSG00000037369 | Kdm6a | -0.168869656 | 0.017594875 |
| ENSMUSG00000044617 | Zbtb39 | -0.169029725 | 0.063338978 |
| ENSMUSG00000025571 | Tnrc6c | -0.169842205 | 0.086796893 |
| ENSMUSG00000021540 | Smad5 | -0.169855239 | 0.000419273 |
| ENSMUSG00000001280 | Sp1 | -0.169926102 | 0.001465044 |
| ENSMUSG00000036097 | Slf2 | -0.170332721 | 0.045217761 |
| ENSMUSG00000022353 | Mtss1 | -0.170564489 | 0.028758887 |
| ENSMUSG00000060657 | Marf1 | -0.171029433 | 0.003447958 |
| ENSMUSG00000021669 | Col4a3bp | -0.173176912 | 0.014216707 |
| ENSMUSG00000003970 | Rpl8 | -0.174705885 | 0.071455515 |
| ENSMUSG00000049800 | Sertad2 | -0.174955207 | 0.079189176 |
| ENSMUSG00000040123 | Zmym5 | -0.175892472 | 0.068658676 |
| ENSMUSG00000035614 | Togaram1 | -0.176366776 | 0.018149642 |
| ENSMUSG00000043384 | Gprasp1 | -0.176571334 | 0.056813269 |
| ENSMUSG00000069495 | Epc2 | -0.17725049 | 0.04756096 |
| ENSMUSG00000037523 | Mavs | -0.177616086 | 0.007281607 |
| ENSMUSG00000056851 | Pcbp2 | -0.178232544 | 0.027465928 |
| ENSMUSG00000017418 | Arl5b | -0.178449877 | 0.078090673 |
| ENSMUSG00000008683 | Rps15a | -0.178911585 | 0.051420985 |
| ENSMUSG00000032594 | Ip6k1 | -0.179217038 | 0.001365614 |
| ENSMUSG00000031209 | Heph | -0.180247413 | 0.056813269 |
| ENSMUSG00000038290 | Smg6 | -0.181402516 | 0.009794965 |
| ENSMUSG00000031618 | Nr3c2 | -0.181651925 | 0.06874662 |
| ENSMUSG00000037742 | Eef1a1 | -0.182891829 | 0.044794459 |
| ENSMUSG00000048379 | Socs4 | -0.182903932 | 0.076742331 |
| ENSMUSG00000060036 | Rpl3 | -0.18327048 | 0.056813269 |
| ENSMUSG00000035666 | Gtf3c4 | -0.183607105 | 0.032798546 |
| ENSMUSG00000022791 | Tnk2 | -0.184624712 | 0.056239177 |
| ENSMUSG00000031216 | Stard8 | -0.185167267 | 0.006680648 |
| ENSMUSG00000097119 | B230354K17Rik | -0.186122964 | 0.076007999 |
| ENSMUSG00000034647 | Ankrd12 | -0.187390834 | 0.081278579 |
| ENSMUSG00000006333 | Rps9 | -0.188109549 | 0.057923754 |
| ENSMUSG00000039831 | Arhgap29 | -0.189164654 | 0.042305009 |
| ENSMUSG00000035469 | Rcbtb1 | -0.189666369 | 0.016109538 |
| ENSMUSG00000090083 | Rnf8 | -0.189949604 | 0.086304013 |
| ENSMUSG00000058600 | Rpl30 | -0.190915457 | 0.016605138 |
| ENSMUSG00000028522 | Mier1 | -0.191087177 | 0.054011335 |
| ENSMUSG00000059486 | Kbtbd2 | -0.191497338 | 0.032198638 |
| ENSMUSG00000063317 | Usp31 | -0.191776939 | 0.069720765 |
| ENSMUSG00000006269 | Atp6v1b1 | -0.193624815 | 0.010541185 |
| ENSMUSG00000038766 | Gabpb2 | -0.193660125 | 0.010346206 |
| ENSMUSG00000071415 | Rpl23 | -0.194422227 | 0.00566107 |
| ENSMUSG00000033721 | Vav3 | -0.194583437 | 0.063530369 |
| ENSMUSG00000050947 | Amigo1 | -0.195095644 | 0.010346206 |
| ENSMUSG00000008435 | Rdh13 | -0.196053935 | 0.056239177 |
| ENSMUSG00000040363 | Bcor | -0.196951628 | 0.0945368 |
| ENSMUSG00000032097 | Ddx6 | -0.197635395 | 0.00041236 |
| ENSMUSG00000042046 | Dstyk | -0.198866102 | 0.000483936 |
| ENSMUSG00000025223 | Ldb1 | -0.199152531 | 0.007185608 |
| ENSMUSG00000020372 | Rack1 | -0.199949779 | 0.016665891 |
| ENSMUSG00000043929 | Klhl15 | -0.200441365 | 0.085052514 |
| ENSMUSG00000021959 | Lats2 | -0.200856073 | 0.04756096 |
| ENSMUSG00000027351 | Spred1 | -0.201204476 | 0.003778971 |
| ENSMUSG00000033411 | Ctdspl2 | -0.202957697 | 0.078090673 |
| ENSMUSG00000062519 | Zfp398 | -0.203093243 | 0.056239177 |
| ENSMUSG00000035530 | Eif1 | -0.204451626 | 0.043271589 |
| ENSMUSG00000029714 | Gigyf1 | -0.204520878 | 0.02289511 |
| ENSMUSG00000051817 | Sox12 | -0.205108357 | 0.014231293 |
| ENSMUSG00000040423 | Rc3h1 | -0.206818255 | 0.003216453 |
| ENSMUSG00000075592 | Nynrin | -0.208753576 | 0.073622685 |
| ENSMUSG00000034111 | Tmed8 | -0.211420958 | 0.002200337 |
| ENSMUSG00000047215 | Rpl9 | -0.211934122 | 0.076020626 |
| ENSMUSG00000052446 | Zfp961 | -0.213572821 | 0.024485496 |
| ENSMUSG00000012848 | Rps5 | -0.215368326 | 0.011364663 |
| ENSMUSG00000019817 | Plagl1 | -0.216553972 | 0.070083852 |
| ENSMUSG00000032624 | Eml4 | -0.217073377 | 0.003783071 |
| ENSMUSG00000024298 | Zfp871 | -0.217188874 | 0.012802269 |
| ENSMUSG00000037172 | E330009J07Rik | -0.218758613 | 0.004227134 |
| ENSMUSG00000054793 | Cadm4 | -0.220122063 | 0.000228531 |
| ENSMUSG00000020357 | Flt4 | -0.220165294 | 0.052388405 |
| ENSMUSG00000021482 | Aaed1 | -0.22075476 | 0.017210925 |
| ENSMUSG00000053604 | Rpia | -0.222340686 | 0.0700399 |
| ENSMUSG00000034189 | Hsdl1 | -0.222649745 | 0.021784448 |
| ENSMUSG00000006216 | Clcnkb | -0.223285134 | 0.09182984 |
| ENSMUSG00000038486 | Sv2a | -0.223605738 | 0.03384048 |
| ENSMUSG00000026694 | Mettl13 | -0.223860786 | 0.06113424 |
| ENSMUSG00000029154 | Cwh43 | -0.224438567 | 0.099296306 |
| ENSMUSG00000017405 | Nek8 | -0.225613943 | 0.0816179 |
| ENSMUSG00000025764 | Jade1 | -0.226066456 | 0.000419273 |
| ENSMUSG00000038544 | Inip | -0.226245474 | 0.089488849 |
| ENSMUSG00000030213 | Atf7ip | -0.228136167 | 0.002617042 |
| ENSMUSG00000028081 | Rps3a1 | -0.229029499 | 0.00117473 |
| ENSMUSG00000046947 | Adck2 | -0.229327906 | 0.016992884 |
| ENSMUSG00000023022 | Lima1 | -0.229765504 | 0.002126867 |
| ENSMUSG00000017667 | Zfp334 | -0.232434461 | 0.091123501 |
| ENSMUSG00000027395 | Polr1b | -0.232567625 | 0.015766717 |
| ENSMUSG00000092558 | Med20 | -0.232719535 | 0.036163942 |
| ENSMUSG00000044026 | Slc35g1 | -0.232841001 | 0.010346206 |
| ENSMUSG00000039789 | Zfp597 | -0.233102758 | 0.08652197 |
| ENSMUSG00000021661 | Ankra2 | -0.23600207 | 0.047944462 |
| ENSMUSG00000006127 | Inpp5k | -0.237129037 | 0.000130059 |
| ENSMUSG00000078202 | Nrarp | -0.237634947 | 0.040002948 |
| ENSMUSG00000066235 | Pomgnt2 | -0.237946664 | 0.051839757 |
| ENSMUSG00000073678 | Pgap1 | -0.23812418 | 0.01639769 |
| ENSMUSG00000026923 | Notch1 | -0.238274961 | 0.041565148 |
| ENSMUSG00000008682 | Rpl10 | -0.238914439 | 0.014216707 |
| ENSMUSG00000029587 | Zfp12 | -0.242629421 | 0.045473507 |
| ENSMUSG00000031365 | Zfp275 | -0.243556594 | 0.054255405 |
| ENSMUSG00000027079 | Clp1 | -0.244797916 | 0.050188786 |
| ENSMUSG00000022604 | Cep97 | -0.24515406 | 0.001919493 |
| ENSMUSG00000043090 | Zfp866 | -0.245371449 | 0.015211362 |
| ENSMUSG00000047878 | A4galt | -0.246859705 | 0.064083097 |
| ENSMUSG00000041378 | Cldn5 | -0.247374697 | 0.017194333 |
| ENSMUSG00000041841 | Rpl37 | -0.249690419 | 0.042863719 |
| ENSMUSG00000021180 | Rps6ka5 | -0.250735537 | 0.097762914 |
| ENSMUSG00000037003 | Tns2 | -0.254353497 | 0.010714927 |
| ENSMUSG00000025019 | Lcor | -0.255186551 | 8.33E-05 |
| ENSMUSG00000038677 | Scube3 | -0.256086003 | 0.053244425 |
| ENSMUSG00000030201 | Lrp6 | -0.25623567 | 0.016316005 |
| ENSMUSG00000031320 | Rps4x | -0.256282323 | 0.00026316 |
| ENSMUSG00000020473 | Aebp1 | -0.259626029 | 0.020006867 |
| ENSMUSG00000022415 | Syngr1 | -0.25990076 | 0.034931484 |
| ENSMUSG00000025927 | Tfap2b | -0.259954915 | 0.033887414 |
| ENSMUSG00000090958 | Lrrc32 | -0.260170193 | 0.058992751 |
| ENSMUSG00000058655 | Eif4b | -0.261926353 | 0.007259395 |
| ENSMUSG00000033857 | Engase | -0.262080077 | 0.054508309 |
| ENSMUSG00000048826 | Dact2 | -0.26300647 | 0.018203679 |
| ENSMUSG00000110185 | Igip | -0.263158053 | 0.007185608 |
| ENSMUSG00000054499 | Dedd2 | -0.264765725 | 0.009925648 |
| ENSMUSG00000064065 | Ipcef1 | -0.264933611 | 0.005175896 |
| ENSMUSG00000032892 | Rangrf | -0.268162497 | 0.091123501 |
| ENSMUSG00000055980 | Irs1 | -0.271732557 | 0.000427458 |
| ENSMUSG00000041075 | Fzd7 | -0.276636696 | 0.002684755 |
| ENSMUSG00000098557 | Kctd12 | -0.277957253 | 0.00566107 |
| ENSMUSG00000061143 | Maml3 | -0.278084046 | 0.003778971 |
| ENSMUSG00000010721 | Lmbr1 | -0.279063599 | 0.05980328 |
| ENSMUSG00000035413 | Tmem98 | -0.280259114 | 0.044892834 |
| ENSMUSG00000015942 | Gtf2ird2 | -0.28099444 | 0.027465928 |
| ENSMUSG00000025197 | Cyp2c23 | -0.284319963 | 0.0937822 |
| ENSMUSG00000020491 | 2810021J22Rik | -0.286943454 | 0.088978657 |
| ENSMUSG00000062563 | Cys1 | -0.288315251 | 0.000600531 |
| ENSMUSG00000106847 | Peg13 | -0.288809693 | 0.003190605 |
| ENSMUSG00000048696 | Mex3d | -0.292292029 | 0.078056757 |
| ENSMUSG00000091811 | Inafm1 | -0.293003366 | 0.050188786 |
| ENSMUSG00000069682 | Gm10275 | -0.294020956 | 0.088348954 |
| ENSMUSG00000039086 | Ss18l1 | -0.301741362 | 0.058992751 |
| ENSMUSG00000020642 | Rnf144a | -0.303851645 | 0.041615056 |
| ENSMUSG00000034156 | Tspoap1 | -0.304479775 | 0.032198638 |
| ENSMUSG00000004105 | Angptl2 | -0.305316052 | 0.003778971 |
| ENSMUSG00000034460 | Six4 | -0.315298795 | 0.001498174 |
| ENSMUSG00000074578 | Zfas1 | -0.318428874 | 0.047944462 |
| ENSMUSG00000031169 | Porcn | -0.321436174 | 0.080034473 |
| ENSMUSG00000070348 | Ccnd1 | -0.321530809 | 0.028967033 |
| ENSMUSG00000110195 | Pde2a | -0.322970176 | 7.57E-05 |
| ENSMUSG00000048285 | Frmd6 | -0.324010552 | 0.098514727 |
| ENSMUSG00000099689 | Zfp383 | -0.326386042 | 0.051741568 |
| ENSMUSG00000027524 | Edn3 | -0.334456907 | 0.050248302 |
| ENSMUSG00000045441 | Gprin3 | -0.338945523 | 0.015590247 |
| ENSMUSG00000058056 | Palld | -0.339769343 | 0.010346206 |
| ENSMUSG00000041329 | Atp1b2 | -0.340476315 | 0.05916198 |
| ENSMUSG00000002265 | Peg3 | -0.343419797 | 0.046668605 |
| ENSMUSG00000020482 | Ccdc117 | -0.344035077 | 0.000482827 |
| ENSMUSG00000053411 | Cbx7 | -0.348504166 | 7.98E-05 |
| ENSMUSG00000015656 | Hspa8 | -0.355537125 | 0.046093823 |
| ENSMUSG00000005483 | Dnajb1 | -0.355847307 | 0.058992751 |
| ENSMUSG00000067586 | S1pr3 | -0.358209262 | 0.047218992 |
| ENSMUSG00000013921 | Clip3 | -0.359056367 | 0.079198705 |
| ENSMUSG00000047632 | Fgfbp3 | -0.36118086 | 0.0935949 |
| ENSMUSG00000097412 | 1810014B01Rik | -0.362183009 | 0.012802269 |
| ENSMUSG00000042659 | Arrdc4 | -0.363825898 | 0.000127169 |
| ENSMUSG00000033350 | Chst2 | -0.366016808 | 0.022510738 |
| ENSMUSG00000031841 | Cdh13 | -0.36681342 | 0.045217761 |
| ENSMUSG00000087150 | BC064078 | -0.369530329 | 0.050188786 |
| ENSMUSG00000045519 | Zfp560 | -0.372420545 | 0.0255989 |
| ENSMUSG00000039783 | Kmo | -0.372831514 | 0.050216875 |
| ENSMUSG00000024900 | Cpt1a | -0.38401891 | 0.043533891 |
| ENSMUSG00000020607 | Fam84a | -0.384154335 | 0.036522168 |
| ENSMUSG00000095597 | Gm6472 | -0.394617005 | 0.084552085 |
| ENSMUSG00000002346 | Slc25a42 | -0.405115455 | 0.031439104 |
| ENSMUSG00000041351 | Rap1gap | -0.405712932 | 0.095782346 |
| ENSMUSG00000039377 | Hlx | -0.40938518 | 0.090316009 |
| ENSMUSG00000086583 | Gm15500 | -0.419665644 | 0.04331772 |
| ENSMUSG00000022122 | Ednrb | -0.423059378 | 0.005175896 |
| ENSMUSG00000105692 | Gm43737 | -0.424095947 | 0.099296306 |
| ENSMUSG00000033327 | Tnxb | -0.428432503 | 0.001296797 |
| ENSMUSG00000025511 | Tspan4 | -0.436903974 | 0.000215727 |
| ENSMUSG00000068742 | Cry2 | -0.448370231 | 0.009925648 |
| ENSMUSG00000018900 | Slc22a5 | -0.448863764 | 0.062405867 |
| ENSMUSG00000038530 | Rgs4 | -0.458491524 | 0.010066584 |
| ENSMUSG00000000901 | Mmp11 | -0.460032779 | 0.022906528 |
| ENSMUSG00000042745 | Id1 | -0.461688277 | 0.076742331 |
| ENSMUSG00000074063 | Osgin1 | -0.477270317 | 0.040296576 |
| ENSMUSG00000017897 | Eya2 | -0.479431769 | 0.027975707 |
| ENSMUSG00000040584 | Abcb1a | -0.518350655 | 0.009925648 |
| ENSMUSG00000037621 | Atoh8 | -0.520937333 | 0.011524868 |
| ENSMUSG00000031167 | Rbm3 | -0.533715806 | 6.77E-07 |
| ENSMUSG00000049241 | Hcar1 | -0.546319186 | 0.009234523 |
| ENSMUSG00000061353 | Cxcl12 | -0.548394671 | 0.003778971 |
| ENSMUSG00000039315 | Clnk | -0.549137451 | 0.078056757 |
| ENSMUSG00000110353 | Gm33543 | -0.558775892 | 0.060105065 |
| ENSMUSG00000060419 | Rps16-ps2 | -0.560638587 | 0.013455723 |
| ENSMUSG00000045382 | Cxcr4 | -0.579260908 | 0.002658016 |
| ENSMUSG00000037465 | Klf10 | -0.581763409 | 0.021621234 |
| ENSMUSG00000040740 | Slc25a34 | -0.58276155 | 0.0337821 |
| ENSMUSG00000027314 | Dll4 | -0.586214002 | 1.99E-05 |
| ENSMUSG00000025815 | Dhtkd1 | -0.590494107 | 7.57E-05 |
| ENSMUSG00000043144 | Aqp6 | -0.604648239 | 0.038925474 |
| ENSMUSG00000003477 | Inmt | -0.620265139 | 0.005733463 |
| ENSMUSG00000053964 | Lgals4 | -0.640082789 | 0.010346206 |
| ENSMUSG00000106062 | Gm43820 | -0.641612024 | 0.080034473 |
| ENSMUSG00000020427 | Igfbp3 | -0.644595584 | 0.03384048 |
| ENSMUSG00000022389 | Tef | -0.649107454 | 3.36E-06 |
| ENSMUSG00000028970 | Abcb1b | -0.652572336 | 0.05884925 |
| ENSMUSG00000023279 | Bmp15 | -0.660443954 | 0.06874662 |
| ENSMUSG00000063681 | Crb1 | -0.664500469 | 0.020795761 |
| ENSMUSG00000038393 | Txnip | -0.680029359 | 8.33E-05 |
| ENSMUSG00000022949 | Clic6 | -0.721720118 | 0.010346206 |
| ENSMUSG00000027796 | Smad9 | -0.723960082 | 0.013373092 |
| ENSMUSG00000050100 | Hmx2 | -0.726004638 | 0.000126048 |
| ENSMUSG00000020889 | Nr1d1 | -0.741276506 | 2.73E-06 |
| ENSMUSG00000021508 | Cxcl14 | -0.750414984 | 0.004160517 |
| ENSMUSG00000002266 | Zim1 | -0.797953279 | 0.003088823 |
| ENSMUSG00000055866 | Per2 | -0.80981235 | 0.000781449 |
| ENSMUSG00000020653 | Klf11 | -0.813883037 | 0.001117469 |
| ENSMUSG00000007872 | Id3 | -0.818955667 | 2.03E-09 |
| ENSMUSG00000021775 | Nr1d2 | -0.842607909 | 2.25E-08 |
| ENSMUSG00000030256 | Bhlhe41 | -0.978924145 | 5.95E-05 |
| ENSMUSG00000070280 | Slc22a14 | -1.104030132 | 0.057470148 |
| ENSMUSG00000015957 | Wnt11 | -1.150958375 | 0.041615056 |
| ENSMUSG00000028957 | Per3 | -1.225849132 | 7.50E-08 |
| ENSMUSG00000027875 | Hmgcs2 | -1.680238936 | 0.001365827 |
| ENSMUSG00000034450 | Gulo | -1.884334243 | 2.79E-05 |
| ENSMUSG00000059824 | Dbp | -2.25897742 | 1.69E-05 |
| ENSMUSG00000109941 | Exosc6 | -3.177140268 | 0.04845807 |

Supplementary Table 2 BALB/c (RR-3 mission) differential gene expression analysis

| gene_id | gene_name | log2FoldChange | padj |
| --- | --- | --- | --- |
| ENSMUSG00000078238 | Gm12854 | 2.372936154 | 0.053604362 |
| ENSMUSG00000085927 | Gm15888 | 1.835958833 | 0.024151443 |
| ENSMUSG00000021250 | Fos | 1.603624278 | 0.002332043 |
| ENSMUSG00000038418 | Egr1 | 1.590728739 | 0.000601048 |
| ENSMUSG00000083558 | Gm12896 | 1.521183948 | 0.021157298 |
| ENSMUSG00000037990 | Sh3rf3 | 1.470202469 | 0.037485154 |
| ENSMUSG00000047586 | Nccrp1 | 1.390722342 | 0.001281054 |
| ENSMUSG00000018822 | Sfrp5 | 1.379166221 | 0.043004949 |
| ENSMUSG00000118124 | AC129217.1 | 1.360066529 | 0.043690857 |
| ENSMUSG00000083287 | Gm13502 | 1.308417413 | 0.00211623 |
| ENSMUSG00000112057 | Gm21321 | 1.304382539 | 0.05796034 |
| ENSMUSG00000113425 | Gm48653 | 1.295687474 | 0.003493291 |
| ENSMUSG00000108597 | Gm44708 | 1.277103808 | 0.010543293 |
| ENSMUSG00000047904 | Sstr2 | 1.271092196 | 0.054519251 |
| ENSMUSG00000048782 | Insc | 1.255114353 | 0.027305016 |
| ENSMUSG00000105659 | 4930509H03Rik | 1.090312191 | 0.021710253 |
| ENSMUSG00000029608 | Rph3a | 1.082037918 | 0.094758467 |
| ENSMUSG00000049892 | Rasd1 | 1.049856471 | 0.024041459 |
| ENSMUSG00000112808 | Gm4739 | 1.036792137 | 0.074460323 |
| ENSMUSG00000022180 | Slc7a8 | 1.00546963 | 0.001673395 |
| ENSMUSG00000096980 | Gm26526 | 0.995349236 | 0.033640061 |
| ENSMUSG00000047986 | Palm3 | 0.993904097 | 0.011144912 |
| ENSMUSG00000105572 | Gm43300 | 0.966319879 | 0.019961525 |
| ENSMUSG00000097088 | Gm26615 | 0.953350895 | 0.021710253 |
| ENSMUSG00000109927 | Gm45264 | 0.926307281 | 0.071965528 |
| ENSMUSG00000030669 | Calca | 0.893718726 | 0.030141693 |
| ENSMUSG00000115190 | Gm49542 | 0.885699423 | 0.050072317 |
| ENSMUSG00000023279 | Bmp15 | 0.882217816 | 0.012690406 |
| ENSMUSG00000085781 | Gm15640 | 0.874911468 | 0.022355337 |
| ENSMUSG00000106288 | Gm43593 | 0.86588307 | 0.076793379 |
| ENSMUSG00000059852 | Kcng2 | 0.86435958 | 0.011778743 |
| ENSMUSG00000114780 | AI197445 | 0.85061953 | 0.02045526 |
| ENSMUSG00000087006 | Gm13889 | 0.840993433 | 0.002282748 |
| ENSMUSG00000027230 | Creb3l1 | 0.837763054 | 0.013904092 |
| ENSMUSG00000000325 | Arvcf | 0.837441212 | 0.001523148 |
| ENSMUSG00000031849 | Comp | 0.830934193 | 0.047883665 |
| ENSMUSG00000112858 | Rnf212b | 0.830025928 | 0.016158493 |
| ENSMUSG00000035910 | Dcdc2a | 0.827105528 | 0.068789096 |
| ENSMUSG00000097367 | Gm26718 | 0.822731881 | 0.081109671 |
| ENSMUSG00000037157 | Il22ra1 | 0.821090131 | 0.096358952 |
| ENSMUSG00000073485 | H3f3aos | 0.815299402 | 0.034219201 |
| ENSMUSG00000074217 | 2210011C24Rik | 0.794087896 | 0.02335584 |
| ENSMUSG00000098221 | Gm27030 | 0.792067635 | 0.065177077 |
| ENSMUSG00000034919 | Ttc22 | 0.791221688 | 0.007013082 |
| ENSMUSG00000053560 | Ier2 | 0.783808465 | 0.002536543 |
| ENSMUSG00000075534 | Gm13262 | 0.77995148 | 0.003604052 |
| ENSMUSG00000048583 | Igf2 | 0.771090597 | 0.008211392 |
| ENSMUSG00000029093 | Sorcs2 | 0.769863851 | 0.008577138 |
| ENSMUSG00000115248 | Gm49037 | 0.765958805 | 0.041509748 |
| ENSMUSG00000064453 | Snord21 | 0.76572205 | 0.078199087 |
| ENSMUSG00000065226 | Gm25791 | 0.764058405 | 0.070431399 |
| ENSMUSG00000060985 | Tdrd5 | 0.755514824 | 0.005073497 |
| ENSMUSG00000034212 | Ankmy1 | 0.745261851 | 0.096112274 |
| ENSMUSG00000104026 | Gm37212 | 0.737102824 | 0.084155292 |
| ENSMUSG00000117333 | AC154486.3 | 0.731258971 | 0.057615729 |
| ENSMUSG00000074580 | 4931440P22Rik | 0.720093953 | 0.070490248 |
| ENSMUSG00000006711 | D130043K22Rik | 0.717327985 | 0.065321247 |
| ENSMUSG00000114172 | Gm36298 | 0.713316586 | 0.023334737 |
| ENSMUSG00000022438 | Parvb | 0.712495405 | 0.003556942 |
| ENSMUSG00000086706 | Gm15848 | 0.711878903 | 0.025449976 |
| ENSMUSG00000098943 | Rnu3b1 | 0.709897292 | 0.058104748 |
| ENSMUSG00000048899 | Rimkla | 0.707774755 | 0.023590249 |
| ENSMUSG00000065947 | mt-Nd4l | 0.702655344 | 0.064625186 |
| ENSMUSG00000068876 | Cgn | 0.697407911 | 0.016266259 |
| ENSMUSG00000037977 | 6430571L13Rik | 0.695873441 | 0.07213013 |
| ENSMUSG00000002250 | Ppard | 0.691233198 | 0.058104748 |
| ENSMUSG00000042487 | Leo1 | 0.688860079 | 0.014258982 |
| ENSMUSG00000064337 | mt-Rnr1 | 0.679835518 | 0.08313012 |
| ENSMUSG00000053613 | Notumos | 0.678945662 | 0.047641488 |
| ENSMUSG00000042115 | Klhdc8a | 0.673630626 | 0.010270516 |
| ENSMUSG00000030825 | Hsd17b14 | 0.67087399 | 0.031163478 |
| ENSMUSG00000028834 | Trim63 | 0.6698942 | 0.09138358 |
| ENSMUSG00000027996 | Sfrp2 | 0.668723869 | 0.009710687 |
| ENSMUSG00000038677 | Scube3 | 0.665766194 | 0.002950698 |
| ENSMUSG00000034456 | Uroc1 | 0.665128091 | 0.008402735 |
| ENSMUSG00000005718 | Tfap4 | 0.657357303 | 0.077065877 |
| ENSMUSG00000020679 | Hnf1b | 0.651827774 | 0.030383446 |
| ENSMUSG00000034159 | 2310007B03Rik | 0.644777215 | 0.000643771 |
| ENSMUSG00000033107 | Rnf125 | 0.644546685 | 0.049301867 |
| ENSMUSG00000024011 | Pi16 | 0.639814535 | 0.0000448 |
| ENSMUSG00000031790 | Mmp15 | 0.639573331 | 0.014255911 |
| ENSMUSG00000030319 | Cand2 | 0.638579318 | 0.029760082 |
| ENSMUSG00000053175 | Bcl3 | 0.638244695 | 0.098213007 |
| ENSMUSG00000096847 | Tmem151b | 0.636038273 | 0.031890817 |
| ENSMUSG00000110393 | Gm36445 | 0.632721083 | 0.074247933 |
| ENSMUSG00000096870 | Gm21816 | 0.627187479 | 0.068924287 |
| ENSMUSG00000045094 | Arhgef37 | 0.623146143 | 0.034593459 |
| ENSMUSG00000055745 | Rtl6 | 0.621428425 | 0.078199087 |
| ENSMUSG00000045136 | Tubb2b | 0.621149829 | 0.014255911 |
| ENSMUSG00000080242 | Atp6v0c-ps2 | 0.601111807 | 0.027874702 |
| ENSMUSG00000107182 | Gm43268 | 0.59550614 | 0.060648268 |
| ENSMUSG00000022580 | Rhpn1 | 0.59305661 | 0.012483782 |
| ENSMUSG00000045215 | Asxl3 | 0.590983995 | 0.081479245 |
| ENSMUSG00000032523 | Hhatl | 0.58935073 | 0.021710253 |
| ENSMUSG00000039328 | Rnf122 | 0.588754484 | 0.005047458 |
| ENSMUSG00000026602 | Nphs2 | 0.582550069 | 0.007150945 |
| ENSMUSG00000103432 | 6720464F23Rik | 0.582286773 | 0.08447515 |
| ENSMUSG00000020121 | Srgap1 | 0.581138401 | 0.015853338 |
| ENSMUSG00000040964 | Arhgef10l | 0.579739784 | 0.036527701 |
| ENSMUSG00000028919 | Arhgef19 | 0.577753336 | 0.078971974 |
| ENSMUSG00000013483 | Card14 | 0.57760011 | 0.089382402 |
| ENSMUSG00000034926 | Dhcr24 | 0.572084859 | 0.067451936 |
| ENSMUSG00000059743 | Fdps | 0.568925847 | 0.007155914 |
| ENSMUSG00000015944 | Castor2 | 0.567788453 | 0.021714667 |
| ENSMUSG00000067158 | Col4a4 | 0.567077746 | 0.00211623 |
| ENSMUSG00000066026 | Dhrs3 | 0.559098044 | 0.017916465 |
| ENSMUSG00000036106 | Prr5 | 0.558451725 | 0.064821181 |
| ENSMUSG00000055633 | Zfp580 | 0.558094578 | 0.092969038 |
| ENSMUSG00000116262 | Gm49544 | 0.556206632 | 0.071965528 |
| ENSMUSG00000036957 | Lrfn3 | 0.554525094 | 0.010270516 |
| ENSMUSG00000097724 | Gm26850 | 0.554231217 | 0.029428938 |
| ENSMUSG00000022861 | Dgkg | 0.546249234 | 0.094256766 |
| ENSMUSG00000027858 | Tspan2 | 0.543290897 | 0.00045677 |
| ENSMUSG00000024727 | Trpm6 | 0.54272663 | 0.004742133 |
| ENSMUSG00000024165 | Jpt2 | 0.542701374 | 0.019083147 |
| ENSMUSG00000044647 | Csrnp3 | 0.541056752 | 0.091953909 |
| ENSMUSG00000026971 | Itgb6 | 0.539654963 | 0.041509748 |
| ENSMUSG00000086054 | Hnf1aos1 | 0.538679298 | 0.033532406 |
| ENSMUSG00000044646 | Zbtb7c | 0.538527273 | 0.058594752 |
| ENSMUSG00000019856 | Fam184a | 0.532072702 | 0.086281099 |
| ENSMUSG00000098022 | Zfp82 | 0.530567644 | 0.060648268 |
| ENSMUSG00000037664 | Cdkn1c | 0.529750857 | 0.000896946 |
| ENSMUSG00000034613 | Ppm1h | 0.527834206 | 0.017663287 |
| ENSMUSG00000041552 | Ptchd1 | 0.525342927 | 0.088964428 |
| ENSMUSG00000034371 | Tkfc | 0.518819479 | 0.079908314 |
| ENSMUSG00000046159 | Chrm3 | 0.51128953 | 0.03078539 |
| ENSMUSG00000018678 | Sp2 | 0.509197996 | 0.050072317 |
| ENSMUSG00000037005 | Xpnpep2 | 0.508933573 | 0.019751377 |
| ENSMUSG00000045294 | Insig1 | 0.507947511 | 0.089169591 |
| ENSMUSG00000008348 | Ubc | 0.502720481 | 0.057615729 |
| ENSMUSG00000087221 | BC037032 | 0.502272236 | 0.081220422 |
| ENSMUSG00000024736 | Tmem132a | 0.497369636 | 0.018253826 |
| ENSMUSG00000034714 | Ttyh2 | 0.497141472 | 0.067827109 |
| ENSMUSG00000054074 | Skida1 | 0.493681494 | 0.062727442 |
| ENSMUSG00000018339 | Gpx3 | 0.487204855 | 0.09565271 |
| ENSMUSG00000031891 | Hsd11b2 | 0.485607515 | 0.058019934 |
| ENSMUSG00000097729 | 2310015A10Rik | 0.483822071 | 0.094090847 |
| ENSMUSG00000066438 | Plekhd1 | 0.483067222 | 0.077278809 |
| ENSMUSG00000091243 | Vgll3 | 0.481979737 | 0.042814969 |
| ENSMUSG00000005125 | Ndrg1 | 0.478617818 | 0.088048381 |
| ENSMUSG00000051169 | Rpusd3 | 0.469609325 | 0.081479245 |
| ENSMUSG00000056413 | Adap1 | 0.463851635 | 0.093921685 |
| ENSMUSG00000007817 | Zmiz1 | 0.461758368 | 0.078835393 |
| ENSMUSG00000036095 | Dgkb | 0.461441802 | 0.052639602 |
| ENSMUSG00000058454 | Dhcr7 | 0.460103289 | 0.014508536 |
| ENSMUSG00000026463 | Atp2b4 | 0.460073647 | 0.002584031 |
| ENSMUSG00000041119 | Pde9a | 0.45997891 | 0.099363633 |
| ENSMUSG00000022037 | Clu | 0.457905444 | 0.016837615 |
| ENSMUSG00000086390 | 1810019D21Rik | 0.456444867 | 0.066578533 |
| ENSMUSG00000000416 | Cttnbp2 | 0.454855698 | 0.002139858 |
| ENSMUSG00000032332 | Col12a1 | 0.453768711 | 0.070916159 |
| ENSMUSG00000063903 | Klk1 | 0.449583043 | 0.018930876 |
| ENSMUSG00000044566 | Cage1 | 0.449110795 | 0.01221796 |
| ENSMUSG00000051495 | Irf2bp2 | 0.443613834 | 0.010348347 |
| ENSMUSG00000061046 | Haghl | 0.442417101 | 0.007155914 |
| ENSMUSG00000018411 | Mapt | 0.441794469 | 0.024121428 |
| ENSMUSG00000025902 | Sox17 | 0.440491223 | 0.03078539 |
| ENSMUSG00000036390 | Gadd45a | 0.439511366 | 0.015126351 |
| ENSMUSG00000027827 | Kcnab1 | 0.433101493 | 0.08313012 |
| ENSMUSG00000078963 | Hsbp1l1 | 0.428215728 | 0.08241924 |
| ENSMUSG00000071856 | Mcc | 0.420991359 | 0.004232819 |
| ENSMUSG00000052595 | A1cf | 0.419989609 | 0.085898767 |
| ENSMUSG00000107881 | Gm44250 | 0.419507928 | 0.004765203 |
| ENSMUSG00000078868 | Gm14412 | 0.418890091 | 0.073511247 |
| ENSMUSG00000085795 | Zfp703 | 0.417963591 | 0.070674228 |
| ENSMUSG00000028059 | Arhgef2 | 0.416565705 | 0.023005631 |
| ENSMUSG00000000093 | Tbx2 | 0.408803287 | 0.048906298 |
| ENSMUSG00000028031 | Dkk2 | 0.403050842 | 0.063128996 |
| ENSMUSG00000029580 | Actb | 0.402005477 | 0.097807265 |
| ENSMUSG00000096054 | Syne1 | 0.401762607 | 0.03185539 |
| ENSMUSG00000054385 | Ceacam2 | 0.400784414 | 0.059493218 |
| ENSMUSG00000016458 | Wt1 | 0.40051561 | 0.061680137 |
| ENSMUSG00000021750 | Fam107a | 0.398979144 | 0.014839713 |
| ENSMUSG00000099966 | 2810402E24Rik | 0.398590809 | 0.03185539 |
| ENSMUSG00000034330 | Plcg2 | 0.398540892 | 0.063953795 |
| ENSMUSG00000074622 | Mafb | 0.398068153 | 0.066460829 |
| ENSMUSG00000031349 | Nsdhl | 0.397761261 | 0.056250027 |
| ENSMUSG00000066357 | Wdr6 | 0.393913645 | 0.064625186 |
| ENSMUSG00000020340 | Cyfip2 | 0.391382067 | 0.064821181 |
| ENSMUSG00000032625 | Thsd7a | 0.39079489 | 0.034853127 |
| ENSMUSG00000078706 | Gm53 | 0.389627289 | 0.099245056 |
| ENSMUSG00000038331 | Satb2 | 0.388995166 | 0.024198723 |
| ENSMUSG00000035275 | Raver2 | 0.388048873 | 0.074460323 |
| ENSMUSG00000070883 | Ccdc173 | 0.38164332 | 0.054203996 |
| ENSMUSG00000048406 | B330016D10Rik | 0.381246396 | 0.059472089 |
| ENSMUSG00000009566 | Fpgs | 0.380516787 | 0.074050556 |
| ENSMUSG00000020741 | Cluh | 0.380511894 | 0.089739548 |
| ENSMUSG00000021559 | Dapk1 | 0.380283864 | 0.026482301 |
| ENSMUSG00000036473 | Tbc1d24 | 0.378621898 | 0.023334737 |
| ENSMUSG00000031520 | Vegfc | 0.377971802 | 0.070018069 |
| ENSMUSG00000025196 | Cpn1 | 0.374064469 | 0.081479245 |
| ENSMUSG00000019647 | Sema6a | 0.373118105 | 0.06893765 |
| ENSMUSG00000108350 | Gm44950 | 0.371517054 | 0.037543358 |
| ENSMUSG00000017639 | Rab11fip4 | 0.370841288 | 0.014326612 |
| ENSMUSG00000017677 | Wsb1 | 0.365951246 | 0.09495426 |
| ENSMUSG00000046079 | Lrrc8d | 0.358408371 | 0.09856681 |
| ENSMUSG00000024565 | Sall3 | 0.353843734 | 0.084155292 |
| ENSMUSG00000026255 | Efhd1 | 0.3513937 | 0.00689822 |
| ENSMUSG00000025885 | Myo5b | 0.345810893 | 0.015213621 |
| ENSMUSG00000019478 | Rab4a | 0.343878901 | 0.031567938 |
| ENSMUSG00000026500 | Cox20 | 0.342753631 | 0.042161245 |
| ENSMUSG00000052632 | Asap2 | 0.338983746 | 0.020093552 |
| ENSMUSG00000069892 | 9930111J21Rik2 | 0.336576971 | 0.070452731 |
| ENSMUSG00000036617 | Etl4 | 0.33538858 | 0.034973243 |
| ENSMUSG00000085037 | 4933421O10Rik | 0.332205427 | 0.095442347 |
| ENSMUSG00000020329 | Polrmt | 0.328794835 | 0.068866925 |
| ENSMUSG00000025212 | Sfxn3 | 0.327816126 | 0.05796034 |
| ENSMUSG00000033389 | Arhgap44 | 0.326117979 | 0.020914045 |
| ENSMUSG00000022763 | Aifm3 | 0.324954964 | 0.086855887 |
| ENSMUSG00000024236 | Svil | 0.324752071 | 0.027128998 |
| ENSMUSG00000079469 | Pigb | 0.324701645 | 0.062474984 |
| ENSMUSG00000034813 | Grip1 | 0.323934374 | 0.091777304 |
| ENSMUSG00000058833 | Rex1bd | 0.321871409 | 0.074247933 |
| ENSMUSG00000031548 | Sfrp1 | 0.321365509 | 0.012668928 |
| ENSMUSG00000025092 | Hspa12a | 0.320930576 | 0.026678457 |
| ENSMUSG00000041895 | Wipi1 | 0.320552462 | 0.073100154 |
| ENSMUSG00000035392 | Dennd1a | 0.319791629 | 0.058680095 |
| ENSMUSG00000073910 | Mob3b | 0.319650135 | 0.019961525 |
| ENSMUSG00000084128 | Esrp2 | 0.319637118 | 0.041509748 |
| ENSMUSG00000061751 | Kalrn | 0.318321121 | 0.012668928 |
| ENSMUSG00000029101 | Rgs12 | 0.317061825 | 0.073170916 |
| ENSMUSG00000032718 | Mansc1 | 0.315834627 | 0.021710253 |
| ENSMUSG00000028917 | Plekhm2 | 0.313352472 | 0.08154222 |
| ENSMUSG00000031881 | Cdh16 | 0.310274157 | 0.021710253 |
| ENSMUSG00000007721 | Ccdc124 | 0.306667992 | 0.091269869 |
| ENSMUSG00000063894 | Zkscan8 | 0.305645794 | 0.038157715 |
| ENSMUSG00000042063 | Zfp386 | 0.303902094 | 0.047095276 |
| ENSMUSG00000027210 | Meis2 | 0.302476576 | 0.074489848 |
| ENSMUSG00000005299 | Letm1 | 0.302397638 | 0.024166593 |
| ENSMUSG00000020642 | Rnf144a | 0.302017051 | 0.074344692 |
| ENSMUSG00000020462 | Cfap36 | 0.301403324 | 0.052646481 |
| ENSMUSG00000073755 | 5730409E04Rik | 0.301150209 | 0.064317008 |
| ENSMUSG00000028238 | Atp6v0d2 | 0.29747946 | 0.03291281 |
| ENSMUSG00000034427 | Myo15b | 0.29547525 | 0.09733203 |
| ENSMUSG00000020744 | Slc25a19 | 0.293883726 | 0.037485154 |
| ENSMUSG00000045691 | Thtpa | 0.293539124 | 0.060648268 |
| ENSMUSG00000025142 | Aspscr1 | 0.293069736 | 0.011293339 |
| ENSMUSG00000090626 | Tex9 | 0.292991136 | 0.048799108 |
| ENSMUSG00000029032 | Arhgef16 | 0.292869622 | 0.069863082 |
| ENSMUSG00000057315 | Arhgap24 | 0.286678977 | 0.086281099 |
| ENSMUSG00000058690 | Ccser2 | 0.285720268 | 0.065437648 |
| ENSMUSG00000000216 | Scnn1g | 0.282150525 | 0.091777304 |
| ENSMUSG00000041134 | Cyyr1 | 0.273114943 | 0.098213007 |
| ENSMUSG00000031227 | Magee1 | 0.272961103 | 0.052646481 |
| ENSMUSG00000032463 | Faim | 0.272050211 | 0.048483854 |
| ENSMUSG00000027523 | Gnas | 0.268827714 | 0.098213007 |
| ENSMUSG00000034109 | Golim4 | 0.266890635 | 0.099025802 |
| ENSMUSG00000026566 | Mpzl1 | 0.263002993 | 0.068924287 |
| ENSMUSG00000071042 | Rasgrp3 | 0.261739403 | 0.084155292 |
| ENSMUSG00000037366 | Pafah2 | 0.261484367 | 0.042814969 |
| ENSMUSG00000065979 | Cpped1 | 0.260695775 | 0.012929757 |
| ENSMUSG00000044712 | Slc38a6 | 0.255937573 | 0.071965528 |
| ENSMUSG00000031626 | Sorbs2 | 0.254242229 | 0.024041459 |
| ENSMUSG00000029729 | Zkscan1 | 0.253164284 | 0.014485884 |
| ENSMUSG00000024269 | Tpgs2 | 0.251191409 | 0.064462059 |
| ENSMUSG00000024169 | Ift140 | 0.251061313 | 0.053604362 |
| ENSMUSG00000044393 | Dsg2 | 0.247957118 | 0.064269001 |
| ENSMUSG00000063410 | Stk24 | 0.24246917 | 0.056875915 |
| ENSMUSG00000032018 | Sc5d | 0.240874055 | 0.046890487 |
| ENSMUSG00000024104 | Washc2 | 0.2385943 | 0.050746761 |
| ENSMUSG00000035954 | Dock4 | 0.237522623 | 0.094256766 |
| ENSMUSG00000027695 | Pld1 | 0.232593343 | 0.094090847 |
| ENSMUSG00000032249 | Anp32a | 0.227352869 | 0.008791593 |
| ENSMUSG00000026672 | Optn | 0.225812097 | 0.040106056 |
| ENSMUSG00000014426 | Map3k4 | 0.223244313 | 0.060648268 |
| ENSMUSG00000032352 | Lrrc1 | 0.220680558 | 0.070273435 |
| ENSMUSG00000041777 | Cir1 | 0.219806193 | 0.092271549 |
| ENSMUSG00000040820 | Hlcs | 0.2159867 | 0.05689402 |
| ENSMUSG00000019971 | Cep290 | 0.214967734 | 0.012022604 |
| ENSMUSG00000026490 | Cdc42bpa | 0.214886173 | 0.058104748 |
| ENSMUSG00000020413 | Hus1 | 0.207048912 | 0.057631801 |
| ENSMUSG00000059923 | Grb2 | 0.201942807 | 0.09036289 |
| ENSMUSG00000027374 | Mrps5 | 0.198386056 | 0.061906057 |
| ENSMUSG00000001768 | Rin2 | 0.196873771 | 0.081202536 |
| ENSMUSG00000037697 | Ddhd1 | 0.196047742 | 0.091269869 |
| ENSMUSG00000102976 | Zc3h11a | 0.190277727 | 0.380214286 |
| ENSMUSG00000025246 | Tbl1x | 0.177331823 | 0.081367583 |
| ENSMUSG00000057329 | Bcl2 | 0.173650296 | 0.049704565 |
| ENSMUSG00000034252 | Senp6 | 0.149311221 | 0.066200932 |
| ENSMUSG00000035949 | Fbxw2 | -0.125914163 | 0.090582483 |
| ENSMUSG00000035284 | Vps13c | -0.140508897 | 0.071965528 |
| ENSMUSG00000025078 | Nhlrc2 | -0.148575149 | 0.064400501 |
| ENSMUSG00000026848 | Tor1b | -0.152372061 | 0.022076678 |
| ENSMUSG00000024122 | Pdpk1 | -0.156913575 | 0.08913105 |
| ENSMUSG00000022781 | Pak2 | -0.157162067 | 0.047883665 |
| ENSMUSG00000036323 | Srp72 | -0.164381074 | 0.074876886 |
| ENSMUSG00000029110 | Rnf4 | -0.164606401 | 0.008797476 |
| ENSMUSG00000036646 | Man1b1 | -0.179108813 | 0.059493218 |
| ENSMUSG00000030967 | Zranb1 | -0.180902969 | 0.067827109 |
| ENSMUSG00000027668 | Mfn1 | -0.184908562 | 0.012022604 |
| ENSMUSG00000048000 | Gigyf2 | -0.18868078 | 0.056918539 |
| ENSMUSG00000015776 | Med22 | -0.190008668 | 0.081479245 |
| ENSMUSG00000037260 | Hgsnat | -0.190335916 | 0.063175216 |
| ENSMUSG00000022066 | Entpd4b | -0.19060507 | 0.074460323 |
| ENSMUSG00000095463 | Entpd4 | -0.19060507 | 0.074460323 |
| ENSMUSG00000026608 | Kctd3 | -0.192523769 | 0.03078539 |
| ENSMUSG00000006494 | Pdk1 | -0.194333823 | 0.076515098 |
| ENSMUSG00000047539 | Fbxo28 | -0.198326078 | 0.010586335 |
| ENSMUSG00000027865 | Gdap2 | -0.198894249 | 0.058218262 |
| ENSMUSG00000054843 | Atrnl1 | -0.199745012 | 0.02271515 |
| ENSMUSG00000021149 | Gtpbp4 | -0.20261701 | 0.012668928 |
| ENSMUSG00000014355 | Anapc1 | -0.204736959 | 0.066746255 |
| ENSMUSG00000031729 | Ist1 | -0.208355942 | 0.024041459 |
| ENSMUSG00000034320 | Slc26a2 | -0.208620853 | 0.027128998 |
| ENSMUSG00000029924 | Slc37a3 | -0.21306323 | 0.08379929 |
| ENSMUSG00000038619 | Ensa | -0.213308436 | 0.047266699 |
| ENSMUSG00000026577 | Blzf1 | -0.215172432 | 0.023410941 |
| ENSMUSG00000026289 | Atg16l1 | -0.21854706 | 0.074489848 |
| ENSMUSG00000028878 | Fam76a | -0.221081231 | 0.081367583 |
| ENSMUSG00000012535 | Tnpo3 | -0.221204039 | 0.03213683 |
| ENSMUSG00000021385 | Ippk | -0.22211939 | 0.088154848 |
| ENSMUSG00000017561 | Crlf3 | -0.222231916 | 0.091953909 |
| ENSMUSG00000020458 | Rtn4 | -0.222232861 | 0.001688368 |
| ENSMUSG00000020248 | Nfyb | -0.22367438 | 0.064087674 |
| ENSMUSG00000022555 | Dgat1 | -0.22395029 | 0.062474984 |
| ENSMUSG00000093752 | Gm20716 | -0.224546003 | 0.091006867 |
| ENSMUSG00000034893 | Cog3 | -0.22471981 | 0.08727355 |
| ENSMUSG00000031918 | Mtmr2 | -0.225395024 | 0.048933785 |
| ENSMUSG00000021518 | Ptdss1 | -0.226160076 | 0.066746255 |
| ENSMUSG00000040599 | Mis12 | -0.226965954 | 0.053604362 |
| ENSMUSG00000020954 | Strn3 | -0.227231313 | 0.074489848 |
| ENSMUSG00000004558 | Ndrg2 | -0.230907545 | 0.096459848 |
| ENSMUSG00000023963 | Cyp39a1 | -0.230997309 | 0.08913105 |
| ENSMUSG00000040325 | Dcaf1 | -0.232416099 | 0.090103031 |
| ENSMUSG00000042472 | Zfp410 | -0.235280551 | 0.032237107 |
| ENSMUSG00000041769 | Ppp2r2d | -0.237240541 | 0.094256766 |
| ENSMUSG00000069495 | Epc2 | -0.241751442 | 0.044678142 |
| ENSMUSG00000027630 | Tbl1xr1 | -0.242055817 | 0.091794323 |
| ENSMUSG00000041797 | Abca9 | -0.242449731 | 0.091009147 |
| ENSMUSG00000027804 | Ppid | -0.243040758 | 0.048483854 |
| ENSMUSG00000035620 | Ric8b | -0.243099107 | 0.055630601 |
| ENSMUSG00000031156 | Slc35a2 | -0.243665409 | 0.070273435 |
| ENSMUSG00000038506 | Dcun1d2 | -0.246292572 | 0.077354625 |
| ENSMUSG00000032437 | Stt3b | -0.246590581 | 0.094910277 |
| ENSMUSG00000026457 | Adipor1 | -0.24780076 | 0.070746278 |
| ENSMUSG00000031358 | Msl3 | -0.247910292 | 0.085413687 |
| ENSMUSG00000071359 | Tbpl1 | -0.249910933 | 0.019840593 |
| ENSMUSG00000030275 | Etnk1 | -0.250515865 | 0.09036289 |
| ENSMUSG00000021830 | Txndc16 | -0.250754944 | 0.084155292 |
| ENSMUSG00000020257 | Wdr82 | -0.251506129 | 0.056875915 |
| ENSMUSG00000067787 | Blcap | -0.251741573 | 0.048799108 |
| ENSMUSG00000015961 | Adss | -0.252447899 | 0.045942643 |
| ENSMUSG00000026238 | Ptma | -0.25249878 | 0.006458583 |
| ENSMUSG00000041712 | Ubr7 | -0.252810757 | 0.03078539 |
| ENSMUSG00000100164 | 2610306M01Rik | -0.254073371 | 0.099556851 |
| ENSMUSG00000021771 | Vdac2 | -0.254337567 | 0.016500344 |
| ENSMUSG00000017802 | Retreg3 | -0.255047426 | 0.008675656 |
| ENSMUSG00000036890 | Gtdc1 | -0.256357027 | 0.042312237 |
| ENSMUSG00000022828 | Gtf2e1 | -0.259360339 | 0.097792106 |
| ENSMUSG00000043131 | Mob1a | -0.26295328 | 0.032396748 |
| ENSMUSG00000028899 | Taf12 | -0.26427541 | 0.094824675 |
| ENSMUSG00000006599 | Gtf2h1 | -0.264539999 | 0.034036016 |
| ENSMUSG00000026698 | Pigc | -0.265117433 | 0.050006794 |
| ENSMUSG00000026335 | Pam | -0.26524819 | 0.09093847 |
| ENSMUSG00000118346 | Tmem179b | -0.266307176 | 0.074247933 |
| ENSMUSG00000031568 | Rwdd4a | -0.266412718 | 0.018031218 |
| ENSMUSG00000026491 | Ahctf1 | -0.266733513 | 0.034958752 |
| ENSMUSG00000029221 | Slc30a9 | -0.267001536 | 0.080906475 |
| ENSMUSG00000028017 | Egf | -0.268273281 | 0.023169737 |
| ENSMUSG00000039725 | Trp53rka | -0.268615555 | 0.067366447 |
| ENSMUSG00000028683 | Eif2b3 | -0.26925461 | 0.079908314 |
| ENSMUSG00000059173 | Pde1a | -0.269470762 | 0.078038352 |
| ENSMUSG00000005813 | Metap1 | -0.270692428 | 0.015013555 |
| ENSMUSG00000027602 | Map1lc3a | -0.271290465 | 0.084155292 |
| ENSMUSG00000030753 | Thap12 | -0.27176775 | 0.096112274 |
| ENSMUSG00000055553 | Kxd1 | -0.273023144 | 0.060648268 |
| ENSMUSG00000019578 | Ubxn6 | -0.274852473 | 0.091777304 |
| ENSMUSG00000021258 | Ccnk | -0.277508692 | 0.005737183 |
| ENSMUSG00000002845 | Tmem39a | -0.280057837 | 0.03245408 |
| ENSMUSG00000066800 | Rnasel | -0.281850301 | 0.058419391 |
| ENSMUSG00000036442 | Thap11 | -0.28422571 | 0.008369837 |
| ENSMUSG00000019891 | Dcbld1 | -0.284388891 | 0.059035783 |
| ENSMUSG00000028420 | Tmem38b | -0.284604156 | 0.058419391 |
| ENSMUSG00000027708 | Dcun1d1 | -0.285427461 | 0.044376059 |
| ENSMUSG00000001700 | Gramd3 | -0.286117253 | 0.034791106 |
| ENSMUSG00000011960 | Ccnt1 | -0.287364545 | 0.09017974 |
| ENSMUSG00000029426 | Scarb2 | -0.290614253 | 0.042252146 |
| ENSMUSG00000041453 | Rpl21 | -0.292250099 | 0.060648268 |
| ENSMUSG00000046808 | Atp10d | -0.292358503 | 0.009451282 |
| ENSMUSG00000002325 | Irf9 | -0.293443728 | 0.047095276 |
| ENSMUSG00000022704 | Qtrt2 | -0.294066122 | 0.018607405 |
| ENSMUSG00000029534 | St7 | -0.294918167 | 0.081366408 |
| ENSMUSG00000053559 | Smagp | -0.295923189 | 0.083573874 |
| ENSMUSG00000035399 | Oser1 | -0.296283204 | 0.031440125 |
| ENSMUSG00000031558 | Slit2 | -0.296949334 | 0.071889022 |
| ENSMUSG00000020610 | Amz2 | -0.297037111 | 0.033451485 |
| ENSMUSG00000015575 | Atp6v0e | -0.298623108 | 0.055630601 |
| ENSMUSG00000022283 | Pabpc1 | -0.298953702 | 0.061468333 |
| ENSMUSG00000022969 | Il10rb | -0.301808706 | 0.085029551 |
| ENSMUSG00000050890 | Pdik1l | -0.30190789 | 0.014787574 |
| ENSMUSG00000026479 | Lamc2 | -0.302432981 | 0.009337762 |
| ENSMUSG00000020014 | Cfap54 | -0.303659257 | 0.089991605 |
| ENSMUSG00000003380 | Rabac1 | -0.308305123 | 0.077600526 |
| ENSMUSG00000014353 | Tmem87b | -0.308932793 | 0.009728111 |
| ENSMUSG00000028773 | Fabp3 | -0.309443528 | 0.078680184 |
| ENSMUSG00000038612 | Mcl1 | -0.309585389 | 0.054035991 |
| ENSMUSG00000029394 | Cdk2ap1 | -0.310129535 | 0.042814969 |
| ENSMUSG00000029998 | Pcyox1 | -0.312297679 | 0.024198723 |
| ENSMUSG00000022570 | Tsta3 | -0.31236478 | 0.058104748 |
| ENSMUSG00000028015 | Ctso | -0.312532815 | 0.09036289 |
| ENSMUSG00000049755 | Zfp672 | -0.315309087 | 0.008402735 |
| ENSMUSG00000021171 | Esyt2 | -0.316505638 | 0.003885287 |
| ENSMUSG00000019528 | Gyg | -0.317757495 | 0.062973935 |
| ENSMUSG00000051557 | Pusl1 | -0.317798817 | 0.073446532 |
| ENSMUSG00000033880 | Lgals3bp | -0.318426412 | 0.077958844 |
| ENSMUSG00000032171 | Pin1 | -0.319615084 | 0.054414776 |
| ENSMUSG00000094870 | Zfp131 | -0.320287914 | 0.074099 |
| ENSMUSG00000071551 | Akr1c19 | -0.321057108 | 0.03498095 |
| ENSMUSG00000031791 | Tmem38a | -0.322513498 | 0.014713832 |
| ENSMUSG00000006931 | P3h4 | -0.322826748 | 0.077532883 |
| ENSMUSG00000039770 | Ypel5 | -0.32483857 | 0.014326612 |
| ENSMUSG00000026869 | Psmd5 | -0.324922618 | 0.067366447 |
| ENSMUSG00000029416 | Slc15a4 | -0.325828705 | 0.036784973 |
| ENSMUSG00000001366 | Fbxo9 | -0.327313968 | 0.025825443 |
| ENSMUSG00000090137 | Uba52 | -0.327515321 | 0.059005362 |
| ENSMUSG00000004880 | Lbr | -0.328980667 | 0.047095276 |
| ENSMUSG00000035297 | Cops4 | -0.329116736 | 0.036447318 |
| ENSMUSG00000022681 | Ntan1 | -0.329212568 | 0.060648268 |
| ENSMUSG00000020063 | Sirt1 | -0.330667522 | 0.059833629 |
| ENSMUSG00000033985 | Tesk2 | -0.335058412 | 0.038563335 |
| ENSMUSG00000022684 | Bfar | -0.33570595 | 0.012090403 |
| ENSMUSG00000024245 | Tmem178 | -0.335800612 | 0.054414776 |
| ENSMUSG00000071632 | 2510002D24Rik | -0.336404046 | 0.035912126 |
| ENSMUSG00000031029 | Eif3f | -0.336545595 | 0.01841976 |
| ENSMUSG00000031209 | Heph | -0.336698072 | 0.032237107 |
| ENSMUSG00000031545 | Gpat4 | -0.338362765 | 0.054123509 |
| ENSMUSG00000025362 | Rps26 | -0.340573954 | 0.064625186 |
| ENSMUSG00000027810 | Eif2a | -0.340643297 | 0.037543358 |
| ENSMUSG00000013275 | Slc41a1 | -0.340668619 | 0.088964428 |
| ENSMUSG00000024725 | Ostf1 | -0.340949653 | 0.073519262 |
| ENSMUSG00000026088 | Mitd1 | -0.344896971 | 0.051838613 |
| ENSMUSG00000031221 | Igbp1 | -0.347514094 | 0.013269086 |
| ENSMUSG00000027180 | Fbxo3 | -0.34955069 | 0.05782848 |
| ENSMUSG00000028709 | Mob3c | -0.352574408 | 0.080942926 |
| ENSMUSG00000028669 | Pithd1 | -0.357274734 | 0.043964357 |
| ENSMUSG00000021629 | Slc30a5 | -0.358394325 | 0.076980136 |
| ENSMUSG00000038525 | Armc10 | -0.359032233 | 0.08918804 |
| ENSMUSG00000004356 | Utp20 | -0.360206268 | 0.03245408 |
| ENSMUSG00000028822 | Tmem50a | -0.361775059 | 0.022355337 |
| ENSMUSG00000031875 | Cmtm3 | -0.362139128 | 0.061906057 |
| ENSMUSG00000039990 | Edrf1 | -0.36244169 | 0.020914045 |
| ENSMUSG00000030888 | Rrp8 | -0.36312563 | 0.005590443 |
| ENSMUSG00000032112 | Trappc4 | -0.363190782 | 0.077065877 |
| ENSMUSG00000063281 | Zfp35 | -0.364487749 | 0.087851422 |
| ENSMUSG00000025049 | Taf5 | -0.364558473 | 0.03986264 |
| ENSMUSG00000017707 | Serinc3 | -0.367490451 | 0.004765203 |
| ENSMUSG00000000056 | Narf | -0.36794854 | 0.050141884 |
| ENSMUSG00000021242 | Npc2 | -0.368271223 | 0.031182829 |
| ENSMUSG00000028889 | Yrdc | -0.370146737 | 0.056153045 |
| ENSMUSG00000056116 | H2-T22 | -0.372411985 | 0.026482301 |
| ENSMUSG00000041126 | H2afv | -0.373686026 | 0.063230682 |
| ENSMUSG00000025436 | Atp23 | -0.374767527 | 0.071965528 |
| ENSMUSG00000049950 | Rpp38 | -0.375295837 | 0.032229411 |
| ENSMUSG00000066232 | Ipo7 | -0.378265136 | 0.0800206 |
| ENSMUSG00000071415 | Rpl23 | -0.380108469 | 0.049640566 |
| ENSMUSG00000014077 | Chp1 | -0.38329794 | 0.019083147 |
| ENSMUSG00000021102 | Glrx5 | -0.384029671 | 0.043048724 |
| ENSMUSG00000006941 | Eif1b | -0.384310965 | 0.054519251 |
| ENSMUSG00000022122 | Ednrb | -0.385330153 | 0.036876524 |
| ENSMUSG00000026110 | Mgat4a | -0.38602118 | 0.01221796 |
| ENSMUSG00000035642 | Aamdc | -0.386062742 | 0.090582483 |
| ENSMUSG00000062683 | Atp5g2 | -0.387921376 | 0.008577138 |
| ENSMUSG00000079491 | H2-T10 | -0.38792464 | 0.07584735 |
| ENSMUSG00000031068 | Glrx3 | -0.388085542 | 0.098213007 |
| ENSMUSG00000027598 | Itch | -0.388651893 | 0.008577138 |
| ENSMUSG00000028494 | Plin2 | -0.390270266 | 0.034036016 |
| ENSMUSG00000068039 | Tcp1 | -0.39054187 | 0.060648268 |
| ENSMUSG00000078941 | Ak6 | -0.39161755 | 0.089759436 |
| ENSMUSG00000059070 | Rpl18 | -0.3916843 | 0.029937616 |
| ENSMUSG00000047040 | Prr15l | -0.391804776 | 0.066746255 |
| ENSMUSG00000024875 | Yif1a | -0.392538821 | 0.047401287 |
| ENSMUSG00000032892 | Rangrf | -0.393770346 | 0.014713832 |
| ENSMUSG00000018761 | Mpdu1 | -0.395039889 | 0.007155914 |
| ENSMUSG00000038975 | Rabggtb | -0.396906896 | 0.006483055 |
| ENSMUSG00000063889 | Crem | -0.397466692 | 0.064400501 |
| ENSMUSG00000025724 | Sec11a | -0.397573995 | 0.026522059 |
| ENSMUSG00000021037 | Ahsa1 | -0.39761215 | 0.042252146 |
| ENSMUSG00000009630 | Ppp2cb | -0.398685802 | 0.000652182 |
| ENSMUSG00000025508 | Rplp2 | -0.400248005 | 0.073594774 |
| ENSMUSG00000039789 | Zfp597 | -0.4011161 | 0.024041459 |
| ENSMUSG00000045503 | Sys1 | -0.401437153 | 0.022844247 |
| ENSMUSG00000022014 | Epsti1 | -0.401943107 | 0.064272375 |
| ENSMUSG00000079104 | Prps1l3 | -0.404538654 | 0.016975829 |
| ENSMUSG00000023572 | Ccndbp1 | -0.404539526 | 0.008361157 |
| ENSMUSG00000025287 | Acot9 | -0.405944902 | 0.081479245 |
| ENSMUSG00000025967 | Eef1b2 | -0.407154945 | 0.056353372 |
| ENSMUSG00000019054 | Fis1 | -0.407653256 | 0.04831594 |
| ENSMUSG00000005054 | Cstb | -0.408947164 | 0.081220422 |
| ENSMUSG00000028633 | Ctps | -0.409104079 | 0.011162185 |
| ENSMUSG00000025586 | Cpeb1 | -0.410554606 | 0.075761131 |
| ENSMUSG00000030729 | Pgm2l1 | -0.411588871 | 0.066200932 |
| ENSMUSG00000031278 | Acsl4 | -0.411742678 | 0.069512026 |
| ENSMUSG00000023307 | Mar-05 | -0.412684632 | 0.022106663 |
| ENSMUSG00000050732 | Vamp8 | -0.413220173 | 0.025449976 |
| ENSMUSG00000037742 | Eef1a1 | -0.416956452 | 0.058001624 |
| ENSMUSG00000010048 | Ifrd2 | -0.42198917 | 0.073198506 |
| ENSMUSG00000079144 | A130010J15Rik | -0.422221486 | 0.043048724 |
| ENSMUSG00000028035 | Dnajb4 | -0.427741049 | 0.058019934 |
| ENSMUSG00000062353 | Gm15772 | -0.429266406 | 0.061244584 |
| ENSMUSG00000031320 | Rps4x | -0.429875797 | 0.017007313 |
| ENSMUSG00000024066 | Xdh | -0.430461489 | 0.098443746 |
| ENSMUSG00000036833 | Pnpla7 | -0.431300062 | 0.008577138 |
| ENSMUSG00000041548 | Hspb8 | -0.431413599 | 0.030141693 |
| ENSMUSG00000025069 | Gsto2 | -0.431545979 | 0.009697507 |
| ENSMUSG00000042198 | Chchd7 | -0.431819491 | 0.046962941 |
| ENSMUSG00000007739 | Cct4 | -0.432573312 | 0.01033327 |
| ENSMUSG00000015966 | Il17rb | -0.434005756 | 0.027112014 |
| ENSMUSG00000042064 | Myo3b | -0.435125726 | 0.069567902 |
| ENSMUSG00000002395 | Use1 | -0.437602841 | 0.023979949 |
| ENSMUSG00000020334 | Slc22a4 | -0.4406027 | 0.037485154 |
| ENSMUSG00000037762 | Slc16a9 | -0.443969013 | 0.089739548 |
| ENSMUSG00000084349 | Rpl3-ps1 | -0.445139533 | 0.005202796 |
| ENSMUSG00000025911 | Adhfe1 | -0.44551733 | 0.099025802 |
| ENSMUSG00000086290 | Snhg12 | -0.446167257 | 0.016880867 |
| ENSMUSG00000041841 | Rpl37 | -0.449202249 | 0.02335584 |
| ENSMUSG00000038489 | Polr2l | -0.450143243 | 0.014326612 |
| ENSMUSG00000068749 | Psma5 | -0.453411262 | 0.074876886 |
| ENSMUSG00000110353 | Gm33543 | -0.454368977 | 0.089759436 |
| ENSMUSG00000036123 | Slc9a3 | -0.456231805 | 0.054203996 |
| ENSMUSG00000017188 | Coa3 | -0.457645689 | 0.093902233 |
| ENSMUSG00000097906 | Gm9625 | -0.458550789 | 0.08202126 |
| ENSMUSG00000027170 | Eif3m | -0.459494607 | 0.091777304 |
| ENSMUSG00000038530 | Rgs4 | -0.459596074 | 0.048273769 |
| ENSMUSG00000030222 | Rerg | -0.460629399 | 0.023067252 |
| ENSMUSG00000075232 | Amd1 | -0.463428856 | 0.060648268 |
| ENSMUSG00000028495 | Rps6 | -0.466153948 | 0.012911524 |
| ENSMUSG00000024436 | Mrps18b | -0.469740594 | 0.006069071 |
| ENSMUSG00000081406 | Rps6-ps4 | -0.472699852 | 0.063230682 |
| ENSMUSG00000028936 | Rpl22 | -0.475640936 | 0.070021972 |
| ENSMUSG00000083061 | Gm12191 | -0.477476395 | 0.016266259 |
| ENSMUSG00000024352 | Spata24 | -0.478440607 | 0.020444757 |
| ENSMUSG00000020376 | Rnf130 | -0.479908263 | 0.066200932 |
| ENSMUSG00000031591 | Asah1 | -0.480677783 | 0.012958158 |
| ENSMUSG00000026669 | Mcm10 | -0.480689498 | 0.096923883 |
| ENSMUSG00000026021 | Sumo1 | -0.481909019 | 0.087736508 |
| ENSMUSG00000053192 | Mllt11 | -0.485323244 | 0.004681039 |
| ENSMUSG00000026193 | Fn1 | -0.491554394 | 0.032185252 |
| ENSMUSG00000027698 | Nceh1 | -0.500948123 | 0.009384964 |
| ENSMUSG00000020372 | Rack1 | -0.504567205 | 0.054519251 |
| ENSMUSG00000019948 | Actr6 | -0.508481449 | 0.043690857 |
| ENSMUSG00000030545 | Pex11a | -0.510551038 | 0.058104748 |
| ENSMUSG00000108414 | Snhg1 | -0.512180993 | 0.061468333 |
| ENSMUSG00000034892 | Rps29 | -0.512709531 | 0.026829431 |
| ENSMUSG00000024694 | Keg1 | -0.513175688 | 0.059035783 |
| ENSMUSG00000067274 | Rplp0 | -0.514832242 | 0.012911524 |
| ENSMUSG00000041046 | Ramp3 | -0.515154668 | 0.005715492 |
| ENSMUSG00000030157 | Clec2d | -0.515578441 | 0.001893084 |
| ENSMUSG00000036814 | Slc6a20a | -0.516359138 | 0.038122128 |
| ENSMUSG00000021018 | Polr2h | -0.521961399 | 0.008488273 |
| ENSMUSG00000030268 | Bcat1 | -0.533457784 | 0.073511247 |
| ENSMUSG00000028111 | Ctsk | -0.535762202 | 0.043986365 |
| ENSMUSG00000021211 | Akr1c12 | -0.536375917 | 0.069471615 |
| ENSMUSG00000015013 | Trappc2l | -0.536513647 | 0.047641488 |
| ENSMUSG00000028081 | Rps3a1 | -0.537596212 | 0.024151443 |
| ENSMUSG00000027635 | Dsn1 | -0.540667338 | 0.014326612 |
| ENSMUSG00000020572 | Nampt | -0.543580525 | 0.014326612 |
| ENSMUSG00000037805 | Rpl10a | -0.545227766 | 0.062892557 |
| ENSMUSG00000003477 | Inmt | -0.548381386 | 0.060648268 |
| ENSMUSG00000010538 | Tsacc | -0.550224294 | 0.052639602 |
| ENSMUSG00000039646 | Vasn | -0.550345697 | 0.096358952 |
| ENSMUSG00000021417 | Eci2 | -0.553811598 | 0.002630447 |
| ENSMUSG00000090553 | Snrpe | -0.556824126 | 0.054203996 |
| ENSMUSG00000020482 | Ccdc117 | -0.55944931 | 0.003404664 |
| ENSMUSG00000002228 | Ppm1j | -0.560953441 | 0.092937686 |
| ENSMUSG00000030031 | Kbtbd8 | -0.564704459 | 0.076980136 |
| ENSMUSG00000056234 | Ncoa4 | -0.566706863 | 0.01221796 |
| ENSMUSG00000107689 | Gm44386 | -0.567049772 | 0.023169737 |
| ENSMUSG00000050097 | Ces2b | -0.569004259 | 0.091269869 |
| ENSMUSG00000079737 | 3110001I22Rik | -0.56959206 | 0.043840555 |
| ENSMUSG00000040812 | Agbl2 | -0.570791727 | 0.092298536 |
| ENSMUSG00000028127 | Abcd3 | -0.574803447 | 0.08241924 |
| ENSMUSG00000063696 | Gm8730 | -0.575030795 | 0.012668928 |
| ENSMUSG00000084983 | Gm11789 | -0.577478141 | 0.094051296 |
| ENSMUSG00000030747 | Dgat2 | -0.578721943 | 0.065990155 |
| ENSMUSG00000060036 | Rpl3 | -0.579444987 | 0.006388295 |
| ENSMUSG00000070427 | Il18bp | -0.583293588 | 0.041066817 |
| ENSMUSG00000118383 | AC132253.9 | -0.583414923 | 0.087164454 |
| ENSMUSG00000033318 | Gstt2 | -0.586917757 | 0.047631697 |
| ENSMUSG00000039395 | Mreg | -0.589151514 | 0.034958752 |
| ENSMUSG00000114774 | Gm35281 | -0.590282753 | 0.064323698 |
| ENSMUSG00000031105 | Slc25a14 | -0.598035458 | 0.023442258 |
| ENSMUSG00000028223 | Decr1 | -0.598578539 | 0.042814969 |
| ENSMUSG00000030968 | Pdilt | -0.604546892 | 0.015126351 |
| ENSMUSG00000005483 | Dnajb1 | -0.607448513 | 0.034958752 |
| ENSMUSG00000002076 | Hsf2bp | -0.607557725 | 0.064317008 |
| ENSMUSG00000069456 | Rdh16 | -0.609469938 | 0.05796034 |
| ENSMUSG00000087178 | A230056P14Rik | -0.613642128 | 0.010586335 |
| ENSMUSG00000074578 | Zfas1 | -0.617368664 | 0.042781899 |
| ENSMUSG00000018900 | Slc22a5 | -0.617655799 | 0.084155292 |
| ENSMUSG00000022579 | Gpihbp1 | -0.619712739 | 0.073519262 |
| ENSMUSG00000031253 | Srpx2 | -0.619877017 | 0.063218799 |
| ENSMUSG00000085203 | Gm12927 | -0.623262863 | 0.056463255 |
| ENSMUSG00000047728 | BC025446 | -0.623770411 | 0.042252146 |
| ENSMUSG00000037563 | Rps16 | -0.627497999 | 0.023005631 |
| ENSMUSG00000033634 | Nat8f2 | -0.629887776 | 0.099231093 |
| ENSMUSG00000028885 | Smpdl3b | -0.630314082 | 0.046947901 |
| ENSMUSG00000110216 | Gm36325 | -0.631951084 | 0.087436123 |
| ENSMUSG00000029322 | Plac8 | -0.636405064 | 0.01208111 |
| ENSMUSG00000039252 | Lgi2 | -0.640334706 | 0.072046655 |
| ENSMUSG00000028563 | Tm2d1 | -0.648316978 | 0.014713832 |
| ENSMUSG00000002346 | Slc25a42 | -0.649921148 | 0.084190091 |
| ENSMUSG00000060938 | Rpl26 | -0.656834362 | 0.018361175 |
| ENSMUSG00000022754 | Tmem45a | -0.660070019 | 0.042814969 |
| ENSMUSG00000028158 | Mttp | -0.667248172 | 0.074489848 |
| ENSMUSG00000041476 | Smpx | -0.673159535 | 0.03245408 |
| ENSMUSG00000040740 | Slc25a34 | -0.676478385 | 0.052613318 |
| ENSMUSG00000041301 | Cftr | -0.67935553 | 0.059035783 |
| ENSMUSG00000008683 | Rps15a | -0.684979367 | 0.033745285 |
| ENSMUSG00000030170 | Wnt5b | -0.688452188 | 0.014713832 |
| ENSMUSG00000066842 | Hmcn1 | -0.693831054 | 0.003840345 |
| ENSMUSG00000032602 | Slc25a20 | -0.694023597 | 0.010160935 |
| ENSMUSG00000117879 | AC138228.2 | -0.699634817 | 0.061680137 |
| ENSMUSG00000093402 | Gm18588 | -0.700590691 | 0.086281099 |
| ENSMUSG00000061947 | Serpina10 | -0.703584913 | 0.063230682 |
| ENSMUSG00000093674 | Rpl41 | -0.704621827 | 0.096112274 |
| ENSMUSG00000027171 | Prrg4 | -0.707822079 | 0.031163478 |
| ENSMUSG00000108763 | Gm36028 | -0.716419619 | 0.021710253 |
| ENSMUSG00000060143 | Gm10076 | -0.724089126 | 0.099245056 |
| ENSMUSG00000030877 | Mfsd13b | -0.730850368 | 0.021197175 |
| ENSMUSG00000003824 | Syce2 | -0.730907183 | 0.006458583 |
| ENSMUSG00000025491 | Ifitm1 | -0.734888618 | 0.03079587 |
| ENSMUSG00000063172 | Hspb11 | -0.743608855 | 0.011900932 |
| ENSMUSG00000039717 | Ralyl | -0.754599356 | 0.060648268 |
| ENSMUSG00000031725 | Ces1f | -0.754685368 | 0.031182829 |
| ENSMUSG00000012187 | Mogat1 | -0.769218563 | 0.060648268 |
| ENSMUSG00000020953 | Coch | -0.769372524 | 0.097792106 |
| ENSMUSG00000089669 | Tnfsf13 | -0.769724853 | 0.013232692 |
| ENSMUSG00000062611 | Rps3a2 | -0.77041162 | 0.007150945 |
| ENSMUSG00000109933 | Gm7600 | -0.771909027 | 0.016266259 |
| ENSMUSG00000019894 | Slc6a15 | -0.78759747 | 0.011293339 |
| ENSMUSG00000094685 | Gm5900 | -0.788086889 | 0.010543293 |
| ENSMUSG00000086583 | Gm15500 | -0.790631614 | 0.066200932 |
| ENSMUSG00000036775 | Decr2 | -0.797525638 | 0.013904092 |
| ENSMUSG00000028655 | Mfsd2a | -0.826747113 | 0.019751377 |
| ENSMUSG00000024827 | Gldc | -0.828160616 | 0.040175186 |
| ENSMUSG00000032690 | Oas2 | -0.831058724 | 0.096358952 |
| ENSMUSG00000116972 | Gm6278 | -0.854306386 | 0.068789096 |
| ENSMUSG00000085551 | 1810063I02Rik | -0.86292088 | 0.058680095 |
| ENSMUSG00000028212 | Ccne2 | -0.872749233 | 0.048186085 |
| ENSMUSG00000081087 | Rps15a-ps7 | -0.888911642 | 0.062566217 |
| ENSMUSG00000105466 | Gm42998 | -0.889007642 | 0.084003987 |
| ENSMUSG00000030256 | Bhlhe41 | -0.894257841 | 0.074460323 |
| ENSMUSG00000048368 | Omd | -0.896245998 | 0.078038352 |
| ENSMUSG00000044749 | Abca6 | -0.905673128 | 0.009083124 |
| ENSMUSG00000108599 | Gm10616 | -0.930001968 | 0.099363633 |
| ENSMUSG00000034402 | Kcnh5 | -0.970880536 | 0.003493291 |
| ENSMUSG00000006641 | Slc5a6 | -0.9775093 | 0.00059172 |
| ENSMUSG00000097296 | Gm26532 | -0.978014085 | 0.033154243 |
| ENSMUSG00000034706 | Dnaic2 | -0.993221892 | 0.050072317 |
| ENSMUSG00000086421 | Gm14091 | -1.002400338 | 0.077748576 |
| ENSMUSG00000081434 | Gm14165 | -1.016540225 | 0.016182902 |
| ENSMUSG00000020928 | Higd1b | -1.016690294 | 0.003448253 |
| ENSMUSG00000030154 | Klrb1f | -1.020016123 | 0.025133473 |
| ENSMUSG00000037440 | Vnn1 | -1.124033043 | 0.008361157 |
| ENSMUSG00000002068 | Ccne1 | -1.125240808 | 0.089759436 |
| ENSMUSG00000032921 | Odf4 | -1.135234536 | 0.057352574 |
| ENSMUSG00000081957 | Ak3l2-ps | -1.138696648 | 0.03925807 |
| ENSMUSG00000021388 | Aspn | -1.19654554 | 0.010965859 |
| ENSMUSG00000021213 | Akr1c13 | -1.250537687 | 0.003393129 |
| ENSMUSG00000026167 | Wnt10a | -1.276911387 | 0.038835355 |
| ENSMUSG00000028715 | Cyp4a14 | -1.28528728 | 0.087736508 |
| ENSMUSG00000021775 | Nr1d2 | -1.288976349 | 0.001365248 |
| ENSMUSG00000052396 | Mogat2 | -1.295249526 | 0.060648268 |
| ENSMUSG00000109341 | Gm30873 | -1.417980309 | 0.031717018 |
| ENSMUSG00000087343 | 1700021N21Rik | -1.428636713 | 0.027688938 |
| ENSMUSG00000057425 | Ugt2b37 | -1.477795021 | 0.051913031 |
| ENSMUSG00000049551 | Fzd9 | -1.503398486 | 0.087523304 |
| ENSMUSG00000058773 | Hist1h1b | -1.608949065 | 0.064087674 |
| ENSMUSG00000006724 | Cyp27b1 | -1.620210141 | 0.042252146 |
| ENSMUSG00000028957 | Per3 | -1.644218099 | 0.082198709 |
| ENSMUSG00000079494 | Nat8f5 | -2.094241354 | 0.037485154 |

Supplementary Table 3 C57BL/6J (RR-1) functional enrichment analysis hallmarks

| NAME | SIZE | NES | NOM p-val | FDR q-val |
| --- | --- | --- | --- | --- |
| HALLMARK_G2M_CHECKPOINT | 196 | 2.508 | 0.000 | 0.000 |
| HALLMARK_E2F_TARGETS | 200 | 2.488 | 0.000 | 0.000 |
| HALLMARK_CHOLESTEROL_HOMEOSTASIS | 73 | 2.477 | 0.000 | 0.000 |
| HALLMARK_INTERFERON_ALPHA_RESPONSE | 94 | 2.452 | 0.000 | 0.000 |
| HALLMARK_MTORC1_SIGNALING | 199 | 2.333 | 0.000 | 0.000 |
| HALLMARK_INTERFERON_GAMMA_RESPONSE | 195 | 2.230 | 0.000 | 0.000 |
| HALLMARK_MITOTIC_SPINDLE | 199 | 2.012 | 0.000 | 0.000 |
| HALLMARK_TNFA_SIGNALING_VIA_NFKB | 194 | 1.922 | 0.000 | 0.000 |
| HALLMARK_IL6_JAK_STAT3_SIGNALING | 83 | 1.754 | 0.000 | 0.003 |
| HALLMARK_ANGIOGENESIS | 35 | 1.632 | 0.016 | 0.012 |
| HALLMARK_UV_RESPONSE_UP | 151 | 1.495 | 0.006 | 0.043 |
| HALLMARK_GLYCOLYSIS | 195 | 1.469 | 0.003 | 0.052 |
| HALLMARK_HYPOXIA | 190 | 1.465 | 0.008 | 0.050 |
| HALLMARK_UNFOLDED_PROTEIN_RESPONSE | 112 | 1.454 | 0.019 | 0.052 |
| HALLMARK_SPERMATOGENESIS | 105 | 1.424 | 0.028 | 0.063 |
| HALLMARK_P53_PATHWAY | 198 | 1.422 | 0.008 | 0.060 |
| HALLMARK_ESTROGEN_RESPONSE_LATE | 183 | 1.413 | 0.010 | 0.061 |
| HALLMARK_COAGULATION | 129 | 1.345 | 0.054 | 0.109 |
| HALLMARK_ANDROGEN_RESPONSE | 98 | 1.339 | 0.060 | 0.108 |
| HALLMARK_APOPTOSIS | 156 | 1.317 | 0.061 | 0.125 |
| HALLMARK_EPITHELIAL_MESENCHYMAL_TRANSITION | 193 | 1.296 | 0.041 | 0.142 |
| HALLMARK_ADIPOGENESIS | 200 | 1.271 | 0.068 | 0.168 |
| HALLMARK_FATTY_ACID_METABOLISM | 152 | 1.241 | 0.104 | 0.203 |
| HALLMARK_INFLAMMATORY_RESPONSE | 187 | 1.209 | 0.109 | 0.245 |
| HALLMARK_PEROXISOME | 99 | 1.201 | 0.158 | 0.249 |
| HALLMARK_IL2_STAT5_SIGNALING | 195 | 1.190 | 0.139 | 0.256 |
| HALLMARK_HEME_METABOLISM | 188 | 1.157 | 0.183 | 0.304 |
| HALLMARK_PI3K_AKT_MTOR_SIGNALING | 102 | 1.147 | 0.223 | 0.311 |
| HALLMARK_PANCREAS_BETA_CELLS | 31 | 1.133 | 0.287 | 0.326 |
| HALLMARK_ESTROGEN_RESPONSE_EARLY | 189 | 1.123 | 0.233 | 0.334 |
| HALLMARK_XENOBIOTIC_METABOLISM | 192 | 1.123 | 0.228 | 0.324 |
| HALLMARK_APICAL_JUNCTION | 193 | 1.070 | 0.316 | 0.428 |
| HALLMARK_ALLOGRAFT_REJECTION | 186 | 1.056 | 0.344 | 0.445 |
| HALLMARK_BILE_ACID_METABOLISM | 109 | 1.034 | 0.372 | 0.483 |
| HALLMARK_MYOGENESIS | 196 | 1.031 | 0.391 | 0.474 |
| HALLMARK_COMPLEMENT | 188 | 0.987 | 0.490 | 0.565 |
| HALLMARK_MYC_TARGETS_V1 | 200 | 0.952 | 0.573 | 0.634 |
| HALLMARK_MYC_TARGETS_V2 | 57 | 0.915 | 0.599 | 0.709 |
| HALLMARK_KRAS_SIGNALING_DN | 168 | 0.904 | 0.686 | 0.715 |
| HALLMARK_APICAL_SURFACE | 43 | 0.867 | 0.677 | 0.781 |
| HALLMARK_OXIDATIVE_PHOSPHORYLATION | 198 | 0.678 | 0.993 | 1.000 |
| HALLMARK_PROTEIN_SECRETION | 95 | 0.498 | 1.000 | 0.999 |

Supplementary Table 4 BALB/c (RR-3) functional enrichment analysis hallmarks

| NAME | SIZE | NES | NOM p-val | FDR q-val |
| --- | --- | --- | --- | --- |
| HALLMARK_MYC_TARGETS_V1 | 200 | 2.248 | 0.000 | 0.000 |
| HALLMARK_OXIDATIVE_PHOSPHORYLATION | 198 | 1.977 | 0.000 | 0.000 |
| HALLMARK_ADIPOGENESIS | 200 | 1.956 | 0.000 | 0.000 |
| HALLMARK_FATTY_ACID_METABOLISM | 154 | 1.950 | 0.000 | 0.000 |
| HALLMARK_MYC_TARGETS_V2 | 57 | 1.770 | 0.000 | 0.004 |
| HALLMARK_P53_PATHWAY | 200 | 1.738 | 0.000 | 0.005 |
| HALLMARK_DNA_REPAIR | 149 | 1.713 | 0.000 | 0.005 |
| HALLMARK_E2F_TARGETS | 200 | 1.685 | 0.000 | 0.007 |
| HALLMARK_IL6_JAK_STAT3_SIGNALING | 85 | 1.625 | 0.004 | 0.012 |
| HALLMARK_EPITHELIAL_MESENCHYMAL_TRANSITION | 195 | 1.498 | 0.004 | 0.044 |
| HALLMARK_HYPOXIA | 195 | 1.468 | 0.007 | 0.054 |
| HALLMARK_COAGULATION | 136 | 1.407 | 0.027 | 0.088 |
| HALLMARK_PROTEIN_SECRETION | 95 | 1.406 | 0.029 | 0.082 |
| HALLMARK_ANGIOGENESIS | 36 | 1.384 | 0.065 | 0.090 |
| HALLMARK_BILE_ACID_METABOLISM | 112 | 1.368 | 0.043 | 0.097 |
| HALLMARK_UNFOLDED_PROTEIN_RESPONSE | 112 | 1.364 | 0.039 | 0.093 |
| HALLMARK_XENOBIOTIC_METABOLISM | 195 | 1.315 | 0.034 | 0.132 |
| HALLMARK_TNFA_SIGNALING_VIA_NFKB | 198 | 1.283 | 0.051 | 0.165 |
| HALLMARK_PEROXISOME | 101 | 1.260 | 0.086 | 0.187 |
| HALLMARK_ALLOGRAFT_REJECTION | 193 | 1.228 | 0.082 | 0.227 |
| HALLMARK_INFLAMMATORY_RESPONSE | 195 | 1.223 | 0.078 | 0.224 |
| HALLMARK_UV_RESPONSE_UP | 156 | 1.195 | 0.111 | 0.255 |
| HALLMARK_G2M_CHECKPOINT | 196 | 1.175 | 0.146 | 0.279 |
| HALLMARK_UV_RESPONSE_DN | 143 | 1.171 | 0.151 | 0.277 |
| HALLMARK_INTERFERON_GAMMA_RESPONSE | 195 | 1.152 | 0.165 | 0.301 |
| HALLMARK_KRAS_SIGNALING_UP | 197 | 1.127 | 0.182 | 0.342 |
| HALLMARK_GLYCOLYSIS | 197 | 1.124 | 0.204 | 0.336 |
| HALLMARK_ESTROGEN_RESPONSE_LATE | 194 | 1.087 | 0.245 | 0.409 |
| HALLMARK_MTORC1_SIGNALING | 199 | 1.079 | 0.276 | 0.415 |
| HALLMARK_HEME_METABOLISM | 189 | 1.078 | 0.290 | 0.402 |
| HALLMARK_APOPTOSIS | 160 | 1.063 | 0.327 | 0.427 |
| HALLMARK_COMPLEMENT | 191 | 1.060 | 0.307 | 0.420 |
| HALLMARK_SPERMATOGENESIS | 122 | 1.048 | 0.348 | 0.433 |
| HALLMARK_ANDROGEN_RESPONSE | 98 | 0.977 | 0.515 | 0.606 |
| HALLMARK_REACTIVE_OXYGEN_SPECIES_PATHWAY | 49 | 0.973 | 0.503 | 0.600 |
| HALLMARK_APICAL_JUNCTION | 197 | 0.899 | 0.693 | 0.790 |
| HALLMARK_INTERFERON_ALPHA_RESPONSE | 95 | 0.876 | 0.727 | 0.828 |
| HALLMARK_IL2_STAT5_SIGNALING | 198 | 0.793 | 0.929 | 0.970 |
| HALLMARK_APICAL_SURFACE | 44 | 0.758 | 0.866 | 0.988 |
| HALLMARK_PI3K_AKT_MTOR_SIGNALING | 105 | 0.726 | 0.955 | 0.988 |
| HALLMARK_HEDGEHOG_SIGNALING | 36 | 0.625 | 0.974 | 0.994 |

Supplementary Table 5 Comparison of enriched biological processes from functional enrichment analysis of DE genes in C57BL/6J (RR-1) and BALB/c (RR-3) (FDR < 0.25)

|  | **RR1 up** | | **RR1 down** |
| --- | --- | --- | --- |
| **RR3 up** | GOBP_CENTROMERE_COMPLEX_ASSEMBLY | | GOBP_COTRANSLATIONAL_PROTEIN_  TARGETING_TO_MEMBRANE |
|  | GOBP_CHROMATIN_REMODELING_AT_CENTROMERE | | GOBP_ESTABLISHMENT_OF_PROTEIN_  LOCALIZATION_TO_ENDOPLASMIC_RETICULUM |
|  | GOBP_TRIGLYCERIDE_METABOLIC_PROCESS | | GOBP_NUCLEAR_TRANSCRIBED_MRNA_  CATABOLIC_PROCESS_NONSENSE_MEDIATED_  DECAY |
|  | GOBP_BROWN_FAT_CELL_DIFFERENTIATION | | GOBP_NUCLEAR_TRANSCRIBED_MRNA_  CATABOLIC_PROCESS |
|  | GOBP_HISTONE_EXCHANGE | | GOBP_PROTEIN_LOCALIZATION_TO_  ENDOPLASMIC_RETICULUM |
|  | GOBP_TRIGLYCERIDE_BIOSYNTHETIC_PROCESS | | GOBP_TRANSLATIONAL_INITIATION |
|  | GOBP_DNA_REPLICATION_INDEPENDENT_NUCLEOSOME_  ORGANIZATION | | GOBP_CIRCADIAN_REGULATION_OF_GENE_  EXPRESSION |
|  | GOBP_MITOCHONDRIAL_TRANSLATIONAL_TERMINATION | | GOBP_ENTRAINMENT_OF_CIRCADIAN_CLOCK |
|  | GOBP_ANTIGEN_PROCESSING_AND_PRESENTATION_OF_  EXOGENOUS_PEPTIDE_ANTIGEN_VIA_MHC_CLASS_I | | GOBP_POSITIVE_REGULATION_OF_SIGNAL_  TRANSDUCTION_BY_P53_CLASS_MEDIATOR |
|  | GOBP_NEUTRAL_LIPID_BIOSYNTHETIC_PROCESS | | GOBP_VIRAL_GENE_EXPRESSION |
|  | GOBP_ANAPHASE_PROMOTING_COMPLEX_DEPENDENT_  CATABOLIC_PROCESS | | GOBP_RRNA_CATABOLIC_PROCESS |
|  | GOBP_POSITIVE_REGULATION_OF_COLD_INDUCED_  THERMOGENESIS | |  |
|  | GOBP_MITOCHONDRIAL_TRANSLATION | |  |
|  | GOBP_CELL_CYCLE_DNA_REPLICATION | |  |
|  | GOBP_DNA_DEPENDENT_DNA_REPLICATION | |  |
|  | GOBP_TRANSLATIONAL_TERMINATION | |  |
|  | GOBP_FOAM_CELL_DIFFERENTIATION | |  |
|  | GOBP_DNA_REPLICATION_INITIATION | |  |
|  | GOBP_REGULATION_OF_NUCLEASE_ACTIVITY | |  |
|  | GOBP_PLASMINOGEN_ACTIVATION | |  |
|  | GOBP_REGULATION_OF_MACROPHAGE_DERIVED_  FOAM_CELL_DIFFERENTIATION | |  |
|  | GOBP_MITOCHONDRIAL_GENE_EXPRESSION | |  |
|  | GOBP_COENZYME_A_METABOLIC_PROCESS | |  |
|  | GOBP_CELLULAR_LIPID_CATABOLIC_PROCESS | |  |
| **RR3 down** | GOBP_CALCIUM_ION_REGULATED_EXOCYTOSIS_  OF_NEUROTRANSMITTER | | GOBP_RENAL_SYSTEM_VASCULATURE_  DEVELOPMENT |
|  | GOBP_RESPONSE_TO_VITAMIN_A | | GOBP_RENAL_TUBULE_DEVELOPMENT |
|  | GOBP_NEGATIVE_REGULATION_OF_MICROTUBULE_  POLYMERIZATION_OR_DEPOLYMERIZATION | | GOBP_CARDIOCYTE_DIFFERENTIATION |

Supplementary Table 6 Genes with non-synonymous mutations between C57BL/6J and BALB/c (Timmermans, Van Montagu and Libert, 2017)

| 1110032A03Rik | Cilp2 | Gm4181 | Mrpl55 | Rab13 | Tpmt |
| --- | --- | --- | --- | --- | --- |
| 1190003K10Rik | Ckap2l | Gm43302 | Mrps10 | Rab29 | Tpo |
| 1520401A03Rik | Cldn10 | Gm43796 | Mrps26 | Rab3gap2 | Tpr |
| 1600002H07Rik | Cldn34c4 | Gm44511 | Mrps27 | Rab44 | Tpsb2 |
| 1600014C23Rik | Clec14a | Gm4924 | Mrps30 | Rab7b | Tpsg1 |
| 1700001J03Rik | Clec2g | Gm4955 | Mrps35 | Rabep2 | Tpte |
| 1700003E16Rik | Clec3b | Gm4969 | Ms4a14 | Rabgap1l | Traf3ip1 |
| 1700003H04Rik | Clec4f | Gm5105 | Ms4a2 | Rabl2 | Traf3ip3 |
| 1700006A11Rik | Clec7a | Gm5114 | Ms4a3 | Rad23b | Trafd1 |
| 1700008P02Rik | Clhc1 | Gm5150 | Ms4a4d | Rad51ap1 | Trak2 |
| 1700011H14Rik | Clip1 | Gm5475 | Ms4a5 | Rad51ap2 | Trappc12 |
| 1700011M02Rik | Clip4 | Gm5565 | Ms4a6b | Rad54l2 | Trappc13 |
| 1700013G24Rik | Clk2 | Gm5622 | Ms4a6d | Raet1d | Treml1 |
| 1700020A23Rik | Clk4 | Gm5678 | Msh2 | Raet1e | Trh |
| 1700024P16Rik | Cln3 | Gm590 | Msh3 | Ralbp1 | Trim10 |
| 1700029P11Rik | Clnk | Gm597 | Mslnl | Ralgapa2 | Trim17 |
| 1700030K09Rik | Clrn1 | Gm6086 | Msr1 | Ralgapb | Trim21 |
| 1700031F05Rik | Clstn2 | Gm609 | Mst1r | Ralgps2 | Trim23 |
| 1700057G04Rik | Clybl | Gm6309 | Mterf4 | Rangap1 | Trim25 |
| 1700066M21Rik | Cmpk1 | Gm6408 | Mtf2 | Rap1gap | Trim26 |
| 1700080E11Rik | Cmpk2 | Gm6563 | Mtfr2 | Rapgef5 | Trim27 |
| 1700123K08Rik | Cmtm5 | Gm6619 | Mtmr6 | Raph1 | Trim28 |
| 1810065E05Rik | Cmya5 | Gm6685 | Mtor | Rarres1 | Trim30b |
| 2010106E10Rik | Cnbd2 | Gm6793 | Mtrr | Rasa2 | Trim30c |
| 2010109I03Rik | Cnksr1 | Gm6812 | Mttp | Rasgrp4 | Trim30d |
| 2010111I01Rik | Cnnm1 | Gm7102 | Mtus1 | Rasip1 | Trim31 |
| 2200002D01Rik | Cnnm2 | Gm7534 | Muc15 | Rb1cc1 | Trim43b |
| 2210017I01Rik | Cntln | Gm7697 | Muc20 | Rbl2 | Trim45 |
| 2210407C18Rik | Cntn4 | Gm8186 | Muc5ac | Rbm20 | Trim56 |
| 2210408I21Rik | Cntn6 | Gm8882 | Muc6 | Rbm28 | Trim58 |
| 2310061I04Rik | Cntnap2 | Gm9495 | Mul1 | Rbm4 | Trim6 |
| 2410004B18Rik | Cntnap3 | Gm9573 | Mup8 | Rbm41 | Triml1 |
| 2410004P03Rik | Cntnap4 | Gm9774 | Mus81 | Rbms3 | Triml2 |
| 2700049A03Rik | Cntnap5a | Gm9881 | Musk | Rbpjl | Trip11 |
| 2700097O09Rik | Coa4 | Gmip | Mvk | Rbpms | Trmt1 |
| 2810004N23Rik | Cobl | Gnai2 | Mxra8 | Rccd1 | Trmt10a |
| 2810021J22Rik | Cog1 | Gnas | Myb | Rcn3 | Trmt10b |
| 2810428I15Rik | Cog2 | Gnpat | Myh1 | Recql4 | Trmt13 |
| 2900026A02Rik | Cog6 | Golga5 | Myh11 | Rem2 | Trmt2b |
| 3100002H09Rik | Cog8 | Gon4l | Mylk2 | Rfx7 | Trmt5 |
| 3110018I06Rik | Coil | Gp6 | Mylk3 | Rgl2 | Trpc4 |
| 3110040N11Rik | Col11a1 | Gpatch3 | Myo15b | Rgs11 | Trpm5 |
| 3632451O06Rik | Col11a2 | Gpatch8 | Myo18b | Rgs14 | Trps1 |
| 4921511C20Rik | Col16a1 | Gpc4 | Myo1b | Rgs20 | Trpv1 |
| 4921511H03Rik | Col19a1 | Gpd1 | Myo1e | Rhbdd3 | Trpv3 |
| 4921524L21Rik | Col20a1 | Gpi1 | Myo1f | Rhbdf1 | Trrap |
| 4921530L21Rik | Col22a1 | Gpn2 | Myo1g | Rhno1 | Tsc2 |
| 4921539E11Rik | Col24a1 | Gpnmb | Myo1h | Rhobtb3 | Tshb |
| 4930402H24Rik | Col27a1 | Gpr146 | Myo3b | Rhoq | Tspan11 |
| 4930415F15Rik | Col4a6 | Gpr149 | Myo5b | Rimbp2 | Tspan32 |
| 4930427A07Rik | Col6a6 | Gpr156 | Myo5c | Rimkla | Tspo2 |
| 4930430A15Rik | Col9a1 | Gpr19 | Myo7a | Rin1 | Tstd2 |
| 4930432K21Rik | Col9a2 | Gpr33 | Myo9b | Rin2 | Ttbk2 |
| 4930435E12Rik | Colec11 | Gpr63 | Myoc | Rin3 | Ttc12 |
| 4930438A08Rik | Commd7 | Gprc5a | Myof | Ring1 | Ttc21a |
| 4930503L19Rik | Cox18 | Gprc5d | Myom1 | Rint1 | Ttc22 |
| 4930519G04Rik | Cox6b2 | Gpx5 | Myrip | Riok1 | Ttc25 |
| 4930524B15Rik | Cpa6 | Gpx6 | Myt1l | Ripk1 | Ttc34 |
| 4930549C01Rik | Cpne1 | Gramd1a | Mzf1 | Rmdn1 | Ttc37 |
| 4930558K02Rik | Cpne3 | Gramd1c | N4bp2 | Rmnd1 | Ttc4 |
| 4930579F01Rik | Cpsf4l | Gramd3 | Naa16 | Rnase1 | Ttc5 |
| 4930595M18Rik | Cptp | Greb1 | Naa25 | Rnase10 | Ttc7 |
| 4931406P16Rik | Cpxcr1 | Grhl1 | Naaa | Rnase11 | Tti2 |
| 4931408C20Rik | Cpxm1 | Grik1 | Nab1 | Rnase2a | Tubgcp5 |
| 4932411E22Rik | Cpz | Grin2c | Nacad | Rnase6 | Tuft1 |
| 4933411K16Rik | Cramp1l | Grin2d | Naip2 | Rnaseh1 | Tusc1 |
| 4933427I04Rik | Crcp | Grip1 | Naip5 | Rnaseh2c | Txndc12 |
| 5430403G16Rik | Crhr2 | Grk6 | Naip6 | Rnf150 | Tyr |
| 5730507C01Rik | Cript | Grm2 | Nbn | Rnf157 | Tyro3 |
| 5730508B09Rik | Crlf2 | Grrp1 | Ncan | Rnf212 | Tyrp1 |
| 5730522E02Rik | Crocc2 | Gsdmc2 | Ncapd2 | Rnf34 | Tyw5 |
| 5830473C10Rik | Crp | Gsg1 | Ncapg2 | Rnf6 | Uba6 |
| 5930422O12Rik | Crtap | Gspt2 | Nccrp1 | Rnmtl1 | Ubash3a |
| 6030458C11Rik | Crtc3 | Gsr | Ncf4 | Rock2 | Ube2l6 |
| 6430550D23Rik | Crybg3 | Gstm6 | Nck1 | Rorc | Ube2ql1 |
| 6430573F11Rik | Csf2rb2 | Gtf2e2 | Nckap5l | Rp1 | Ube2u |
| 7420426K07Rik | Csf3r | Gtf2h3 | Ncoa3 | RP23-100C12.1 | Ube2z |
| 7420461P10Rik | Cspg5 | Gtf2h4 | Ncoa6 | RP23-281F13.1 | Ube4a |
| 9030624G23Rik | Csrnp2 | Gtf3c1 | Ncr1 | RP23-348B21.1 | Ubiad1 |
| 9030624J02Rik | Cstl1 | Gtpbp6 | Ncstn | RP23-414I24.11 | Ubox5 |
| 9130019O22Rik | Ctbp2 | Guca1b | Ndc80 | RP24-351P7.8 | Ubqlnl |
| 9630041A04Rik | Ctbs | Gucy1a3 | Ndst3 | Rpap1 | Ubr1 |
| 9930022D16Rik | Ctgf | Gucy1b2 | Ndufaf1 | Rpap2 | Ubr2 |
| a | Ctif | Gucy2c | Ndufaf7 | Rpe65 | Ubr4 |
| A130010J15Rik | Cts3 | Gucy2d | Nectin1 | Rpgrip1 | Ubtf |
| A430033K04Rik | Ctse | Gusb | Nek11 | Rpgrip1l | Ubxn2b |
| A830010M20Rik | Ctsm | Gykl1 | Nek2 | Rps18 | Ubxn4 |
| A930018P22Rik | Ctss | H13 | Nek3 | Rps6ka6 | Ucp1 |
| Aadac | Cttnbp2 | H2-Aa | Nelfe | Rptn | Ufl1 |
| Aars | Cuedc1 | H2-Ab1 | Nes | Rpusd1 | Uggt2 |
| Abca13 | Cul9 | H2-D1 | Neto1 | Rpusd2 | Ugt2b36 |
| Abca14 | Cux1 | H2-DMb1 | Neurl1a | Rrp12 | Ugt8a |
| Abca17 | Cwc25 | H2-Eb1 | Neurl1b | Rsad2 | Uhrf1bp1l |
| Abca3 | Cxcl5 | H2-Eb2 | Nfatc2 | Rsl1 | Ulk4 |
| Abca5 | Cxcl9 | H2-K1 | Nfe2l3 | Rsph3b | Umod |
| Abca6 | Cxcr4 | H2-Ke6 | Nfkbib | Rtbdn | Umodl1 |
| Abca9 | Cyb5d1 | H2-M10.5 | Nfkbid | Rtf1 | Unc13b |
| Abcb10 | Cyb5r4 | H2-M10.6 | Nfu1 | Rtkn | Unc45a |
| Abcc2 | Cyp2a12 | H2-M2 | Ngdn | Rtkn2 | Unc5cl |
| Abcc3 | Cyp2a4 | H2-Ob | Ngfr | Rtn1 | Uncx |
| Abcc4 | Cyp2b23 | H2-Q6 | Nhs | Ryr1 | Unkl |
| Abcc6 | Cyp2c29 | H2-T22 | Nhsl1 | Sacs | Upf2 |
| Abhd10 | Cyp2c37 | H60b | Nid2 | Sall2 | Uqcc1 |
| Abhd4 | Cyp2c38 | H6pd | Nin | Sall4 | Urb2 |
| Abhd8 | Cyp2c50 | Habp4 | Nipal2 | Samd11 | Uri1 |
| Abi3bp | Cyp2c68 | Hacd4 | Nipsnap3b | Samd14 | Usb1 |
| Ablim1 | Cyp2c69 | Hagh | Nit1 | Samd5 | Use1 |
| Acacb | Cyp2d34 | Hapln3 | Nkapl | Samd8 | Ush1c |
| Acad10 | Cyp2f2 | Harbi1 | Nkd1 | Samd9l | Ush2a |
| Acad11 | Cyp2j12 | Hars | Nktr | Sapcd1 | Ushbp1 |
| Acads | Cyp2j13 | Has1 | Nkx2-2 | Saraf | Usp18 |
| Acan | Cyp39a1 | Has3 | Nkx2-3 | Sash1 | Usp33 |
| Acap3 | Cyp3a25 | Haus4 | Nlgn2 | Sbk2 | Usp38 |
| Acat3 | Cyp3a57 | Haus5 | Nlrc5 | Sbk3 | Usp44 |
| Acin1 | Cyp3a59 | Haus6 | Nlrp12 | Sbp | Usp48 |
| Ackr1 | Cyp4a10 | Haus8 | Nlrp14 | Sbpl | Utp14b |
| Acox1 | Cyp4x1 | Havcr1 | Nlrp1b | Sc5d | Utp20 |
| Acpp | Cyp7a1 | Havcr2 | Nlrp4e | Scaf1 | Uvrag |
| Acsm3 | Cyp7b1 | Hba-x | Nlrp4f | Scand1 | Vars2 |
| Acss2 | Cyp8b1 | Hbb-bh2 | Nlrp5 | Scd4 | Vcan |
| Actl11 | Cypt3 | Hbb-bs | Nlrp9a | Scfd2 | Vcl |
| Acvrl1 | Cytl1 | Hbb-bt | Nlrp9c | Scgb1b12 | Veph1 |
| Acy1 | D130040H23Rik | Hbq1a | Nmb | Scgb2b11 | Vezt |
| Adal | D130052B06Rik | Hcar1 | Nme8 | Scgb2b12 | Vgll1 |
| Adam11 | D430041D05Rik | Hcls1 | Noc3l | Scgb2b18 | Vill |
| Adam12 | D430042O09Rik | Hcn1 | Nod2 | Scgb2b2 | Vit |
| Adam17 | D5Ertd579e | Hcrtr2 | Nol10 | Scn5a | Vmn1r172 |
| Adam18 | D630036H23Rik | Hdac4 | Nol12 | Scpep1 | Vmn1r173 |
| Adam24 | D8Ertd82e | Heatr3 | Nop14 | Sctr | Vmn1r181 |
| Adam25 | D930015E06Rik | Heatr5a | Nop2 | Sdad1 | Vmn1r232 |
| Adam29 | D930020B18Rik | Hebp1 | Nop56 | Sdccag8 | Vmn1r236 |
| Adam33 | Dact1 | Heca | Nop58 | Sdhaf2 | Vmn1r66 |
| Adam39 | Dag1 | Heph | Notch4 | Sdhaf4 | Vmn1r67 |
| Adam6a | Dao | Hesx1 | Noxo1 | Sdk2 | Vmn1r72 |
| Adamts18 | Dars | Hgsnat | Npc1l1 | Sec1 | Vmn2r121 |
| Adamts19 | Dars2 | Hhat | Npcd | Sec16b | Vmn2r30 |
| Adamts3 | Dcaf15 | Hif1an | Npffr2 | Sec22c | Vmn2r88 |
| Adamts7 | Dchs2 | Hif3a | Nphp3 | Sec24d | Vmn2r89 |
| Adamtsl3 | Dclk3 | Hipk3 | Nphs1 | Sec31b | Vnn3 |
| Adamtsl4 | Dcp2 | Hirip3 | Nphs2 | Sectm1b | Vps13b |
| Adcy7 | Dcpp1 | Hivep2 | Npm2 | Sel1l3 | Vps13c |
| Adgra3 | Dcpp2 | Hivep3 | Npr1 | Selenbp1 | Vps33a |
| Adgrb1 | Dcpp3 | Hk3 | Nprl3 | Sema3a | Vps37b |
| Adgre5 | Dcst2 | Hmgcs2 | Nptxr | Sema3b | Vps52 |
| Adgrf1 | Dctn1 | Hnf1a | Nr0b2 | Sema3f | Vrk2 |
| Adgrf4 | Dctn3 | Hnrnpul2 | Nr1h4 | Sema4f | Vrk3 |
| Adgrg6 | Ddhd1 | Homer3 | Nr2c1 | Sema5b | Vsig10 |
| Adgrv1 | Ddi2 | Homez | Nrg1 | Sema6a | Vtcn1 |
| Adh4 | Ddost | Hook2 | Nsl1 | Senp6 | Vti1b |
| Adh6a | Ddx18 | Hormad1 | Nsun2 | Serinc4 | Vwa5b1 |
| Adh6b | Ddx20 | Hoxa4 | Ntn4 | Serpina12 | Vwa8 |
| Adh7 | Ddx55 | Hoxd4 | Ntsr2 | Serpina1a | Vwce |
| Adrbk2 | Decr1 | Hp1bp3 | Nudt7 | Serpina1c | Vwde |
| Afap1l1 | Defa24 | Hps1 | Nuggc | Serpina1d | Vwf |
| Aff2 | Defa30 | Hps6 | Numa1 | Serpina1f | Wbp11 |
| Aftph | Defa35 | Hr | Nup210l | Serpina3b | Wdr46 |
| Agbl1 | Defb40 | Hrct1 | Nup62 | Serpina3c | Wdr72 |
| Agrn | Defb8 | Hs3st3a1 | Nup85 | Serpina3g | Wdr76 |
| Agt | Dek | Hs3st4 | Nup98 | Serpina3j | Wfdc6b |
| AI429214 | Dennd2d | Hs3st6 | Nupl2 | Serpina3k | Wfikkn2 |
| AI481877 | Dennd4c | Hscb | Nwd2 | Serpina3m | Whsc1 |
| AI607873 | Depdc1b | Hsf3 | Nxpe2 | Serpina3n | Wisp2 |
| Aim1l | Depdc5 | Hsh2d | Nxpe4 | Serpina9 | Wwc1 |
| Aim2 | Dgkq | Hspa12a | Oacyl | Serpinb3b | Xkr5 |
| Aimp2 | Dgkz | Hspb2 | Oasl2 | Serpinb3c | Xkr9 |
| Ak7 | Dguok | Hspb6 | Obox3 | Serpinb3d | Xkrx |
| Akap6 | Dhdh | Hspg2 | Ocel1 | Serpinb9b | Xlr5b |
| Akap9 | Dhtkd1 | Htr1d | Odc1 | Serpine1 | Xpnpep2 |
| Akr1c12 | Dhx30 | Htr5b | Ofcc1 | Serpine3 | Xpo7 |
| Akr1c13 | Dhx34 | Htra2 | Oip5 | Sertad1 | Xpot |
| Akr1c21 | Dhx36 | Huwe1 | Olfml3 | Setd1b | Xrcc4 |
| Akr7a5 | Diablo | Hvcn1 | Olfr101 | Setmar | Xrcc6 |
| Aldh1l2 | Diaph3 | Hyal1 | Olfr1015 | Sez6l | Xrra1 |
| Aldh4a1 | Disc1 | Hyal2 | Olfr102 | Sfi1 | Xxylt1 |
| Alg14 | Disp1 | Hyal3 | Olfr1026 | Sfrp1 | Ybx3 |
| Allc | Disp2 | Hyi | Olfr1029 | Sfta2 | Yipf1 |
| Aloxe3 | Dixdc1 | Iah1 | Olfr1036 | Sfxn4 | Zan |
| Alpk1 | Dlat | Ibsp | Olfr1049 | Sgcz | Zbed5 |
| Alpk2 | Dlc1 | Iffo1 | Olfr1055 | Sgol2b | Zbtb17 |
| Alpk3 | Dlgap4 | Ifi202b | Olfr1056 | Sgsh | Zbtb2 |
| Alpl | Dlx4 | Ifi205 | Olfr107 | Sgsm1 | Zbtb3 |
| Als2 | Dmp1 | Ifi44 | Olfr108 | Sgtb | Zbtb32 |
| Als2cl | Dmtn | Ifit1 | Olfr110 | Sh2d5 | Zbtb38 |
| Als2cr11 | Dmwd | Ifna2 | Olfr1102 | Sh3bp1 | Zbtb39 |
| Amer2 | Dnah11 | Ifnab | Olfr1115 | Sh3gl2 | Zbtb7c |
| Amph | Dnah12 | Ifngr1 | Olfr114 | Sh3gl3 | Zc3h7b |
| Amtn | Dnah6 | Ifrd2 | Olfr1157 | Sh3pxd2b | Zcchc11 |
| Ang4 | Dnah7a | Ift140 | Olfr1161 | Sh3tc2 | Zcchc6 |
| Angptl1 | Dnah8 | Igbp1b | Olfr1180 | Shb | Zdhhc2 |
| Ank2 | Dnah9 | Igf1r | Olfr1215 | Shc4 | Zeb1 |
| Ankhd1 | Dnaja2 | Igflr1 | Olfr1225 | Shisa9 | Zfp105 |
| Ankle1 | Dnajb13 | Igsf11 | Olfr125 | Shkbp1 | Zfp112 |
| Ankmy2 | Dnajc11 | Igsf21 | Olfr128 | Shmt2 | Zfp113 |
| Ankrd11 | Dnajc17 | Igsf8 | Olfr1315-ps1 | Shprh | Zfp157 |
| Ankrd17 | Dnajc24 | Igsf9 | Olfr1362 | Shroom3 | Zfp160 |
| Ankrd27 | Dnajc6 | Igtp | Olfr137 | Siglec1 | Zfp180 |
| Ankrd36 | Dnhd1 | Ikbip | Olfr1406 | Sipa1 | Zfp185 |
| Ankrd55 | Dnm3 | Ikbkap | Olfr1434 | Sipa1l2 | Zfp213 |
| Ankrd61 | Dnmbp | Ikbke | Olfr1438-ps1 | Sirpa | Zfp235 |
| Anks6 | Dntt | Il12rb1 | Olfr1441 | Sirpb1a | Zfp251 |
| Anxa3 | Dock2 | Il17b | Olfr1509 | Sirpb1b | Zfp27 |
| Aoah | Dock3 | Il17ra | Olfr1510 | Sirpb1c | Zfp273 |
| Aox2 | Dock5 | Il17rd | Olfr154 | Sis | Zfp282 |
| Aox3 | Dok1 | Il20ra | Olfr155 | Ska1 | Zfp30 |
| Aox4 | Dpcr1 | Il20rb | Olfr160 | Skap1 | Zfp334 |
| Ap4e1 | Dpep1 | Il21r | Olfr172 | Skint1 | Zfp341 |
| Ap5m1 | Dpp10 | Il27ra | Olfr181 | Skint10 | Zfp365 |
| Apc | Dpp6 | Il31ra | Olfr187 | Skint2 | Zfp367 |
| Apeh | Dscaml1 | Il3ra | Olfr199 | Skint7 | Zfp369 |
| Apoa1 | Dspp | Il4i1 | Olfr205 | Skint8 | Zfp382 |
| Apoa2 | Dtd2 | Il4ra | Olfr209 | Slamf7 | Zfp384 |
| Apob | Dthd1 | Il9r | Olfr221 | Slamf8 | Zfp386 |
| Apobec3 | Duox1 | Immp1l | Olfr248 | Slamf9 | Zfp39 |
| Apoc3 | Duoxa1 | Ina | Olfr25 | Slc12a9 | Zfp398 |
| Apol10b | Dus4l | Inafm1 | Olfr313 | Slc13a1 | Zfp40 |
| Apol7c | Dusp16 | Inpp4b | Olfr314 | Slc15a1 | Zfp408 |
| Aqr | Dusp19 | Insl3 | Olfr316 | Slc16a12 | Zfp42 |
| Arap1 | Dusp4 | Insl5 | Olfr317 | Slc17a1 | Zfp429 |
| Arap2 | Dynap | Ints10 | Olfr320 | Slc17a3 | Zfp446 |
| Arap3 | E130208F15Rik | Ints8 | Olfr324 | Slc20a2 | Zfp456 |
| Arg2 | E2f2 | Ints9 | Olfr325 | Slc22a13 | Zfp457 |
| Arhgap1 | E2f6 | Ip6k3 | Olfr328 | Slc22a15 | Zfp458 |
| Arhgap19 | E4f1 | Ipcef1 | Olfr329-ps | Slc22a17 | Zfp462 |
| Arhgap23 | Eapp | Iqcf6 | Olfr370 | Slc24a4 | Zfp493 |
| Arhgap36 | Ear6 | Iqcg | Olfr374 | Slc26a11 | Zfp507 |
| Arhgap39 | Ece1 | Iqgap1 | Olfr414 | Slc26a9 | Zfp536 |
| Arhgap40 | Echdc2 | Iqgap2 | Olfr418 | Slc27a1 | Zfp551 |
| Arhgap8 | Eci2 | Irf2bp2 | Olfr420 | Slc27a3 | Zfp566 |
| Arhgap9 | Ecm1 | Irgm2 | Olfr429 | Slc27a6 | Zfp568 |
| Arhgef11 | Ect2l | Irs4 | Olfr464 | Slc28a1 | Zfp592 |
| Arhgef12 | Edem2 | Irx5 | Olfr49 | Slc28a2 | Zfp595 |
| Arhgef17 | Efcab6 | Isg20l2 | Olfr521 | Slc2a9 | Zfp598 |
| Arhgef2 | Efcc1 | Isoc1 | Olfr544 | Slc30a9 | Zfp605 |
| Arhgef28 | Efs | Ispd | Olfr557 | Slc34a1 | Zfp609 |
| Arhgef37 | Egf | Itga9 | Olfr560 | Slc35c2 | Zfp618 |
| Arhgef40 | Ehbp1 | Itgae | Olfr566 | Slc35d2 | Zfp619 |
| Arl14 | Ehbp1l1 | Itgal | Olfr574 | Slc35e1 | Zfp64 |
| Arntl2 | Ehf | Itgav | Olfr6 | Slc35f4 | Zfp644 |
| Arrdc2 | Eif2ak2 | Itgb2l | Olfr600 | Slc38a3 | Zfp655 |
| Arsk | Eif2ak4 | Itgb4 | Olfr601 | Slc39a2 | Zfp672 |
| Art4 | Eif2b3 | Itgbl1 | Olfr605 | Slc39a7 | Zfp677 |
| Art5 | Elk3 | Itih1 | Olfr609 | Slc44a1 | Zfp68 |
| As3mt | Elovl5 | Itih3 | Olfr62 | Slc46a2 | Zfp683 |
| Asap2 | Emc1 | Itih4 | Olfr623 | Slc4a11 | Zfp69 |
| Asb1 | Eme2 | Itpa | Olfr629 | Slc4a5 | Zfp692 |
| Asb14 | Emilin2 | Itpr3 | Olfr652 | Slc5a5 | Zfp7 |
| Asb16 | Endov | Jade1 | Olfr653 | Slc5a7 | Zfp708 |
| Asb3 | Enoph1 | Jak3 | Olfr69 | Slc5a8 | Zfp709 |
| Ascc2 | Enox1 | Jhy | Olfr70 | Slc5a9 | Zfp710 |
| Ash1l | Enpp1 | Jmy | Olfr711 | Slc6a17 | Zfp712 |
| Asl | Enpp3 | Kap | Olfr723 | Slc6a4 | Zfp729a |
| Aspa | Enpp5 | Kars | Olfr724 | Slc7a2 | Zfp738 |
| Asph | Enpp7 | Katna1 | Olfr727 | Slc7a5 | Zfp746 |
| Asphd2 | Entpd1 | Kbtbd11 | Olfr733 | Slc9a3r1 | Zfp748 |
| Asrgl1 | Eogt | Kbtbd4 | Olfr745 | Slc9a3r2 | Zfp759 |
| Aste1 | Epas1 | Kcnb2 | Olfr747 | Slc9c1 | Zfp760 |
| Astn1 | Epb41l1 | Kcnd3 | Olfr750 | Slco2b1 | Zfp772 |
| Asxl3 | Epb41l2 | Kcnh1 | Olfr761 | Slco5a1 | Zfp773 |
| Atad5 | Epb41l4b | Kcnip3 | Olfr875 | Slf1 | Zfp775 |
| Atat1 | Epb42 | Kcnj10 | Olfr893 | Slit1 | Zfp790 |
| Atf6b | Epc1 | Kcnj14 | Olfr902 | Slit2 | Zfp791 |
| Atf7ip | Epha3 | Kcns3 | Olfr905 | Slitrk3 | Zfp81 |
| Atg4b | Epha5 | Kctd14 | Olfr93 | Smad9 | Zfp82 |
| Atg4c | Epm2a | Kctd2 | Olfr996 | Smc2 | Zfp820 |
| Atp10a | Epn3 | Kctd5 | Olfr998 | Smco2 | Zfp84 |
| Atp13a3 | Eprs | Kdm2b | Ooep | Smco3 | Zfp865 |
| Atp1a4 | Eps15l1 | Kdm8 | Oosp3 | Smek2 | Zfp868 |
| Atp5g1 | Epx | Khdc1a | Orc1 | Smg6 | Zfp874a |
| Atp5j | Eqtn | Khdrbs2 | Orc6 | Smim10l1 | Zfp874b |
| Atp5s | Erc2 | KIAA1683 | Osbpl2 | Smim5 | Zfp882 |
| Atp6ap1l | Ercc1 | Kidins220 | Otogl | Smo | Zfp930 |
| Atp6v0b | Ercc6l2 | Kif11 | Otud3 | Smoc1 | Zfp932 |
| Atp6v1h | Ercc8 | Kif13a | Ovgp1 | Smok3c | Zfp935 |
| Atp7a | Eri1 | Kif17 | Oxa1l | Smpdl3b | Zfp938 |
| Atp8a2 | Eri2 | Kif1c | Oxgr1 | Snai3 | Zfp940 |
| Atrip | Erich2 | Kif20b | Oxsm | Sned1 | Zfp944 |
| Atrnl1 | Erich3 | Kif4 | P2rx4 | Snx1 | Zfp945 |
| Atrx | Erp27 | Kif7 | P3h1 | Snx20 | Zfp948 |
| Atxn7l1 | Erp29 | Kifc1 | Pabpn1l | Snx7 | Zfp949 |
| AU018091 | Esco1 | Kifc3 | Padi2 | Soat1 | Zfp950 |
| Aunip | Esyt2 | Kirrel2 | Padi3 | Socs5 | Zfp951 |
| Avp | Etaa1 | Kiz | Padi4 | Son | Zfp954 |
| AW551984 | Etv2 | Klf1 | Pafah2 | Sorcs1 | Zfp961 |
| Axdnd1 | Etv3 | Klf11 | Pak6 | Sorl1 | Zfp963 |
| AY761184 | Eva1a | Klf17 | Palm3 | Sox6 | Zfp974 |
| Aym1 | Evc | Klhl1 | Pamr1 | Spaca4 | Zfp994 |
| B2m | Evc2 | Klhl28 | Pan2 | Spaca6 | Zfyve19 |
| B3glct | Evpl | Klhl41 | Pan3 | Spag11b | Zfyve9 |
| B3gnt3 | Exd1 | Klk1b26 | Pank4 | Sparcl1 | Zgrf1 |
| B3gnt4 | Exo1 | Klk1b27 | Pappa2 | Spata13 | Zkscan16 |
| Babam1 | Exoc3l2 | Klra10 | Paqr8 | Spata18 | Zkscan3 |
| Bace2 | Exoc8 | Klra17 | Parp2 | Spata2l | Zkscan5 |
| Bag3 | Exog | Klra2 | Parp8 | Spata31 | Zkscan7 |
| Bahcc1 | Exosc10 | Klra3 | Parpbp | Spata31d1b | Zmym4 |
| Bahd1 | Ext2 | Klra4 | Pars2 | Spata31d1c | Zmym6 |
| Baiap3 | Extl1 | Klra7 | Pask | Spata31d1d | Zmynd10 |
| Bank1 | Extl3 | Klra8 | Patl2 | Spata33 | Zmynd8 |
| Batf3 | F2rl1 | Klrb1 | Pax1 | Spata45 | Znhit2 |
| BC016579 | F830016B08Rik | Klrb1a | Pbx2 | Spata6 | Znrf3 |
| BC026585 | F830045P16Rik | Klrb1c | Pbx4 | Spdya | Zp2 |
| BC034090 | Faap100 | Klrc1 | Pcbp4 | Speer4b | Zranb3 |
| BC035044 | Fads2 | Klrc3 | Pcdh11x | Spen | Zscan10 |
| BC037034 | Fahd1 | Klrd1 | Pcdh18 | Spg11 | Zscan2 |
| BC048403 | Faim2 | Klri1 | Pcdh20 | Spg7 | Zscan29 |
| BC048546 | Fam109b | Klrk1 | Pcdh9 | Spink13 | Zscan4b |
| BC049715 | Fam114a1 | Kmt2e | Pcdha3 | Spink2 | Zscan4d |
| BC051142 | Fam114a2 | Kndc1 | Pcdha5 | Sppl2a | Zswim2 |
| BC053393 | Fam126a | Kntc1 | Pcdha9 | Spred2 | Zswim6 |
| BC107364 | Fam151a | Kpna2 | Pcdhb10 | Spred3 | Zw10 |
| BC117090 | Fam160b2 | Kpna4 | Pcdhga10 | Sprr2i |  |
| Bcas1 | Fam171a2 | Krt13 | Pcdhga11 | Sprtn |  |
| Bcdin3d | Fam171b | Krt24 | Pcdhga12 | Spsb3 |  |
| Bcl2a1b | Fam174b | Krt27 | Pcdhga3 | Spta1 |  |
| Bcl2a1c | Fam178a | Krt39 | Pcdhga4 | Sptbn2 |  |
| Bdh1 | Fam179a | Krt5 | Pcdhga6 | Sptlc1 |  |
| Bdnf | Fam186a | Krt74 | Pcdhga7 | Sqrdl |  |
| Bdp1 | Fam188b | Krt83 | Pcdhga9 | Sra1 |  |
| Best2 | Fam189a2 | Krt90 | Pcdhgb1 | Srd5a1 |  |
| Bid | Fam204a | Krtap4-2 | Pcdhgb2 | Srp68 |  |
| Bloc1s6 | Fam208b | Krtap4-7 | Pcdhgb5 | Srpr |  |
| Blvra | Fam214a | Krtap4-9 | Pcdhgb6 | Srrm2 |  |
| Bmp5 | Fam214b | Krtap5-3 | Pcdhgb7 | Ssc4d |  |
| Boc | Fam221b | Krtap5-5 | Pcdhgc3 | Ssfa2 |  |
| Bod1 | Fam234b | Ksr2 | Pced1a | Ssh1 |  |
| Borcs5 | Fam63b | L1td1 | Pcf11 | Sspo |  |
| Bpifa5 | Fam71e1 | Lacc1 | Pclo | Ssx2ip |  |
| Bpifa6 | Fam71e2 | Lactbl1 | Pcm1 | Stambpl1 |  |
| Bpifb4 | Fam72a | Lama1 | Pcsk9 | Stard4 |  |
| Bpifb6 | Fam73a | Lama2 | Pdcd11 | Stard7 |  |
| Bpnt1 | Fam81b | Lama3 | Pdcl3 | Stard8 |  |
| Brca2 | Fam83c | Lama5 | Pde11a | Stard9 |  |
| Brd8 | Fam90a1a | Lamb1 | Pde12 | Stat2 |  |
| Brinp2 | Fam98c | Lao1 | Pde2a | Steap3 |  |
| Bst1 | Fanca | Laptm4b | Pde3b | Stfa1 |  |
| Btaf1 | Fancc | Larp1 | Pde4b | Stil |  |
| Btbd10 | Farp1 | Larp4 | Pde4d | Stk17b |  |
| Btbd19 | Farp2 | Larp7 | Pde6a | Stk25 |  |
| Btg4 | Farsa | Lbp | Pdf | Stk31 |  |
| Btla | Fas | Lbx2 | Pdgfb | Stk32b |  |
| Btnl1 | Fasl | Lce1a2 | Pdgfra | Stra6l |  |
| Btnl10 | Fastkd1 | Lce1c | Pdhx | Stt3b |  |
| Btnl2 | Fastkd3 | Lce1m | Pdia4 | Stx3 |  |
| Bub1b | Fastkd5 | Lce3d | Pdia6 | Stxbp5l |  |
| Bzrap1 | Fat4 | Lct | Pdilt | Stxbp6 |  |
| Bzw2 | Fblim1 | Lect2 | Pdlim5 | Styk1 |  |
| C130079G13Rik | Fbln7 | Lelp1 | Pdpr | Suclg1 |  |
| C1qa | Fbn1 | Leng1 | Pdzd7 | Suco |  |
| C1qc | Fbn2 | Leo1 | Pdzk1ip1 | Sugp1 |  |
| C1ra | Fbxl5 | Leprot | Pebp1 | Sugp2 |  |
| C2cd2 | Fbxo2 | Letmd1 | Peg12 | Sult2b1 |  |
| C2cd3 | Fbxo27 | Lgi4 | Peli2 | Sult5a1 |  |
| C2cd4a | Fbxo34 | Lilra5 | Perm1 | Sult6b1 |  |
| C4b | Fbxo38 | Lima1 | Pex19 | Sumf2 |  |
| C5ar2 | Fbxw14 | Lmntd1 | Pfpl | Supt20 |  |
| C7 | Fbxw16 | Lmod3 | Pgam2 | Supv3l1 |  |
| Cables1 | Fbxw18 | Lmtk3 | Pgam5 | Susd1 |  |
| Cacna1h | Fcer1a | Lnp1 | Pgap2 | Susd5 |  |
| Cacna1i | Fcer1g | Lnx1 | Pgbd5 | Svs3b |  |
| Cacna2d4 | Fcgr3 | Loxl4 | Pgc | Syde2 |  |
| Cad | Fcho1 | Lpar2 | Pggt1b | Syne4 |  |
| Cadps | Fcmr | Lpcat2 | Pgls | Syngr4 |  |
| Cadps2 | Fcrl1 | Lpcat2b | Pglyrp3 | Synj2 |  |
| Cage1 | Fcrl5 | Lpgat1 | Phc1 | Synm |  |
| Calm4 | Fcrl6 | Lpin1 | Phf20 | Syt7 |  |
| Calm5 | Fcrls | Lpl | Phf21b | Sytl1 |  |
| Camk1g | Fdxr | Lpo | Phldb1 | Syvn1 |  |
| Camkv | Fer1l6 | Lrba | Phyh | Szt2 |  |
| Cap1 | Fez1 | Lrig1 | Pi4k2b | Taf1b |  |
| Capn15 | Ffar2 | Lrig3 | Piezo1 | Tango6 |  |
| Capn5 | Fga | Lrmp | Pif1 | Tap1 |  |
| Capn9 | Fgb | Lrp10 | Pigb | Tap2 |  |
| Capns2 | Fgd6 | Lrp2 | Pigg | Tapbp |  |
| Car12 | Fgf20 | Lrp5 | Pign | Tapbpl |  |
| Car2 | Fgf21 | Lrp6 | Pigo | Tarm1 |  |
| Car5a | Fgg | Lrrc19 | Pigv | Tas1r2 |  |
| Car5b | Fgl1 | Lrrc25 | Pik3c2b | Tas1r3 |  |
| Car6 | Fgr | Lrrc27 | Pik3c2g | Tas2r105 |  |
| Carf | Fhad1 | Lrrc43 | Pik3c3 | Tas2r115 |  |
| Cars | Fkbpl | Lrrc6 | Pik3r2 | Tas2r117 |  |
| Casc1 | Flg | Lrrc66 | Pik3r4 | Tas2r129 |  |
| Casc5 | Flt1 | Lrrc9 | Pink1 | Tas2r130 |  |
| Casp8 | Flt3 | Lrrfip1 | Pira2 | Tas2r131 |  |
| Casp8ap2 | Flywch1 | Lta4h | Pirb | Tbc1d2 |  |
| Casp9 | Flywch2 | Ltb | Pkd1 | Tbc1d30 |  |
| Casz1 | Fmn2 | Ltbp3 | Pkd2 | Tbx18 |  |
| Catsper2 | Fmo4 | Ltbp4 | Pkd2l1 | Tbx3 |  |
| Catsper3 | Fmo6 | Ltk | Pkd2l2 | Tbx6 |  |
| Catsperg1 | Fndc7 | Luzp1 | Pkhd1 | Tc2n |  |
| Cbll1 | Focad | Ly6a | Pkmyt1 | Tceal6 |  |
| Cbs | Foxa1 | Ly6c1 | Pla1a | Tcf12 |  |
| Cbx7 | Foxb2 | Ly6c2 | Pla2g2c | Tcfl5 |  |
| Cc2d1a | Foxd4 | Ly6g6e | Pla2g2e | Tdg |  |
| Cc2d2a | Foxm1 | Ly6i | Pla2g4b | Tdh |  |
| Ccdc114 | Foxn2 | Ly9 | Pla2g4e | Tdo2 |  |
| Ccdc122 | Foxo6 | Lyl1 | Pla2g7 | Tdrd3 |  |
| Ccdc124 | Fpr3 | Lypd8 | Plbd1 | Tdrd5 |  |
| Ccdc154 | Fras1 | Lyplal1 | Plcb2 | Tdrd6 |  |
| Ccdc163 | Frmd4b | Lyrm5 | Plch2 | Tecta |  |
| Ccdc180 | Frmd5 | Lzts1 | Plcz1 | Tekt4 |  |
| Ccdc191 | Frmpd1 | Lzts3 | Plekha4 | Telo2 |  |
| Ccdc24 | Frrs1 | Madd | Plekha7 | Tenm4 |  |
| Ccdc28b | Fryl | Maf1 | Plekhb1 | Tex15 |  |
| Ccdc30 | Frzb | Mageb3 | Plekhg6 | Tex16 |  |
| Ccdc34 | Fsip1 | Magi3 | Plekhh1 | Tex26 |  |
| Ccdc38 | Fstl5 | Majin | Plekhn1 | Tex35 |  |
| Ccdc64b | Ftl1 | Malt1 | Plin2 | Tff1 |  |
| Ccdc68 | Ftmt | Mamld1 | Plk3 | Tfip11 |  |
| Ccdc73 | Fto | Mamstr | Plvap | Tg |  |
| Ccdc80 | Fubp1 | Man2b1 | Plxnb1 | Tgfbr3l |  |
| Ccdc91 | Fuk | Manba | Pms1 | Tgm3 |  |
| Ccdc92 | Fut10 | Mansc1 | Pnisr | Tgm5 |  |
| Ccdc92b | Fuz | Mansc4 | Pnkp | Tgm6 |  |
| Ccdc93 | Fxn | Map1a | Pnp | Tgm7 |  |
| Ccer2 | Gaa | Map1b | Pnp2 | Thap3 |  |
| Cchcr1 | Gabrr3 | Map2k6 | Pof1b | Thap4 |  |
| Ccm2l | Gadd45gip1 | Map3k1 | Pold2 | Thbs1 |  |
| Ccnf | Galnt10 | Map3k6 | Pole | Them7 |  |
| Ccni | Galnt9 | Map4k1 | Pole2 | Thoc5 |  |
| Ccsap | Ganc | Map6 | Poli | Tiam2 |  |
| Cd180 | Gars | Map7d1 | Poln | Ticrr |  |
| Cd2 | Gart | Map9 | Polr2i | Tie1 |  |
| Cd200 | Gas1 | Marc1 | Pom121l12 | Timm22 |  |
| Cd22 | Gast | March2 | Pomk | Tln1 |  |
| Cd244 | Gba2 | Marco | Pou6f1 | Tln2 |  |
| Cd300c2 | Gbp10 | Mark1 | Pp2d1 | Tlr11 |  |
| Cd300ld | Gbp6 | Masp2 | Ppargc1b | Tlr12 |  |
| Cd300ld4 | Gcat | Mast2 | Ppih | Tlr2 |  |
| Cd300lg | Gcdh | Mast4 | Ppip5k1 | Tlr4 |  |
| Cd38 | Gcfc2 | Matn2 | Ppm1m | Tlr5 |  |
| Cd40 | Gcm2 | Matn4 | Ppp1r14a | Tlr9 |  |
| Cd44 | Gcn1l1 | Mbd1 | Ppp1r14d | Tm4sf19 |  |
| Cd48 | Gdap2 | Mbd2 | Ppp1r18 | Tm6sf1 |  |
| Cd72 | Gdf1 | Mbd6 | Ppp1r21 | Tm6sf2 |  |
| Cd84 | Gdf15 | Mblac1 | Ppp1r3g | Tm7sf2 |  |
| Cdan1 | Gdf3 | Mboat4 | Ppp5c | Tm7sf3 |  |
| Cdc14b | Gdf5 | Mboat7 | Pprc1 | Tmc2 |  |
| Cdc25a | Gdi1 | Mbtd1 | Pqlc3 | Tmc4 |  |
| Cdc34b | Gdpd4 | Mc5r | Pram1 | Tmc5 |  |
| Cdc7 | Gdpgp1 | Mcfd2 | Prb1 | Tmco4 |  |
| Cdca7l | Gemin5 | Mcmdc2 | Prcc | Tmed5 |  |
| Cdh11 | Gen1 | Mcph1 | Prdm14 | Tmem104 |  |
| Cdh15 | Gfer | Mctp1 | Prdm15 | Tmem108 |  |
| Cdh24 | Gfpt1 | Mdm1 | Prdm16 | Tmem125 |  |
| Cdh6 | Gga3 | Mdm4 | Prdm6 | Tmem132b |  |
| Cdh7 | Ggct | Med13l | Prdx6 | Tmem151a |  |
| Cdhr3 | Ggh | Med8 | Prh1 | Tmem151b |  |
| Cdk20 | Ggn | Medag | Prkar2b | Tmem171 |  |
| Cdk5rap3 | Ggnbp1 | Megf6 | Prkca | Tmem181a |  |
| Cdkl2 | Gimap8 | Mei1 | Prkd3 | Tmem201 |  |
| Cdkn1b | Gip | Mepe | Prkdc | Tmem221 |  |
| Cdkn2a | Glb1 | Mertk | Prlhr | Tmem232 |  |
| Cdon | Glb1l3 | Methig1 | Prmt5 | Tmem237 |  |
| Cdpf1 | Gli1 | Mettl13 | Procr | Tmem241 |  |
| Cdsn | Gli2 | Mettl14 | Prodh2 | Tmem252 |  |
| Ceacam15 | Glod4 | Mettl17 | Prom1 | Tmem253 |  |
| Ceacam3 | Glp2r | Mettl7a2 | Proser2 | Tmem260 |  |
| Ceacam5 | Gls2 | Mettl7a3 | Prp2 | Tmem28 |  |
| Cebpe | Glt8d2 | Mex3c | Prpmp5 | Tmem33 |  |
| Cebpz | Glyctk | Mfap3 | Prr14l | Tmem35b |  |
| Cecr5 | Gm10032 | Mfsd14b | Prr18 | Tmem38a |  |
| Cela2a | Gm10306 | Mfsd7b | Prr30 | Tmem39b |  |
| Celsr1 | Gm10309 | Mgat4d | Prr36 | Tmem55b |  |
| Celsr3 | Gm10382 | Mgp | Prrc2a | Tmem59l |  |
| Cemip | Gm1043 | Micall1 | Prrc2c | Tmem62 |  |
| Cenpc1 | Gm1045 | Micall2 | Prss32 | Tmem70 |  |
| Cenpk | Gm10639 | Mif4gd | Prss41 | Tmem71 |  |
| Cenpl | Gm10750 | Mina | Prss50 | Tmem88b |  |
| Cep112 | Gm10775 | Mis18bp1 | Prss52 | Tmprss11g |  |
| Cep120 | Gm10912 | Mks1 | Prss55 | Tmprss13 |  |
| Cep131 | Gm11127 | Mlip | Prune2 | Tmprss2 |  |
| Cep170 | Gm1113 | Mlkl | Prx | Tmprss3 |  |
| Cep192 | Gm11559 | Mmp24 | Psd3 | Tmprss7 |  |
| Cep250 | Gm11563 | Mmp9 | Psg27 | Tmtc1 |  |
| Cep350 | Gm11568 | Mms19 | Psmd8 | Tmtc2 |  |
| Cep44 | Gm11595 | Mn1 | Psme4 | Tmub2 |  |
| Cep68 | Gm12169 | Mnda | Ptar1 | Tnfaip3 |  |
| Cep78 | Gm12253 | Mndal | Ptcd1 | Tnfrsf12a |  |
| Cep89 | Gm12258 | Mogat1 | Ptchd2 | Tnfrsf14 |  |
| Cers5 | Gm12888 | Mon1b | Ptchd3 | Tnfrsf19 |  |
| Ces1a | Gm13741 | Morc2b | Ptchd4 | Tnfrsf21 |  |
| Ces1c | Gm13757 | Morc3 | Ptdss1 | Tnfsf11 |  |
| Cfap43 | Gm14085 | Mpc1 | Ptgr1 | Tnk1 |  |
| Cfap46 | Gm14137 | Mpdz | Ptpn22 | Tnk2 |  |
| Cfap54 | Gm14548 | Mpeg1 | Ptprd | Tnks1bp1 |  |
| Cfap61 | Gm15292 | Mpg | Ptprf | Tnn |  |
| Cgnl1 | Gm15448 | Mphosph10 | Ptprh | Tnr |  |
| Chac1 | Gm1553 | Mpl | Ptprn2 | Tns3 |  |
| Chd1 | Gm156 | Mpo | Ptpro | Tnxb |  |
| Chd4 | Gm15698 | Mpp3 | Purb | Tob2 |  |
| Chd8 | Gm16181 | Mpp4 | Pwwp2b | Toe1 |  |
| Chd9 | Gm17078 | Mpp5 | Pxdn | Tom1l1 |  |
| Chdh | Gm17359 | Mptx1 | Pycr1 | Tonsl |  |
| Chek2 | Gm19410 | Mrap2 | Pydc3 | Topaz1 |  |
| Chm | Gm28042 | Mrgprd | Pydc4 | Topbp1 |  |
| Chn2 | Gm28372 | Mrgprf | Pyhin1 | Tor1aip1 |  |
| Chrna4 | Gm28557 | Mro | Pyroxd2 | Tor1aip2 |  |
| Chst14 | Gm29106 | Mroh7 | Pzp | Tor3a |  |
| Chst4 | Gm3336 | Mrpl22 | Qprt | Tox |  |
| Chst9 | Gm34302 | Mrpl48 | Qrich2 | Tox4 |  |
| Chtf18 | Gm382 | Mrpl49 | R3hcc1l | Tph2 |  |
